# Testing Adaptive Therapy Protocols using Gemcitabine and Capecitabine on a Mouse Model of Endocrine-Resistant Breast Cancer

**DOI:** 10.1101/2023.09.18.558136

**Authors:** Sareh Seyedi, Ruthanne Teo, Luke Foster, Daniel Saha, Lida Mina, Donald Northfelt, Karen S. Anderson, Darryl Shibata, Robert Gatenby, Luis Cisneros, Brigid Troan, Alexander R. A. Anderson, Carlo C. Maley

## Abstract

Highly effective cancer therapies often face limitations due to acquired resistance and toxicity. Adaptive therapy, an ecologically inspired approach, seeks to control therapeutic resistance and minimize toxicity by leveraging competitive interactions between drug-sensitive and drug-resistant subclones, prioritizing patient survival and quality of life over maximum cell kill. In preparation for a clinical trial in breast cancer, we used large populations of MCF7 cells to rapidly generate endocrine-resistance breast cancer cell line. We then mimicked second line therapy in ER+ breast cancers by treating the endocrine-resistant MCF7 cells in a mouse xenograft model to test adaptive therapy with capecitabine, gemcitabine, or the combination of those two drugs. Dose-modulation adaptive therapy with capecitabine alone increased survival time relative to MTD, but not statistically significant (HR: 0.22, 95% CI 0.043-1.1 P = 0.065). However, when we alternated the drugs in both dose modulation (HR = 0.11, 95% CI: 0.024 - 0.55, P = 0.007) and intermittent adaptive therapies significantly increased survival time compared to high dose combination therapy (HR = 0.07, 95% CI: 0.013 - 0.42; P = 0.003). Overall, survival time increased with reduced dose for both single drugs (P < 0.01) and combined drugs (P < 0.001). Adaptive therapy protocols resulted in tumors with lower proportions of proliferating cells (P = 0.0026) and more apoptotic cells (P = 0.045). The results show that Adaptive therapy outperforms high-dose therapy in controlling endocrine-resistant breast cancer, favoring slower-growing tumors, and showing promise in two-drug alternating regimens.

## INTRODUCTION

Therapeutic resistance and therapeutic toxicity are two of the biggest challenges in oncology. However, resistance often comes at a fitness cost for the resistant cells (*1,2*). Pest managers have long ago understood that pesticide resistance also comes at a fitness cost (*3*). Pests that are sensitive to the pesticide can out-compete pests that are resistant in the absence or at low doses of the pesticide. Pest managers learned to use the minimum effective dose (MED) of pesticides rather than the MTD (*3*). Current cancer treatment is aimed at killing as many tumor cells as possible in the shortest time, accomplished by utilizing drugs at the maximum tolerated dose (MTD). This approach is highly toxic and often results in a selective advantage for therapy-resistant cells which eventually kill the patient (Fig 1A) (*4*). Under the assumption that prior to treatment a tumor is mainly composed of drug sensitive cells with only a few pre-existing resistant cells, then treatment rapidly kills most sensitive cells, resulting in competitive release of the resistant cells. In fact, evolutionary theory predicts that the fastest way to select for therapeutic resistance is to use the MTD (*5*). Following this principle, we expect that in the absence of therapy, sensitive cancer cells will probably proliferate at the expense of the less-fit resistant cells. Thus, according to integrated pest management (IPM), a therapeutic strategy that is explicitly designed to maintain therapeutically sensitive organisms, could increase a patient’s survival by using sensitive cells to suppress the growth of resistant cells and thereby being the inspiration for adaptive therapy in cancer treatment (*4*).

**Fig. 1.**
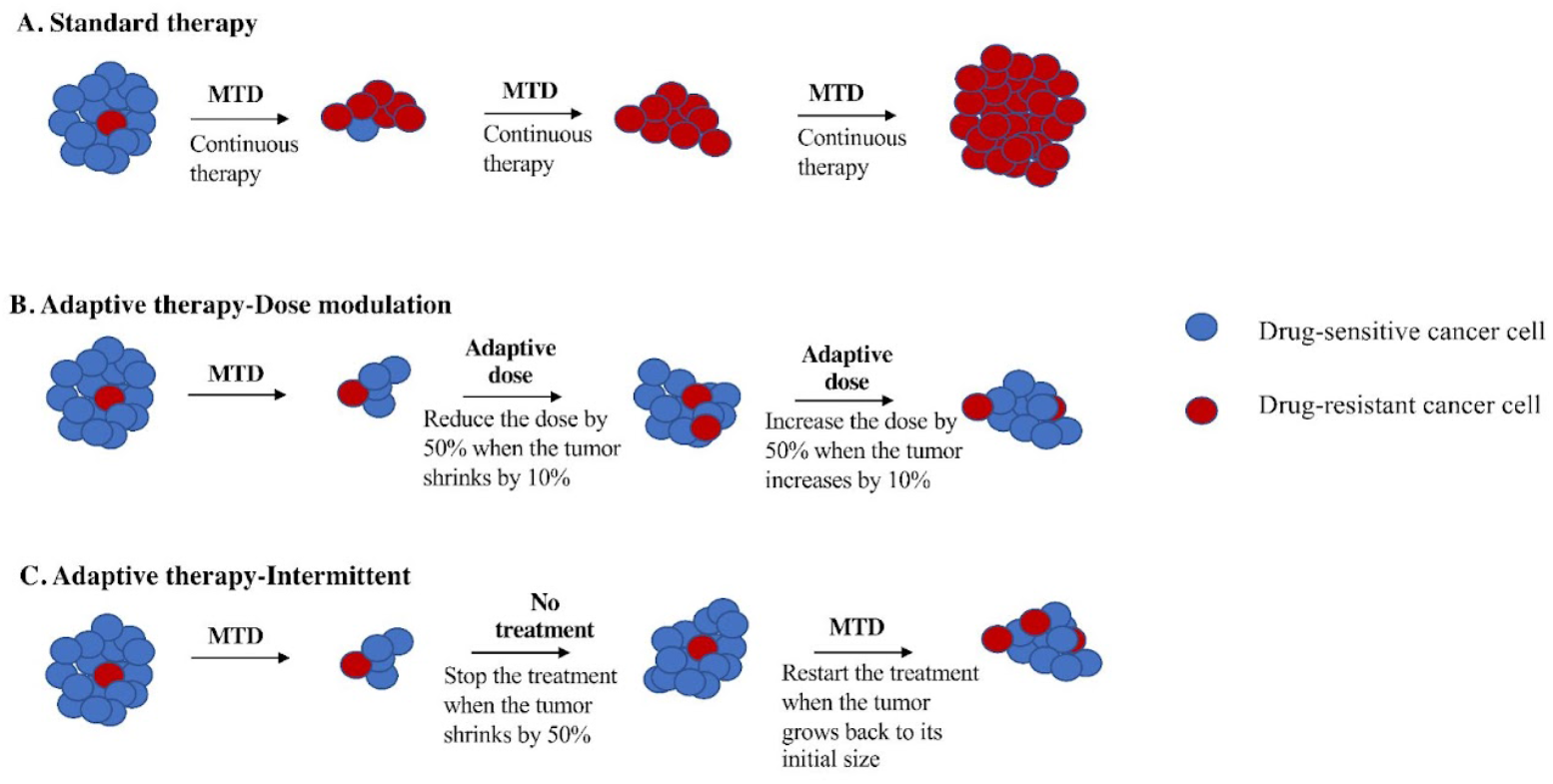
Schematic Figure of Comparison Adaptive Therapy Protocols with Standard Therapy. **(A)** Standard therapy selects for cells (red) that are resistant to treatment and tumor relapse. Adaptive therapy maintains a stable tumor volume by preserving drug-sensitive cells (blue) that suppress the growth of less fit, resistant cells (red). (**B)** Dose modulation adaptive therapy that raises the dose if the tumor grows and lowers the dose if the tumor shrinks. Previous mouse models used a tumor burden change of 20% to trigger a change in dose (*2*). Our simulation studies suggest a lower threshold is better (*17*), so here we have used a 10% change in tumor burden to trigger a change in dose. C. Intermittent adaptive therapy stops dosing altogether if the tumor burden falls below a threshold (e.g., 50% of its initial value), and restarts treatment if the tumor recovers (e.g., to 100% of its initial value) (*6,9*).

Adaptive therapy exploits intra-tumor cell competition to prolong patient survival, by allowing sensitive cells to compete with resistant cells during periods of drug holiday or low drug dose (*2,6*). Initial tests of adaptive therapy have been conducted with single drugs (*2,4,6*). Two protocols have been tested in pre-clinical models: dose modulation (a.k.a., AT1) (*2,4,6*) and dose skipping (a.k.a., AT2) (*2,4,6*). The dose modulation protocol, compares the current size of the tumor to the previous measurement then, raises the drug dose if the tumor has grown (>20%), lowers the dose if the tumor shrank (>20%), and keeps the dose the same if the tumor burden has stayed within 80–120% (Fig 1B). The dose skipping protocol uses the MTD dose but skips a dose if the tumor shrank or remained stable since the last measurement. Results from pre-clinical models have shown that adaptive therapy with dose modulation is able to achieve indefinite control in mouse xenografts of metastatic triple-negative (MDA-MD-231) and an estrogen receptor-positive (ER+) (MCF7) breast cancer (*2,4,6*), as well as in ovarian cancer (OVCAR3) xenografts (*4*). In contrast, the dose skipping (drug holiday) protocol was unable to prevent tumors from growing compared with standard of car (*2,4,6*). On/off regimens were generally ineffective for fast-growing tumors due to a loss of control during the off cycle, but they found applicaion this method in the context of slow-growing tumors, leading to their implementation in a clinical trial for prostate cancer (*2,7,8*).

The first clinical trial of single-drug adaptive therapy used a different protocol on metastatic, castration-resistant prostate cancer. This intermittent adaptive therapy protocol (Fig 1C) used MTD of single agent abiraterone until the tumor burden (measured in blood by prostate specific antigen [PSA]) fell below 50% of its initial level. At that point, treatment stopped and was only resumed if the PSA level returned to its initial level. The administration of abiraterone is determined using mathematical models informed by evolutionary principles in order to prolong the emergence of resistance. This resulted in a doubling of time to progression, using less than half of the accumulative drug dose compared to standard of care (*6,8,9*).

Our motivating goal is to test if adaptive therapy can improve clinical outcomes such as time to progression, overall survival and reductions in toxicity in breast cancer, but more specifically to test multidrug adaptive therapies as well as single drug therapy on endocrine-resistant breast cancer. Although a variety of adaptive therapy protocols have been tested in mice (*2,4,6,10*) and computational simulations (*11–13*), it remains an open question which version of adaptive therapy is best, and under which conditions one protocol might be better than another (*14*). Here we test different adaptive therapy protocols, and compare them to the time to progression (TTP) of MTD in a preclinical model of breast cancer. We focused on endocrine-resistant ER+ breast cancer, an ongoing major clinical challenge. We conducted a preclinical experiment to study the effect of gemcitabine and capecitabine on a xenograft model of endocrine-resistant ER+ cells in NOD SCID gamma (NSG) mice. We compared gemcitabine to capecitabine to help inform the choice of drug for a future clinical trial. Furthermore, because we were interested in how multiple drugs might be combined in adaptive therapy, based on previous simulation results (*14*), we tested four different multidrug adaptive therapy protocols. Although the combination of gemcitabine and capecitabine is not theoretically justified, since they have the similar mechanisms of action, there are some studies that suggest little cross-resistance between gemcitabine and capecitabine (*15,16*) (Supplementary Methods, Data S1). We capitalized on the opportunity to test multidrug adaptive therapies for the first time, while also addressing the primary goal with the single drug adaptive therapy. These studies provide a framework for the rational design of adaptive combination chemotherapy protocols to control therapeutic resistance in breast cancer.

## RESULTS

We tested whether adaptive therapy works in preclinical mouse models with endocrine-resistant breast cancer and compared adaptive therapy with standard therapy. We tested the two leading protocols for adaptive therapy (Dose modulation vs. Intermittent) in mice, using gemcitabine and/or capecitabine on the human breast cancer cell line MCF7 that evolved to be resistant to fulvestrant and palbociclib.

### Endocrine-Resistant MCF7 Cell Line

Typically, it takes 6-18 months to select for therapeutically resistant cells in a cell line (*24*). However, resistance often emerges in the clinic in a matter of a few months (*25–27*). We hypothesized that this discrepancy may be due to the use of unrealistically small cell population sizes *in vitro*. Population size is a key parameter of the rate of evolution (*28*). By using a hyperflask to culture a population of ∼10^8^ MCF7 cells, we were able to generate cells that were resistant to 5-fold higher doses of fulvestrant and palbociclib in one month (Fig 2). After a month of exposure to both fulvestrant and palbociclib, MCF7 resistant (MCF7 R) cells had evolved a half-maximal concentration inhibitory concentration (IC50) value of 62.65 uM compared to 13.13 uM for the sensitive parental cells (MCF7 S).

**Fig. 2.**
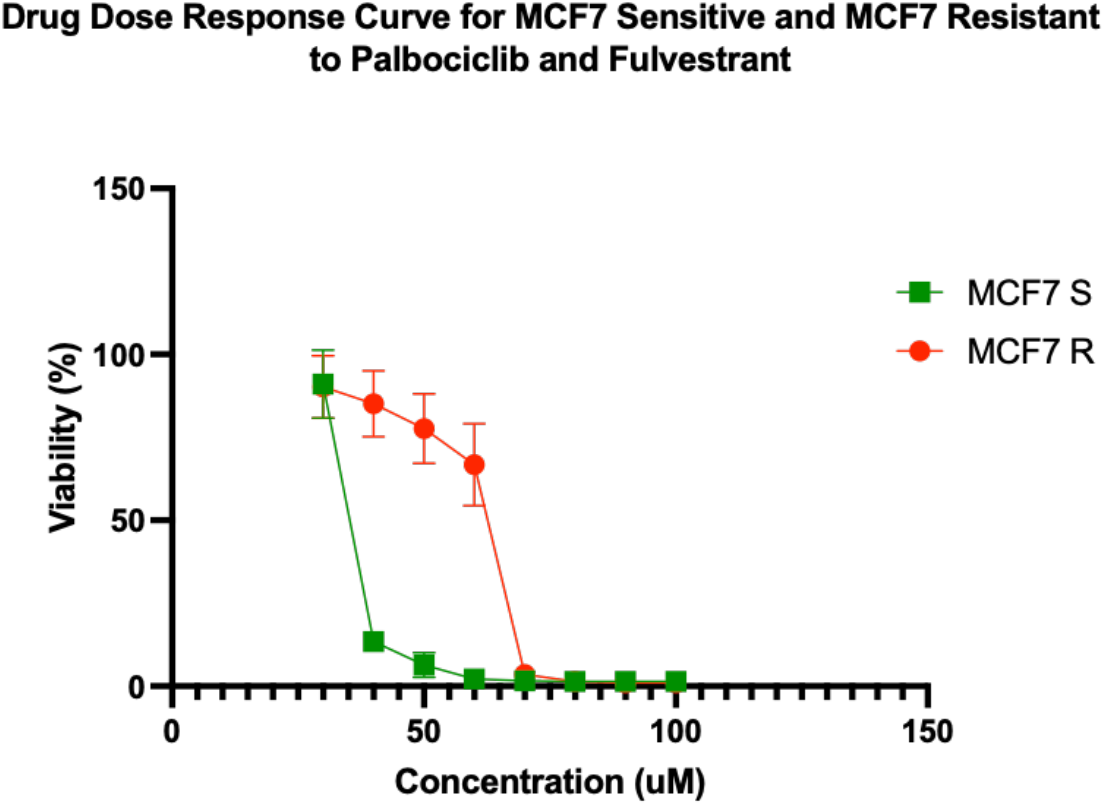
Drug dose-response curves for sensitive and resistant MCF7/luc cell lines. MCF7 resistant (MCF7 R) and sensitive (MCF7 S) cell lines showed an IC50 value of 62.65 uM and 13.13 uM respectively. The percentage of viability at 40 uM, 50 uM, and 60 uM were significantly different between MCF7 R and MCF7 S cell lines. Each data point represents 8 replicates.

### Tumor Growth Control After Therapy Cessation

In 20 of the 48 mice on adaptive therapy protocols, the tumor burden shrank below our minimum threshold at which we stopped therapy, and then remained at low tumor burden for many weeks until the mice died or had to be sacrificed for other reasons (Table S1). In an additional 7 mice, the tumor stayed stable after the cessation of therapy, but eventually did regrow. It is not clear why the tumor cells, still detectable by bioluminescence, did not grow in the absence of drug or a functional immune system.

### Prolonged Survival Benefit of Adaptive Therapy

#### Single-drug therapy

In capecitabine, single-drug adaptive therapy (Fig 3), survival was better, though not statistically significant, for dose modulation compared to no treatment (Cox proportional hazards HR = 0.24, 95% CI 0.042- 1.4, *P =* 0.1) and compared to MTD (HR = 0.22, 95% CI 0.043- 1.1, *P* = 0.065). There was some evidence that dose modulation prolonged survival compared to intermittent therapy (Fig 3, HR = 0.26, 95% CI 0.049- 1.4, *P =* 0.09). There were no differences between intermittent therapy relative to no treatment (HR = 0.96, 95% CI 0.247- 3.7, *P =* 0.95) and MTD (HR: 0.78, 95% CI 0.236- 2.6, *P =* 0.68). There was no significant benefit in prolonging survival between MTD therapy relative to no treatment (HR= 1.2, 95% CI 0.323- 4.5, *P =* 0.78) (Fig 3A). In addition, a lower average total drug dose was used in dose modulation (533.8 mg/kg) compared to intermittent (1226.6 mg/kg) and MTD therapies (1373.3 mg/kg) (Table S2). In gemcitabine single-drug adaptive therapy, there were no statistically significant differences in survival time for any of the protocols (Fig 3B). However, we found that a lower average total drug dose was used in dose modulation (256.9 mg/kg) compared to intermittent (333.3 mg/kg) and MTD (783.3 mg/kg) therapies (Table S2).

**Fig. 3.**
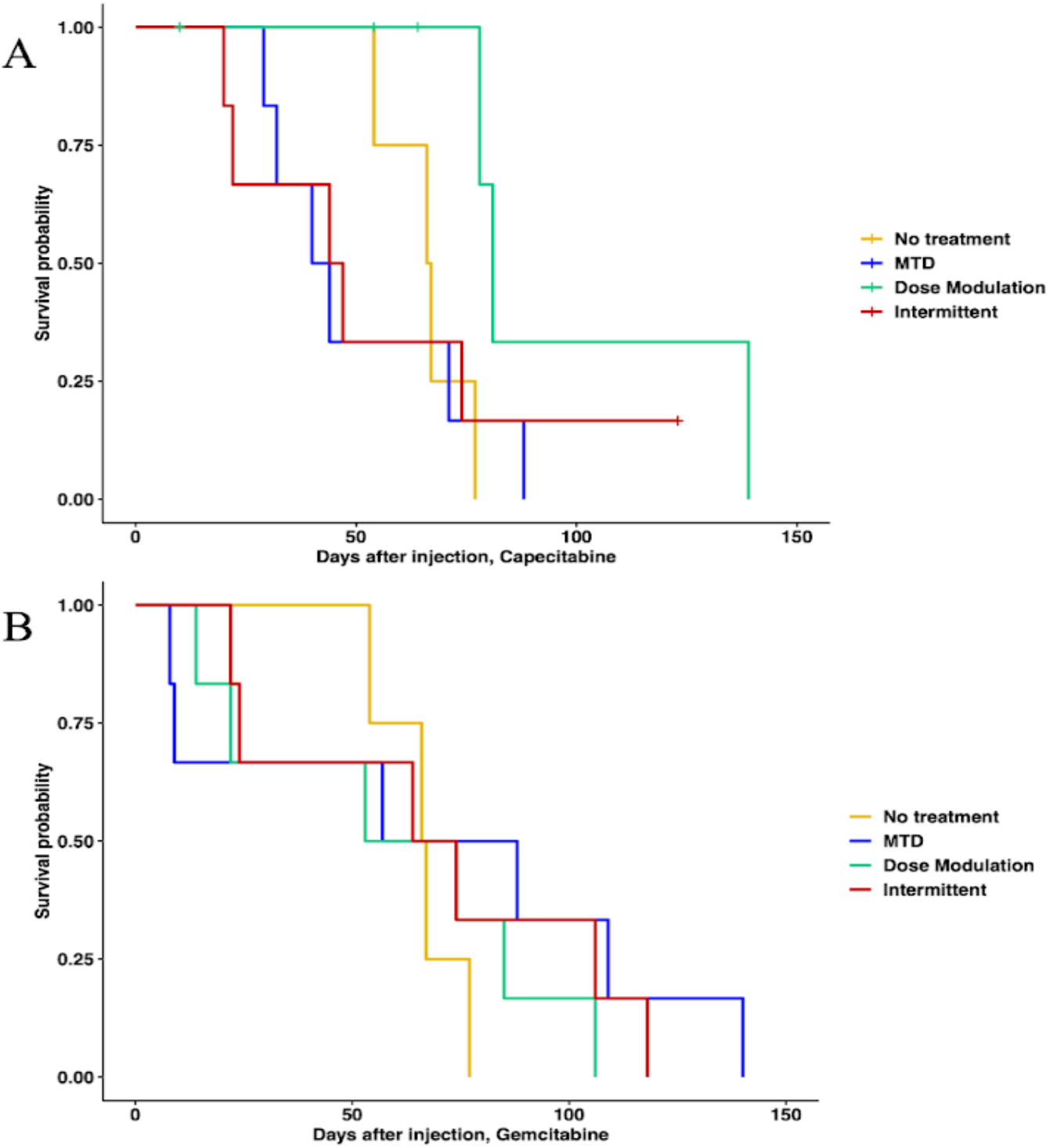
Survival analysis of Single-drug therapies. **(A)** Survival analysis of capecitabine protocols. Cox Regression (CAP Dose modulation relative to no treatment HR: 0.24, 95% CI 0.042- 1.4, *P =* 0.1; CAP Intermittent relative to no treatment HR: 0.96, 95% CI 0.247- 3.7, *P =* 0.95; CAP MTD relative to no treatment HR: 1.2, 95% CI 0.323- 4.5, *P =* 0.78). Cox Regression (CAP Dose modulation relative to MTD HR: 0.22, 95% CI 0.043- 1.1 *P* = 0.065; CAP Intermittent relative to MTD HR: 0.78, 95% CI 0.236- 2.6, *P =* 0.68). (**B)** Survival analysis of gemcitabine protocols. None of the protocols are statistically significantly different using Cox regressions.

#### Multi-drug Therapy

When using both gemcitabine and capecitabine, survival was better for ping-pong intermittent treatment relative to no treatment (HR = 0.13, 0.13, 95% CI: 0.02- 0.84, *P =* 0.032). There was evidence that survival was also better for ping-pong dose modulation compared to no treatment, but it was not quite statistically significant (Cox proportional hazards HR = 0.2, 95% CI: 0.038 - 1.09, *P =* 0.06). In comparison to MTD, both ping-pong dose modulation and ping-pong intermittent protocols prolonged survival (dose modulation HR = 0.11, 95% CI: 0.024 - 0.55, *P =* 0.007; intermittent therapy HR = 0.07, 95% CI: 0.013 - 0.42; *P* = 0.003). Survival under MTD with both drugs was worse than no treatment, but not significantly (HR = 1.9, 95% CI: 0.48 - 7.4; *P* = 0.36) (Fig 4A). Moreover, the average amount of drugs used per day was lower for both AT therapies (GEM: 4.9 mg/kg - CAP: 6.1 mg/kg per day for ping-pong dose modulation and GEM: 6.4 mg/kg - CAP: 8 mg/kg per day for ping-pong intermittent) compared to MTD (GEM:20.6 mg/kg - CAP: 30.6 mg/kg per day) (Table S2).

**Fig. 4.**
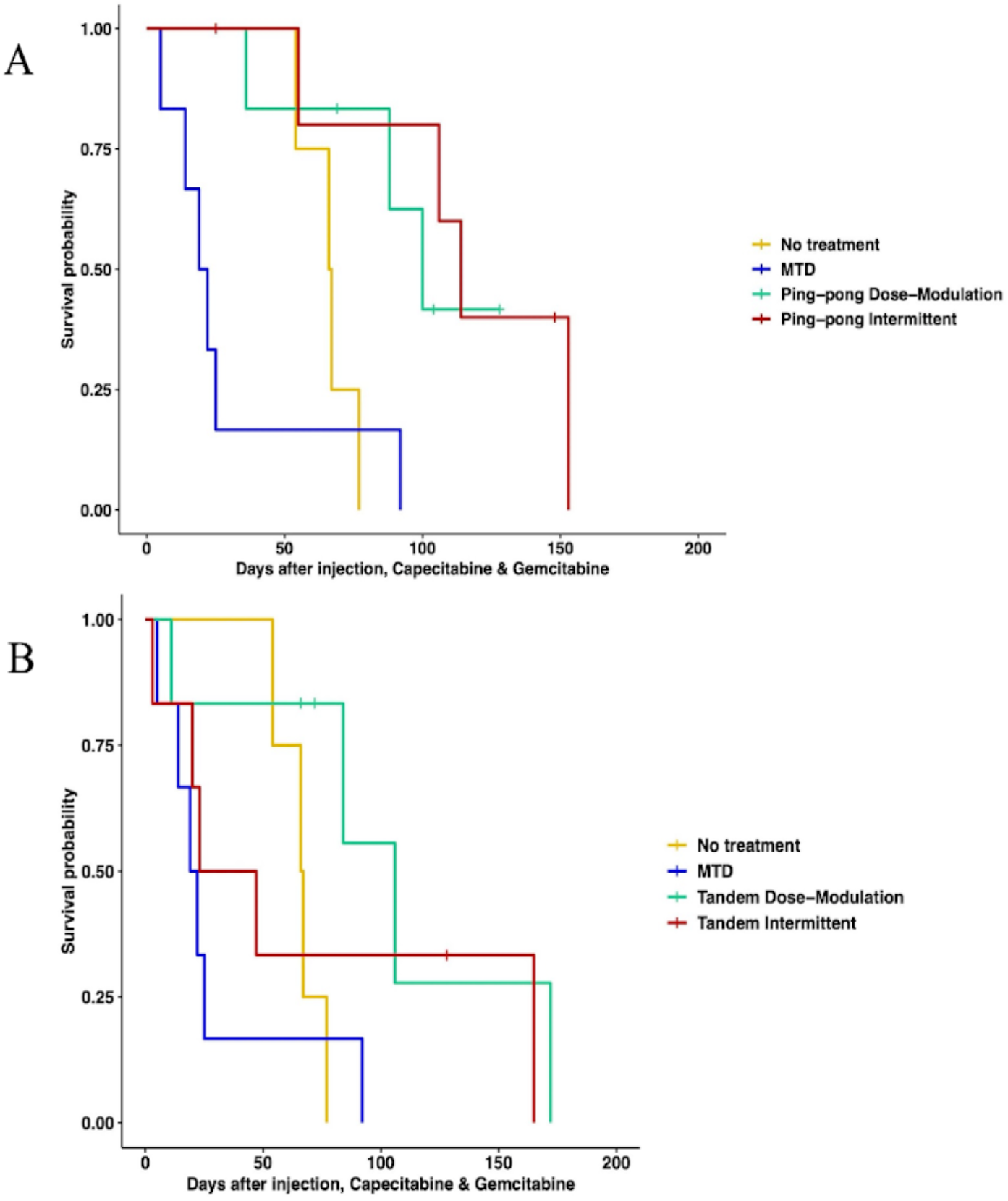
Survival analysis of multi-drug therapies. **(A)** Multi-drug ping-pong adaptive therapy protocols versus MTD or no treatment (vehicle control). In ping-pong protocols, only one drug is used at a time. Here MTD is the application of MTD of both gemcitabine and capecitabine using the same dose scheduling as single drug MTD. (**B**) Multi-drug tandem adaptive therapy protocols versus MTD or no treatment (vehicle control). In the tandem and MTD protocols, both gemcitabine and capecitabine are given (and modulated) at the same time.

With tandem combination adaptive therapy, where both drugs are used at the same time, dose modulation and intermittent protocols were better than combined MTD therapy, but the improvement was only statistically significant for dose-modulation (Cox proportional hazards for tandem dose-modulation vs. MTD HR = 0.22, 95% CI: 0.05 - 0.94, *P* = 0.04; for intermittent vs. MTD HR: 0.34, 95% CI: 0.07 - 1.6, *P* = 0.17). Neither form of tandem adaptive therapy was significantly better than no treatment (tandem dose modulation relative to no treatment HR: 0.34, 95% CI: 0.07 - 1.6, *P* = 0.17; Tandem intermittent relative to no treatment HR: 0.73, 95% CI: 0.17 - 3, *P* = 0.66). MTD was worse than no treatment in survival but the difference was not statistically significant (HR = 1.9, 95% CI: 0.48 - 7.4; *P* = 0.36) (Fig 4B).

### A Strong Correlation Between the Percentage of Maximum Tolerated Drug Dose Used with Survival Time

We calculated the cumulative total drug dose administered and divided it by the number of days between the start of therapy and death to get the average drug dose per day, normalized by the MTD. We found a strong negative correlation between the %MTD drug dose per day and the survival time of mice in combination therapy (*P*-value <0.0001, Fig 5A). We also observed a significant negative correlation between the %MTD drug dose per day and the survival time of mice in single drug therapies (capecitabine or gemcitabine) (*P*-value = 0.0074, Fig 5B). There was no relationship between the final tumor burden size and the survival time (Fig S1), though there was a trend such that higher concentrations of drug resulted in larger tumors at the end of the experiment (*P*=0.076, Fig S2).

**Fig. 5.**
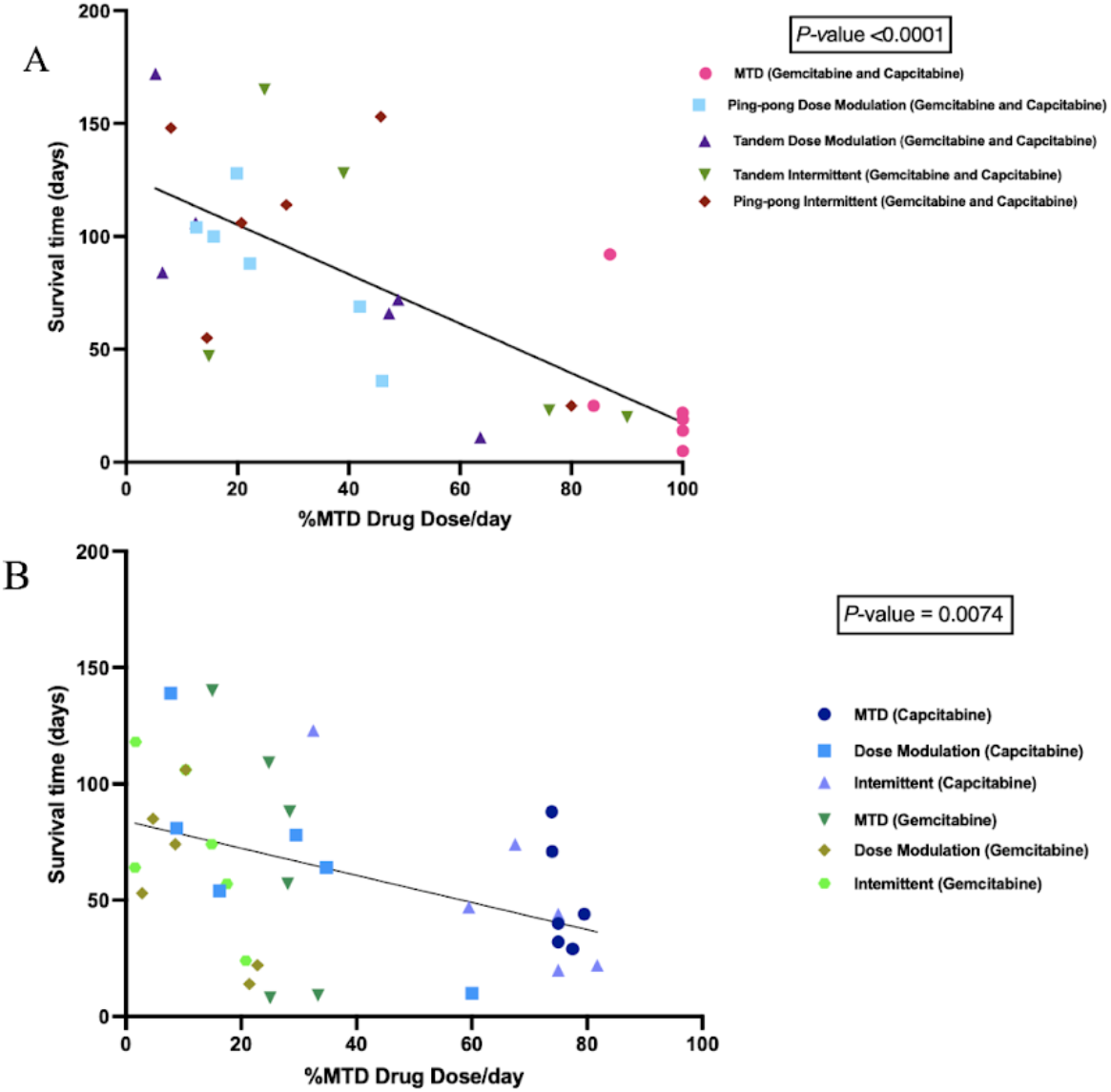
Correlation Between the Percentage of Maximum Tolerated Drug Dose Used with Survival Time. (**A)** Survival time as a function of the amount of combined gemcitabine and capecitabine that was given per day. Mice that were given more chemotherapy tended to have a shorter survival time (P-value <0.0001, R^2^ = 0.56). (**B)** Survival time as a function of the amount of single drug (either gemcitabine or capecitabine) that was given per day. Although there is a lot of variances associated with different protocols, there was still a strong trend that mice treated with more drug had a shorter survival time (P-value = 0.0074, R^2^ = 0.19).

### Different Chemosensitivity in Cell Lines Retrieved from Different Treatment Groups

To determine the chemosensitivity of retrieved cell lines from the tumors, we generated drug dose response (DDR) curves for single drug therapies and combination therapies (Fig S3). We generated DDR curves on parental pre-engrafted MCF7 breast cancer (ER+) cell lines with single and combination treatment of capecitabine and gemcitabine. Then we compared the sensitivity of engrafted cell lines under different treatment strategies for each drug to pre-engrafted MCF7 cell lines treated with that drug. In single-drug therapy using gemcitabine and combination therapy, we found cell lines from MTD groups showed the highest IC50s values, with the apparent exception of intermittent therapy with capecitabine (Fig 6). However, the two cell lines from tumors that were treated with the capecitabine intermittent protocol both came from tumors that never fell below 50% of their initial tumor burden, and so never had a drug holiday. Thus, they were treated the same as the MTD condition, which may explain why they resulted in relatively high IC50 values.

**Fig. 6.**
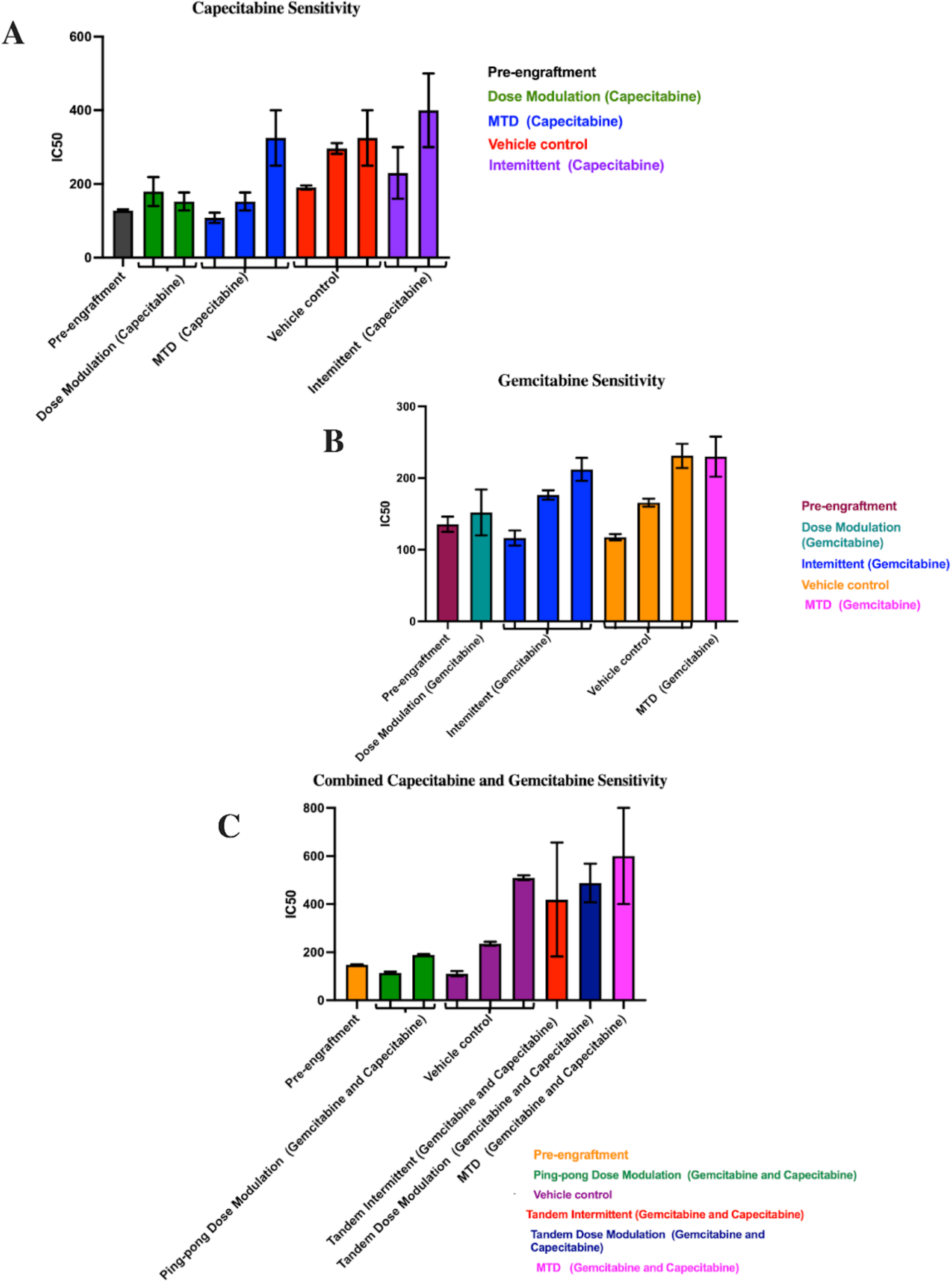
IC50 values for cell lines derived from tumors under different treatment conditions. In each panel, the protocols have been ordered from lowest to highest mean IC50 values (bars indicate means). (**A)** Capecitabine IC50 values for mice treated with capecitabine alone. (**B)** Gemcitabine IC50 values for mice treated with gemcitabine alone. (**C)** Combined capecitabine and gemcitabine IC50 values for mice treated with both drugs together. Error bars show the 95% confidence intervals on the IC50 values.

In capecitabine single therapy, we were able to retrieve cell lines from 3 mice of the MTD group, 2 mice in the dose modulation group, 2 mice in the intermittent group, and 3 mice that were not treated. Figure 6A shows the capecitabine IC50 values for those cell lines (Drug dose response curves are shown in Fig S3).

Gemcitabine IC50 values for mice treated with gemcitabine alone are shown in Fig 6B (Drug dose response curves are shown in Fig S3). For mice treated with the combination of both drugs, there were more dramatic differences in the IC50 values between conditions (Fig 6C). For the two cell lines from Mice 55 and 58 treated with the ping-pong dose modulation protocol and the one cell line from Mouse 50 treated with the intermittent tandem protocol, the cells were statistically significantly more sensitive to combination therapy compared to the cell line from Mouse 38 treated at MTD with both drugs (T-test *P*<0.001 at 100 uM, 150 uM and 250 uM, comparing Mice 55 and 58 vs. 38; for Mouse 50 vs. 38 *P* = 0.0026 at 100uM, and *P*< 0.0001 at both 150 uM and 250 uM; Supplemental DDR curves Fig S3).We were unable to derive a cell line from mice treated with the ping-pong intermittent protocol, and so they are not included in Fig 6C.

### Correlation Between IC50 Values with both Tumor Burden and Drug Dose

We hypothesized that a growing tumor at the time of death might indicate therapeutic resistance. We analyzed the relationship between the IC50 values of cell lines retrieved from the mice and the last measured tumor burden in those mice, based on bioluminescence. We found a significant positive correlation (*P*-value = 0.0041, R^2^ = 0.43; Fig 7A). Moreover, there was a positive correlation between the average drug dose applied per day for each mouse and the resulting IC50 values of the cells derived from those tumors (*P*-value = 0.04, R^2^ = 0.25; Fig 11), but this result is not statistically significant after correcting for multiple testing.

**Fig. 7.**
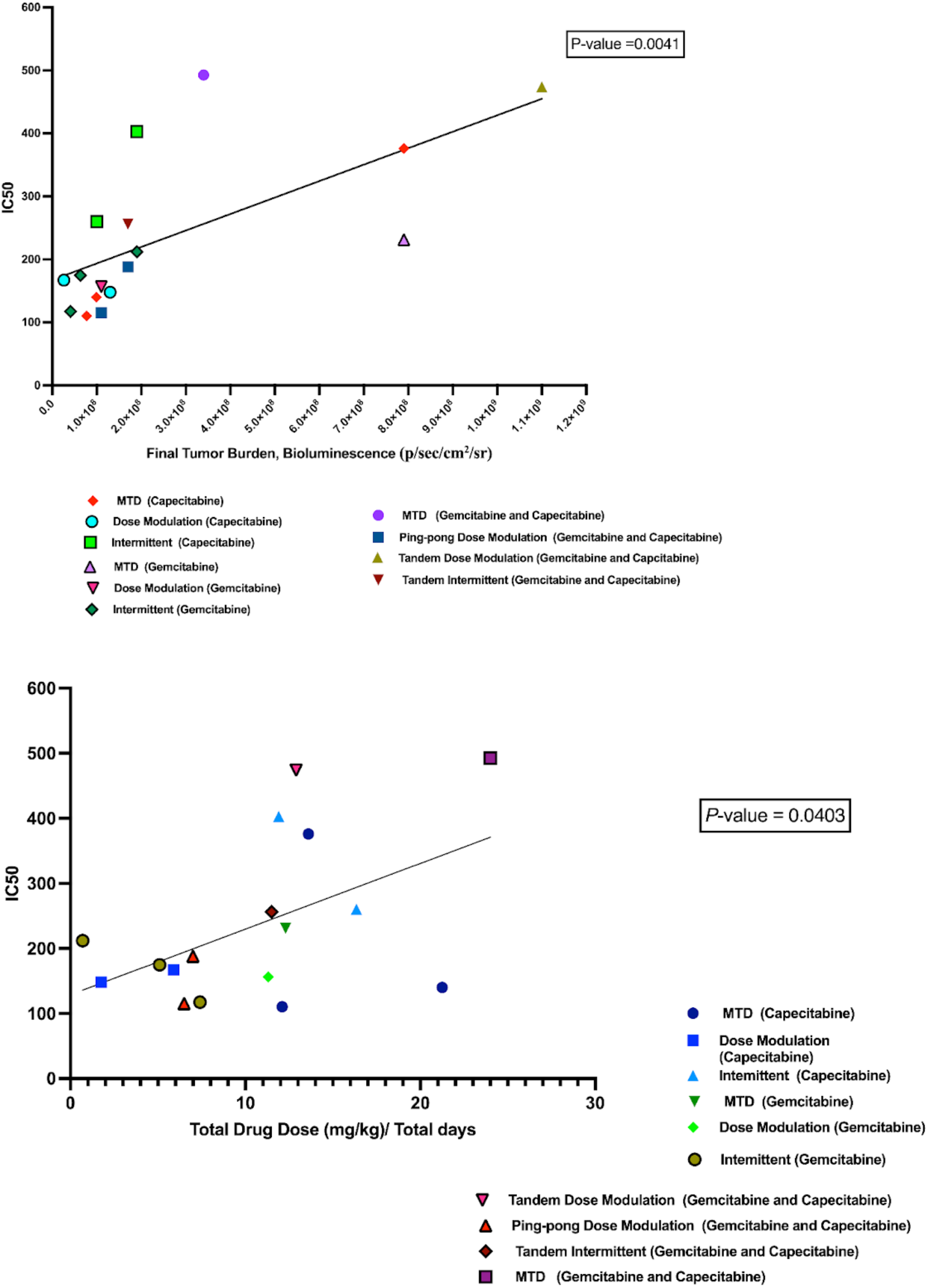
Correlation Between IC50 Values with both Tumor Burden and Drug Dose. **(A)** The relationship between the tumor burden at death and the IC50 values of the cell lines derived from those tumors. (**B)** The relationship between the average amount of drug used per day to treat a mouse and the resulting IC50 of the cells derived from that mouse’s tumor.

### Immunohistochemistry and Histological Analysis

We analyzed the percentage of apoptotic cells (Caspace-3 positive) and percentage of proliferating tumor cells (Ki-67 positive) out of the total number of cells examined (a standard field of view was utilized to compare the proportion of positive cells) on our tumor sections for all the tumors (Table S3). The maximum tolerated dose of both capecitabine and gemcitabine in single and combination strategies led to higher expression of Ki-67 than adaptive therapy protocols (t-test *P* = 0.0026; Fig 8A). In contrast, the maximum tolerated dose of both capecitabine and gemcitabine in single and combination strategies had lower caspase-3 index (lower apoptosis) compared to all the adaptive therapy groups (t-test *P* = 0.0457; Fig 8B), though this was not statistically significant after multiple testing correction. There was no evidence of a correlation between proliferation and apoptotic indices in the tumors (Fig S5).

**Fig. 8.**
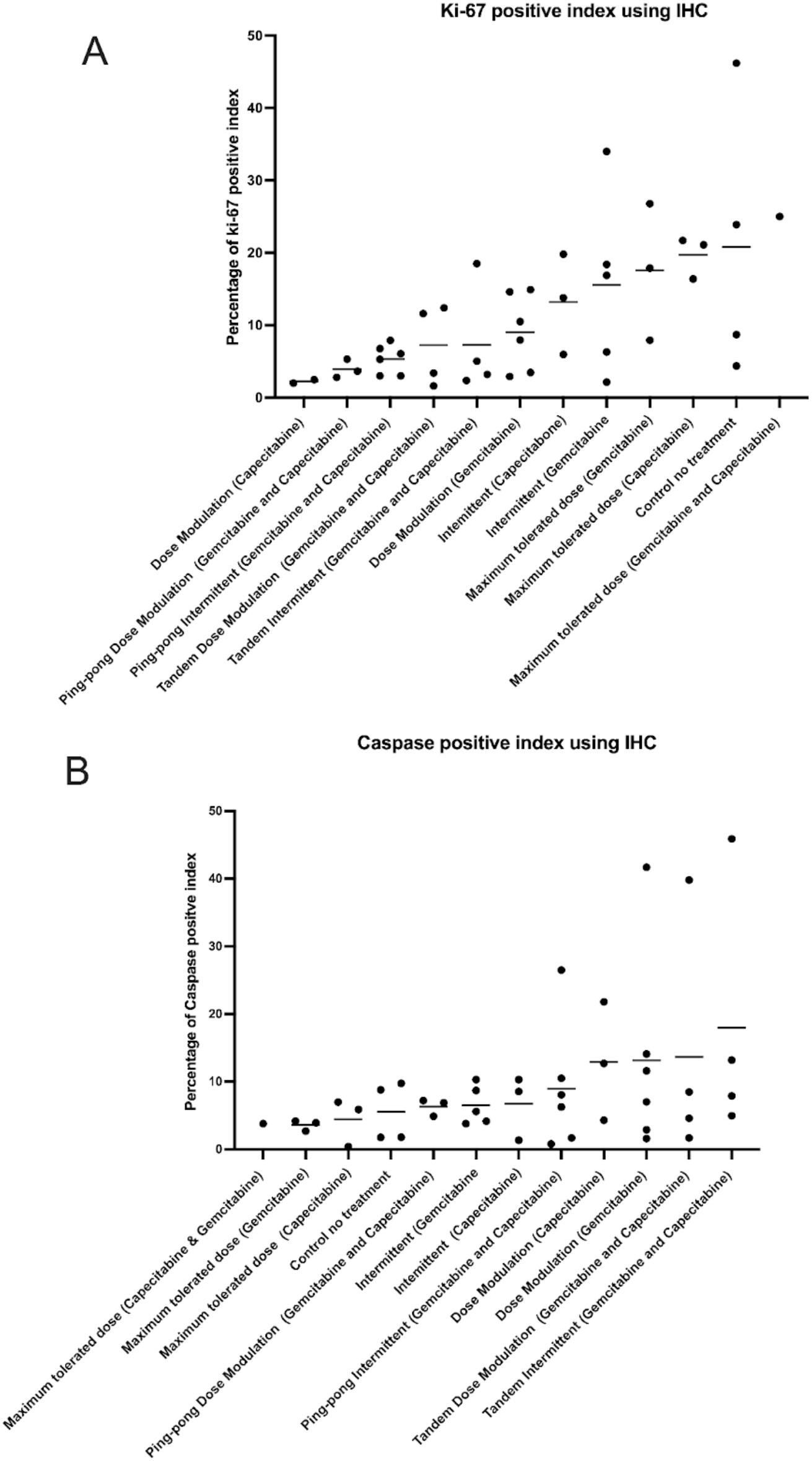
Immunohistochemistry Analysis. **(A)** The percentage of proliferating (Ki-67 positive) cells and **(B)** apoptotic (caspase-3 positive) cells in the tumors at the end of the different treatment protocols. Protocols have been ordered by increasing mean values (shown by the bars).

### Computational Simulations Match the Rank Orders for the Different Adaptive Therapy Protocols

We previously developed a hybrid agent-based model to explore adaptive therapy protocols using one (*29*) or two cytotoxic drugs (*17*). Although our model is simple and does not represent the details of breast cancer, gemcitabine or capecitabine, the results of the simulations and mouse experiments are consistent with respect to the rank order of their success. In both the modeling (Fig 9A) and experiments (Fig 9B) with capecitabine alone, the dose-modulation protocol works best, and the intermittent protocol is essentially equivalent to MTD. For the two-drug protocols, the dose-modulation ping-pong protocol, the dose-modulation tandem protocol and the ping-pong intermittent protocol are all equivalent and better than the other protocols in both the simulation and experimental results (Fig 9). The standard MTD protocol and the tandem intermittent protocols are equivalent, and worse than the other protocols in the mouse experiment, though in 3 of the 6 mice in the tandem intermittent group, the tumor never shrank below the threshold to stop dosing at MTD, and so they actually received the same treatment as the MTD group (Table S1). In the simulations, the tandem intermittent protocol does result in treatment holidays and it does better than MTD, but still not as well as the dose-modulation and ping-pong protocols (Fig 9C).

**Fig. 9.**
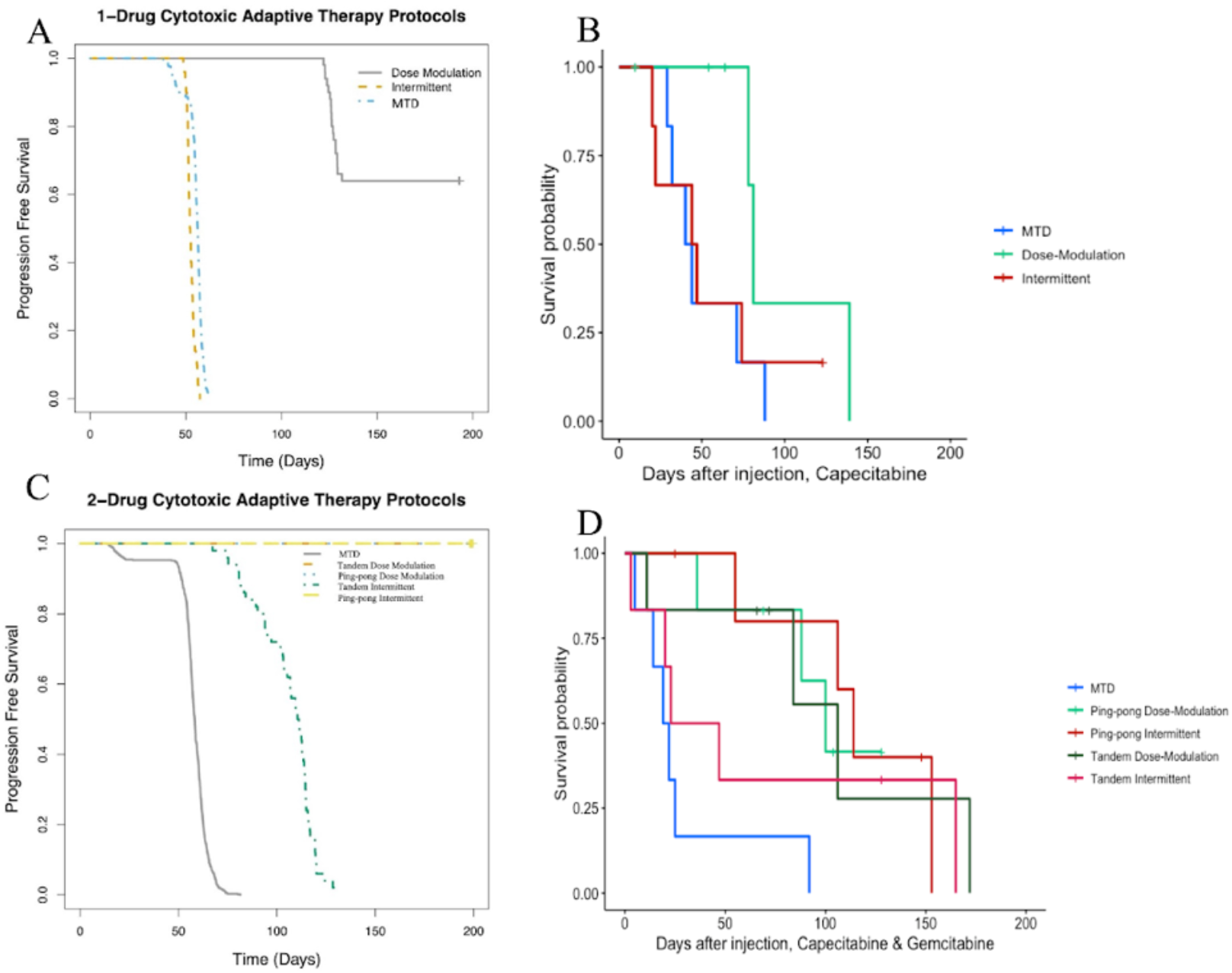
Computational Simulations Match the Rank Orders for the Different Adaptive Therapy Protocols. Comparison of **(A)** simulation results to **(B)** mouse experimental results for adaptive therapy and MTD protocols using capecitabine alone. **(C)** Comparison of simulation results to D. mouse experimental results for adaptive therapy and MTD protocols using both capecitabine and gemcitabine.

## DISCUSSION

Cancers are highly dynamic and cancer cells evolve phenotypic strategies to deal with the challenges of therapy. This suggests that our treatments should also be dynamic and adjust to how the cancer is changing (*2,4,6*). Standard therapy using MTD of cancer drugs imposes the strongest selective pressure to select for resistant phenotypes (*30*). Typically, that eliminates sensitive populations, resulting in the competitive release of resistant cells, and ultimately treatment failure (*5*). Adaptive therapy was translated from integrated pest management as a strategy for preventing therapeutically resistant clones from growing out of control, thereby improving patient survival, with the added benefit of reducing toxicity.

The innovation of this experiment was to test adaptive therapy with multiple drugs, testing ping-pong and intermittent protocols for the first time in mice. It is also the first time that adaptive therapy has been tested on an ER+ but endocrine-resistant breast cancer cell line. One advantage of the ping-pong protocols is that only one drug is applied at a time, which should both reduce toxicity and limit selection for multidrug resistance. Those strategies work best in both our simulations (*17*) and experiments. In general, dose modulation adaptive therapies work better than fixed (MTD) doses, probably because reducing the dose both reduces toxicity and selection for resistance (Figs 7, 8 and 11). Furthermore, in the intermittent protocol, we only stop dosing if the tumor burden falls below 50% of its initial value. This means that if the tumor never fell below that threshold, we kept dosing at MTD. In those cases, there was no difference between the intermittent and MTD protocols.

In many cases, adaptive therapy was able to maintain control over tumor growth with low doses. In 27 of the mice treated with adaptive therapy, the tumor burden fell below 2.25 × 10^8^ (photons/sec/cm2/sr) or 150 mm3 which triggered a cessation of treatment. In 20 of those 27, the tumor continued to shrink or stayed stable after therapy was stopped (Table S.1). This was a surprise. It may be due to some form of an Allee effect in which small, fragmented cancer cell populations have difficulty regrowing (*31–33*). Alternatively, it may be the result of neutrophils helping to control the tumors, since NSG mice still have functional neutrophils (*34*). But the mechanism of that effect has yet to be determined. We should also note that in 3 of the 4 control mice that were only treated with saline, the tumor stayed relatively stable over time (Table S.1).

By deriving cell lines from many of the tumors at the end of the experiment, we were able to test levels of therapeutic resistance that had evolved under the different protocols. We found that tumors exposed to long term MTD evolved more resistance than tumors treated with adaptive therapies. The resistance that evolved was correlated with the average dose applied per day.

Evolutionary life history theory suggests that there is likely to be a tradeoff between maximizing survival versus maximizing proliferation in cancer cells (*35*). Populations in stable environments tend to evolve slow life histories because they expand until resources are scarce and then have to compete for those limited resources (*36*). We had hypothesized that adaptive therapy, by attempting to maintain a stable tumor size, might select for cancer cells with a slower life history strategy than MTD. There are two counter-arguments to this hypothesis. The fact that we achieve that stable population size through external mortality (cytotoxic drugs), rather than resource limitations, might select for fast life history strategies. Furthermore, constant high dosing with cytotoxins might select for forms of resistance that invest heavily in survival over reproduction. In fact, adaptive therapy is based on the assumption that resistance has a fitness cost in the absence of drugs. However, when therapy is applied at high dose with recovery periods between doses, there may be selection for clones that can withstand therapy and then rapidly proliferate in the periods between therapy. Our results show that tumors that evolved under constant high dose treatment had high levels of proliferation and low levels of apoptosis, despite being exposed to cytotoxic drugs. In contrast, tumors that evolved under adaptive therapies had low levels of proliferation and higher levels of apoptosis, which should lead to a slower net proliferation rate compared to MTD (Figs 12 & 13). Thus, it appears that adaptive therapy protocols selected for slower life history strategies.

We discovered a simple way to increase the speed with which we can evolve resistant cell lines *in vitro*, based on evolutionary theory. Theory suggests that the rate of evolution is determined by five parameters: population size, generation time, mutation rate, selective coefficients and the heritability of those selected phenotypes. We hypothesized that the reason that acquired therapeutic resistance often evolves in the clinic in a matter of a few months, while most *in vitro* protocols require 6-18 months, often with slowly escalating doses, is due to a difference in the population sizes. Clinical tumors are estimated to have 10^8^ cells per cm^3^ (*37*), whereas typical cell culture conditions often have only 10^5^ cells. We treated 10^8^ cells in a hyperflask at IC70 of both fulvestrant and palbociclib, and were able to evolve a resistant variant of MCF7 cells in just one month.

This study had some caveats that could be addressed in follow-up investigations. First, with only 6 mice per group, our statistical power was limited for detecting survival differences between the protocols. We used two drugs with similar, though not identical mechanisms of action. Ideally, drugs that select for very different mechanisms of resistance should be used, which suggests we should choose drugs with very different mechanisms of action. It is reassuring that there may be little cross resistance between gemcitabine and capecitabine (*15,16*). Much of the advantage of adaptive therapies with multiple drugs that we observed was likely due to the comparison with our highly toxic constant “MTD” dose control condition. The fact that our multidrug “MTD” condition was worse than no treatment suggests that the doses we used for the combined gemcitabine and capecitabine caused a lot of toxicity and were actually above the maximum tolerable dose.

## CONCLUSION

Adaptive therapies show promise in this ER+ endocrine-resistant model of breast cancer. We found mice under adaptive therapy treatments not only had better survival but their tumors also evolved a lower proliferation index and higher cell death rate compared to MTD protocols. In fact, across all the variations of protocols we tested, we found that the more drug that was used, the shorter the survival time and the more resistance evolved in the tumors. This implies, in the setting of late-stage cancers and second-line therapy, or any other setting where cancers are not being treated with intent to cure, we may be able to extend time to progression by reducing the dose of anti-cancer drugs, and perhaps even use evolutionary approaches to prevent the evolution of acquired therapeutic resistance. In the setting of endocrine-resistant breast cancer, our results also support clinician’s preference of capecitabine over gemcitabine. The success of adaptive therapy here could provide justification for a future clinical trial using dose-modulation adaptive therapy with capecitabine in ER+ metastatic, endocrine-resistantbut chemo-naive breast cancer. If a second drug were added to capecitabine, we would recommend using them in a dose-modulation, ping-pong on progression protocol in which you use the standard de-escalation of doses as long as the tumor is shrinking on a single drug, but instead of raising the dose if the tumor grows, switch to the other drug and resume de-escalation until progression (*14*). By changing the way that we use current drugs, so as to prevent the expansion of resistant clones, we have the opportunity to transform cancer from an acute, lethal disease, into one that is chronic and manageable.

## MATERIALS AND METHODS

### Study Design

The aim of this study was to test an evolution-based strategy to treat ER+ endocrine-resistant breast cancer. We first evolved the estrogen-positive (ER+) breast cancer cell line MCF7 to be resistant to fulvestrant (a selective estrogen receptor modulator [SERM]), and palbociclib (a CDK4/6 inhibitor), both of which are commonly used clinically for ER+ breast cancer. We then tested a variety of adaptive therapy protocols, compared to constant MTD therapy as well as vehicle control (no therapy), measuring time until death as the primary outcome.

In all cohorts the mice were randomly assigned to the following treatment and control groups (Table 1). The primary endpoint for the mouse experiments was time to death (see below). We monitored tumor burden by the Xenogen IVIS Spectrum in vivo imaging system.

**Table 1:**
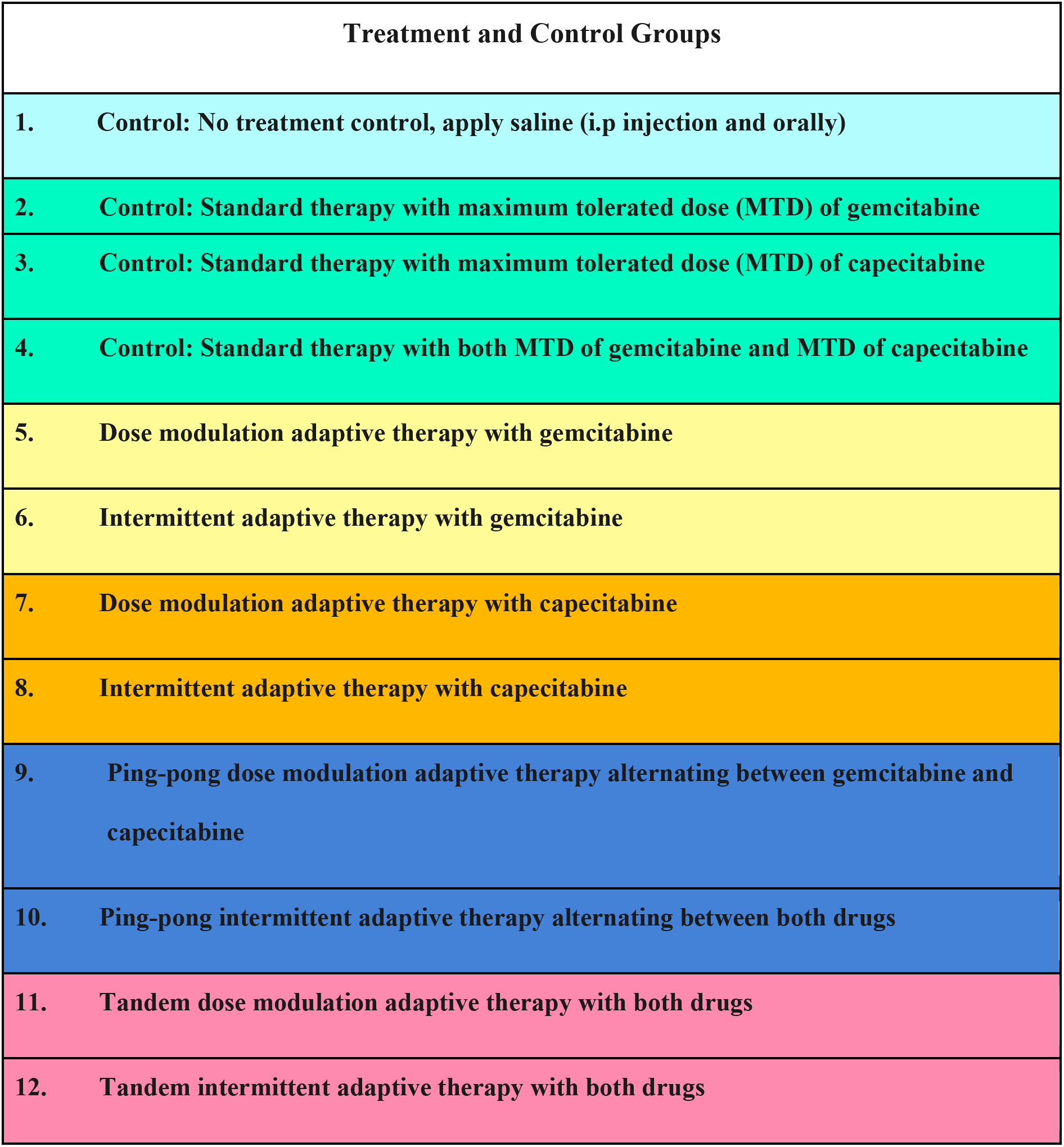
Treatment and control groups used in this preclinical experiment on endocrine-resistant breast cancer. Each treatment groups shown in different colors: vehicle in light blue, Control MTD (single and combination therapy) in green, Gemcitabine single therapy in yellow, Capecitabine single therapy in orange, combination pin-pong strategy in dark blue and combination tandem strategy in dark pink.

### In-Vitro Experiments

#### Cell Culture

##### Preparing Endocrine-Resistant Cell Lines

Human bioluminescent human ER+ breast cancer cell line, that was able to express firefly luciferase (MCF7/luc), was cultured in RPMI 1640 (Gibco, ThermoFisher) and supplemented with 10% fetal bovine serum (FBS, (Gibco, ThermoFisher)) and 1% Penicillin-Streptomycin, an antibiotic (Gibco, ThermoFisher). MCF7/luc human breast cancer cell lines were evolved to become resistant to fulvestrant and palbociclib. In order to obtain resistant cell lines, 1 × 10^7^ MCF7/luc cells were inoculated into a hyper-flask and grown for 1 week with RPMI 1640 media, 500 ml +10% FBS until they reached approximately 1 × 10^9^ cells. We then treated them for one month with 70% inhibitory concentration (IC70, 14.8 uM) for the first two weeks, IC80 (third week, 16 uM), and IC90 (last week, 18 uM) of the drugs. We changed the media of the hyperflask once per week when we added the drugs to the media. When changing the media, the media was poured into an empty bottle by holding the hyperflask at an angle followed by adding 100 ml of PBS to wash the flask. Finally, the fresh media with drugs was poured into the flask.

##### Drug Dose-Response Curve Analysis of Resistant Cell Lines

We analyzed the Drug Dose-response curve (DDR) on MCF7/luc cells that evolved to be resistant to fulvestrant and palbociclib versus the parental cell line (sensitive MCF7/luc cells) to compare their sensitivity to the drugs. We cultured the cells in Petri dishes with complete media (RPMI 1640 media, +10% FBS), then harvested the cell using Accutase to detach the cells followed by incubation in the incubator until the cells detached (about 5-10 minutes). Once the cells detached, equal amounts of complete media were added, washed, and pipetted to bring all the cells into suspension. The cells were transferred into a tube and then counted, followed by centrifuging for 5 min at 1100 rpm. The media was then removed and the pellet was resuspended in a suitable volume for plating cells in order to get 10^3^ cells per well.

We used a 96-well plate with 2 controls (media without drug and media plus the solvent of each drug) and 10 treatments (drugs with different concentrations, 30uM, 40uM, 50uM, 60uM, 70uM, 80uM, 90 uM, and 100uM), with 8 replicates per treatment. Equal amounts of cells were plated on all wells on Day 1. After 24 hours, the cells were treated with the drugs at the specified concentrations. The cells were incubated under the drugs for 72 hours to cover at least one doubling time of the cell lines (MCF7S cells had a doubling time of 41 hours and MCF7R cells had a doubling time of 48 hours). At the end of the experiment, an equal amount of 100ul of CellTitre-Glo (Promega) was added to each well, followed by incubation for five minutes. CellTitre-Glo releases ATP from live cells (*18*), so the quantified luminescence data using a plate reader (SpectraMax M5 with SoftMax Pro 6.2.2 software) corresponds to live cells under each condition. These values were then normalized to construct a drug dose-response curve (DDR). Statistical analysis was performed on DDR results using GraphPad Prism. First, we log-transformed the concentration and then used a non-linear regression with the following command: “(log(inhibitor) vs response - variable slope (four parameters))”, which gave us a corrected DDR curve and an IC50 value.

### In-Vivo Experiment

#### Xenograft Model of Human Breast Cancer

We orthotopically implanted endocrine-resistant MCF7 cell lines tagged with Firefly luciferase into the mammary fat pads of 8-week-old NOD/SCID gamma (NSG) mice. We used 3 x10^6^ cells/100ul for each mouse (suspended in DPBS/Matrigel 1:1). We used 6 mice per treatment group and 4 mice for the control group based on a power analysis: Balanced one-way analysis of variance power calculation for 11 experimental arms and controls: groups = 11, n = 6, f = 0.53, power = 0.8, sig. level = 0.05. A total of 70 mice were used for this study. Orthotopic mouse xenograft experimental protocols were approved by the Institutional Animal Care and Use Committee (IACUC) at Arizona State University. NSG mice are immunodeficient, so they were maintained and evaluated under pathogen-free conditions in accordance with IACUC standards of care at the ASU vivarium.

For harvesting 3 x10^6^ cells/mouse, the cells were first washed with Dulbecco’s phosphate-buffered saline (DPBS; Corning^TM^) for 15 minutes to remove any trace of FBS from the cells. Then the cells were detached using Accutase incubated at 37°C, 5% CO2 for 5-10 minutes. Then cells were suspended in RPMI 1640 growth medium +10% FBS to neutralize the Accutase followed by centrifuge for 5 minutes at 1100 RPM. The pellet of cells was resuspended and mixed with 1:1 matrigel: PBS (3e+10^6^ cells/100ul). Matrigel was used to augment the tumor growth for our cancer cell lines (*19*).

For the MCF7 xenograft model, 17b-estradiol 90-day release pellets, 0.36 mg per pellet (Innovative Research of America) were implanted on the dorsal region of the mice on the day of cell injections in order to grow human breast cancer cells that are estrogen receptor-positive (*20,21*). The incision was closed with surgical glue and the mouse was monitored and kept warm while recovering from the anesthesia. They were returned to their cage once they were recovered. This was repeated every 90 days while the mice were under observation.

#### Tumor Burden Measurement

Tumor burden was measured by caliper and bioluminescence imaging under isoflurane anesthesia twice a week. The tumor volume measured by the caliper was calculated according to the formula: volume = ***π*** (short diameter^2^) ✕ (long diameter)/6. Imaging of live mice using the IVIS® Spectrum revealed tumor size and density. For live imaging, we injected 100 ul of diluted XenoLight D-Luciferin-K+salt, a bioluminescence substrate (PerkinElmer, Inc) intraperitoneally at a concentration of 150 mg/kg body weight. Then, mice were anesthetized with isoflurane (3% induction dose and 1.5%-2% maintenance dose via a precision vaporizer) followed by being placed in the chamber for fluorescent imaging using IVIS Spectrum (PerkinElmer, Inc) and imaged ventrally.

#### Starting Therapy and Drug Dosing

When the measured tumor volume exceeded 200 mm^3^, according to calipers, or 3 × 10^8^ (photons/sec/cm^2^/sr)measured by bioluminescence imaging, mice were randomly divided into the control group and treatment groups. In treatment groups, we used gemcitabine at 50 mg/kg as the maximum tolerated dose (MTD) which we injected intraperitoneally twice weekly,and capecitabine given at 40 mg/kg/day as MTD which we administered orally with a 20 **μ**l pipette tip, five days (Monday-Friday) per week. For the control group (no treatment) we injected saline (the solvent for both gemcitabine and capecitabine) twice a week (as a control for the twice-weekly injection of gemcitabine), and applied saline orally twice a week (as a control for the thrice-weekly oral application of capecitabine).

#### Standard Therapy Protocol

To model standard therapy, we applied the MTD of a drug continuously: 40 mg/kg/day of capecitabine five days a week or 50 mg/kg of gemcitabine twice a week. In the combined standard therapy, mice were given both drugs at MTD on those same schedules (gemcitabine twice a week and capecitabine five days a week).

#### Dose Modulation Adaptive Therapy Protocol

We adjusted drug dosage based on the tumor burden twice per week and optimized our experimental protocol according to dose modulation strategy found in our agent-based modeling work (*14*). Specifically, if tumor burden decreased by >10% since the last measurement, based on bioluminescence, we decreased the dose by 50%. If the tumor burden increased by >10%, we increased the dose by 50% but not exceeding the MTD (40 mg/kg/day for capecitabine, and 50 mg/kg for gemcitabine). When the tumor shrank below 2.25 × 10^8^ (photons/sec/cm^2^/sr) or a minimum volume cutoff of 150 mm^3^ based on caliper measurements, we skipped the treatment until the tumor grew back to 3 × 10^8^ (photons/sec/cm^2^/sr).

#### Intermittent Adaptive Therapy Protocol

In this protocol we started with the maximum tolerated dose of the drug, 40 (mg/kg/day) of capecitabine or 50 (mg/kg/2 days in a week) of gemcitabine. If the tumor burden fell below 50% of its value at the start of treatment, we withdrew the treatment. Then, if it rose above the initial tumor burden at the beginning of treatment, we started the treatment again using the MTD of each drug.

#### Ping-Pong Dose Modulation Adaptive Therapy Protocol

We used the dose modulation protocol as described above, except that we alternated between applying gemcitabine and capecitabine as follows:

Monday: Dose with gemcitabine.
Tuesday: Measure tumor burden and set the dose for the next application of gemcitabine (the following Friday) based on the tumor burden difference since last Friday. Start dosing with capecitabine.
Wednesday: Dose with capecitabine.
Thursday: Dose with capecitabine.
Friday: Measure tumor burden and set the dose for the next application of capecitabine on the following Tuesday, based on the change in tumor burden since Tuesday. Dose with gemcitabine.
Weekend: No treatment.

The dose was adjusted as described in the dose modulation protocol, adjusting the dose of a drug depending on whether the tumor grew or shrank the last time that drug was applied. Note that gemcitabine has a half-life of less than 1 hour in mice (*22*) and capecitabine has a half-life of 1-4 hours in mice (*22*).

#### Ping-Pong Intermittent Adaptive Therapy Protocol

We used one drug at a time, and like the single drug intermittent adaptive therapy protocol above, we stopped treatment when the tumor burden fell below 50%. The only difference was that when the tumor burden returned to its initial value prior to treatment, we switched drugs rather than continuing with the same drug.

#### Tandem Dose Modulation Adaptive Therapy Protocol

This protocol was the same as dose modulation adaptive therapy protocol above, except that both drugs were given and modulated in tandem (gemcitabine twice a week and capecitabine five days a week, on the weekdays). We measured tumor burden twice a week, and when the tumor grew more than 10% since the last measurement, we increased the dose of both drugs by 50% (up to but never going above their MTD). Similarly, if the tumor burden shrank by more than 10%, we decreased the dose of both drugs by 50%.

#### Tandem Intermittent Adaptive Therapy Protocol

This was the same as the intermittent adaptive therapy protocol except that both drugs were given at the same time. Both drugs were also withdrawn if the tumor burden fell below 50% of its initial burden and restarted if the tumor burden ever grew above 100% of its initial burden.

#### Endpoints

The primary endpoint for the mouse experiments was time to death. Mice were monitored daily and euthanized if they developed excessive lethargy, inappetence, excessive tumor burden, or other serious clinical conditions as determined by the veterinary staff such as difficult labored breathing, tumor interfering with locomotor activity, or ability to obtain food or water. Excessive tumor burden was defined as 3 × 10^9^ (photons/sec/cm^2^/sr) according to the luminescence data from IVIS or 2000 mm^3^ based on the calipers. Mice that had to be sacrificed due to the tumor interfering with locomotion or died during the surgery to implant the estrogen pellets were coded as censored for survival, as long as the caliper measurements of tumor burden were below the threshold for sacrifice. All other reasons for sacrifice were coded as a mortality event, including sacrifice for excessive lethargy, inappetence, excessive tumor burden, or other serious clinical conditions as determined by the veterinary staff such as difficult labored breathing, hunching, or ability to obtain food or water (table S.1).

### Ex-Vivo Experiments

#### Derivation of Cancer Cell Lines from the Mice

When mice were euthanized, we harvested the tumors. Half of the tumor was used for deriving a cell line and the other half was saved for tissue-based assays. To derive a cell line, the tumor was sliced and diced into small pieces and then 7 ml of the digest media (6ml collagenase (3mg/ml PBS) + 10 ml TrypLE Express + 34 ml 1X sterile PBS) were added to the plate and incubated for 30 minutes at 37°C. After incubation, the digest media and cells were separated from any remaining tissue and transferred into a conical tube. Then another 3 ml digest media was added to the remaining tissue and sliced again followed by adding 7 ml digest media and incubating the plate in a tissue culture incubator for another 30 minutes. Next, we collected media and cells from the dish and added them to the same conical tube we had. Finally, we plated the cells in a petri dish with complete media that contained Normocin at a 1:500 ratio and put it in the incubator to grow. Then cell lines were frozen and kept at -80 °C.

#### Histological Analysis and Immunohistochemistry

The samples were embedded in a paraffin block for hematoxylin and eosin (H&E) staining and immunohistochemistry (IHC). Tumor tissue was sectioned in 6 um slices using Leica Ultracut-R Microtome. We performed IHC using the following antibodies: Ki67 is used as a marker of proliferation, to assess the proliferative activity of the tumor and to determine tumor response to our treatment protocols (Ki-67 Recombinant Rabbit Monoclonal Antibody (SP6), invitrogen#MA5-14520), and caspase 3 (Cleaved Caspase-3 (Asp175) Antibody #9661, Cell Signaling) which is an important component of apoptosis (Supplementary methods, Data S1).

#### Statistical Methods

We analyzed the survival results using Kaplan-Meier analyses and Cox proportional hazards models. We analyzed our histology and IHC comparing MTD and adaptive therapy groups with the Mann-Whitney test.

#### Computational Modeling

We modified our previously published hybrid agent-based model (*14*), which is an extension to the Hybrid Automata Library (HAL) agent-based modeling framework (*23*), to simulate adaptive therapy using a single cytotoxic or two cytotoxic drugs. The description of the model can be found in (*14*), with the following changes:

In section 2.2.2. (Entities, State Variables, and Scales), for treatment with a single cytotoxic drug, we have considered two different cell types: sensitive and resistant. In section 2.4.11 (Observation), we modified our criterion for progression: At any point after the initiation of therapy, if the tumor burden equalled or exceeded 99% of the carrying capacity, or if the rolling average of the resistant cells (doubly resistant for treatment with two drugs or single resistant for treatment with one drug) over 500 time-steps equaled or exceeded 50% of the carrying capacity, then the particular run is scored as “Progressed”. In section 2.5 (Initialization), we consider two different cell types–sensitive and resistant. In section 2.7.1 (Cell Death), for treatment with a single cytotoxic drug, the equation for the probability of cell death is as follows: Probability of cell death per hour = background death probability per hour+S1*[Drug1]\***Ψ**1, where S1 is the binary indicator variable for the cell’s sensitivity to drug 1, [Drug1] being the concentration of drug 1 (non-negative real values), and **Ψ**1 is the drug potency (non-negative real values), quantified as the probability of cell death per unit drug concentration per hour. In section 2.7.2 (Cell Division), the cell division rates for the sensitive cells is 0.06 per hour and the cell division rate for the resistant cells is 0.02 per hour. In section 2.7.4 (Mutation), for treatment with a single cytotoxic drug, the default value for the mutation rate parameter is 1 × 10^-3^ per cell division, accounting for the transition from sensitive to resistant cell types, and vice versa. Protocols for treatment with a single cytotoxic drug and two cytotoxic drugs described in Supplementary method (Data S2). The parameter values that were used to run the model for treatment using a single cytotoxic drug are shown in Table 2, and that were used to run the model for treatment using two cytotoxic drugs are indicated in Table 2. MTD dosage value for treatment using a single drug was set to 2.5 units, and MTD dosage value for treatment using two cytotoxic drugs was set to 2.5 units for each of the two drugs, thus the total dosage equaled 5 units for a cocktail application, and 2.5 units of the specific drug for the ping-pong protocols.

**Table 2:** Parameter values for matching the results of the mice experiments using a single cytotoxic drug.

| Parameter | Value |
| --- | --- |
| Cell division rate: sensitive | 0.06 per hour |
| Cell division rate: resistant | 0.02 per hour |
| Background death rate | 0.01 per hour |
| *Replacement probability* | *1.0* |
| Delta Tumor | 10% |
| Delta Dose | 50% |
| Probability of death due to drug potency ($\Psi$) | 0.04 per unit drug concentration |
| Maximum tolerated dose (MTD) | 2.5 units |
| Minimum drug dose | 0.5 units |
| Drug on time | 1 hour |
| Frequency of drug application | Once every 24 hour |
| Check tumor burden | Every 3 days |
| Drug decay | 10% per hour |
| Drug diffusion rate | 2.0 |
| Tumor size triggering treatment | Tumor burden is 50% or more of the carrying capacity |
| Mutation rate | 1e-3 per cell division |
| Measurement noise standard deviation (SD) | 5 cells |
| Total grid size | 100 by 100 |
| Duration of simulation | 5000 hour |
| Stop dosing/initiate treatment vacation when (DM protocols only): | Tumor burden is less than or equal to 25% of carrying capacity |
| Doubling time of sensitive cells | 13.86 hour |
| Doubling time of resistant cells | 69.3 hour |

**Table 2:**
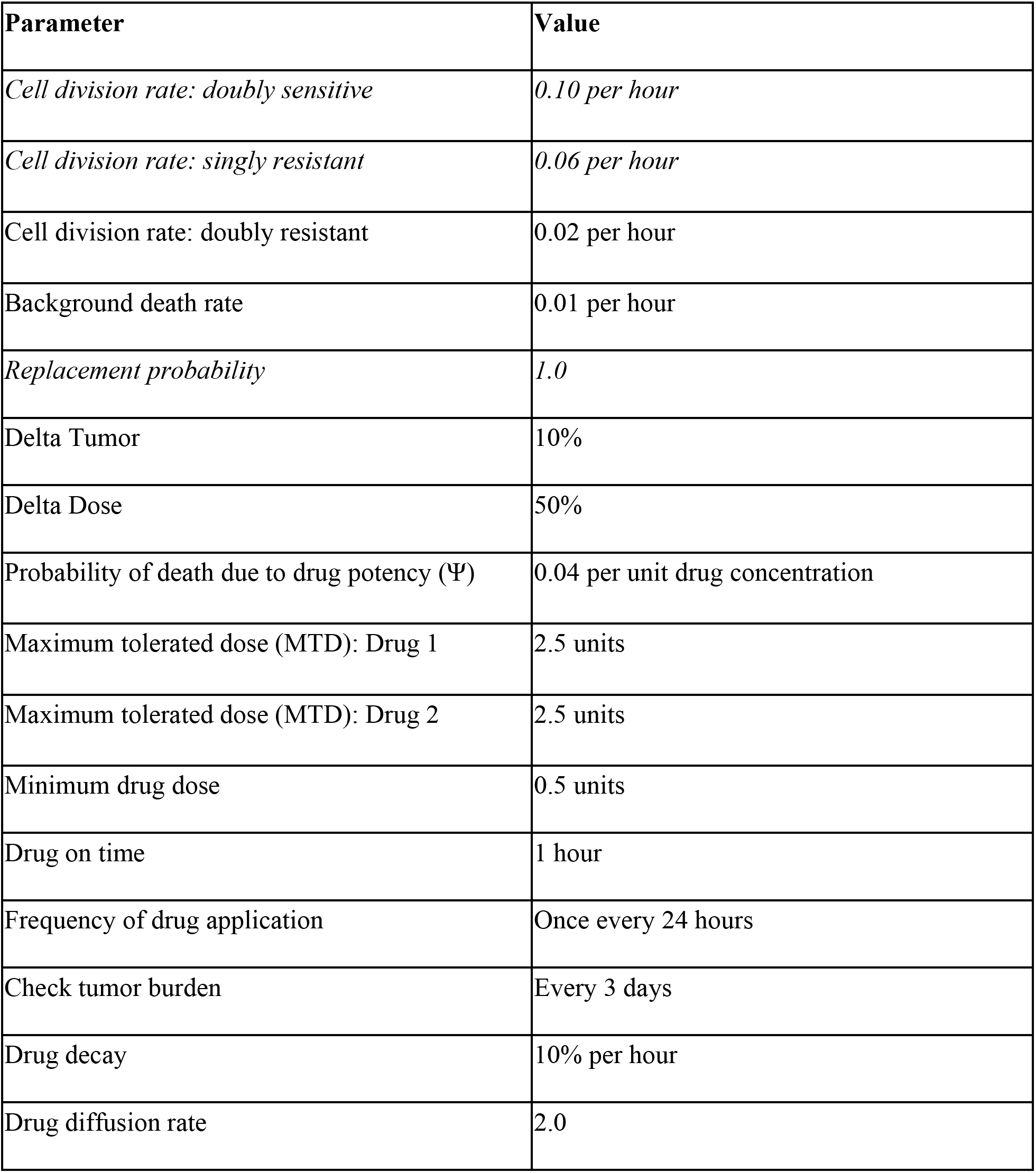
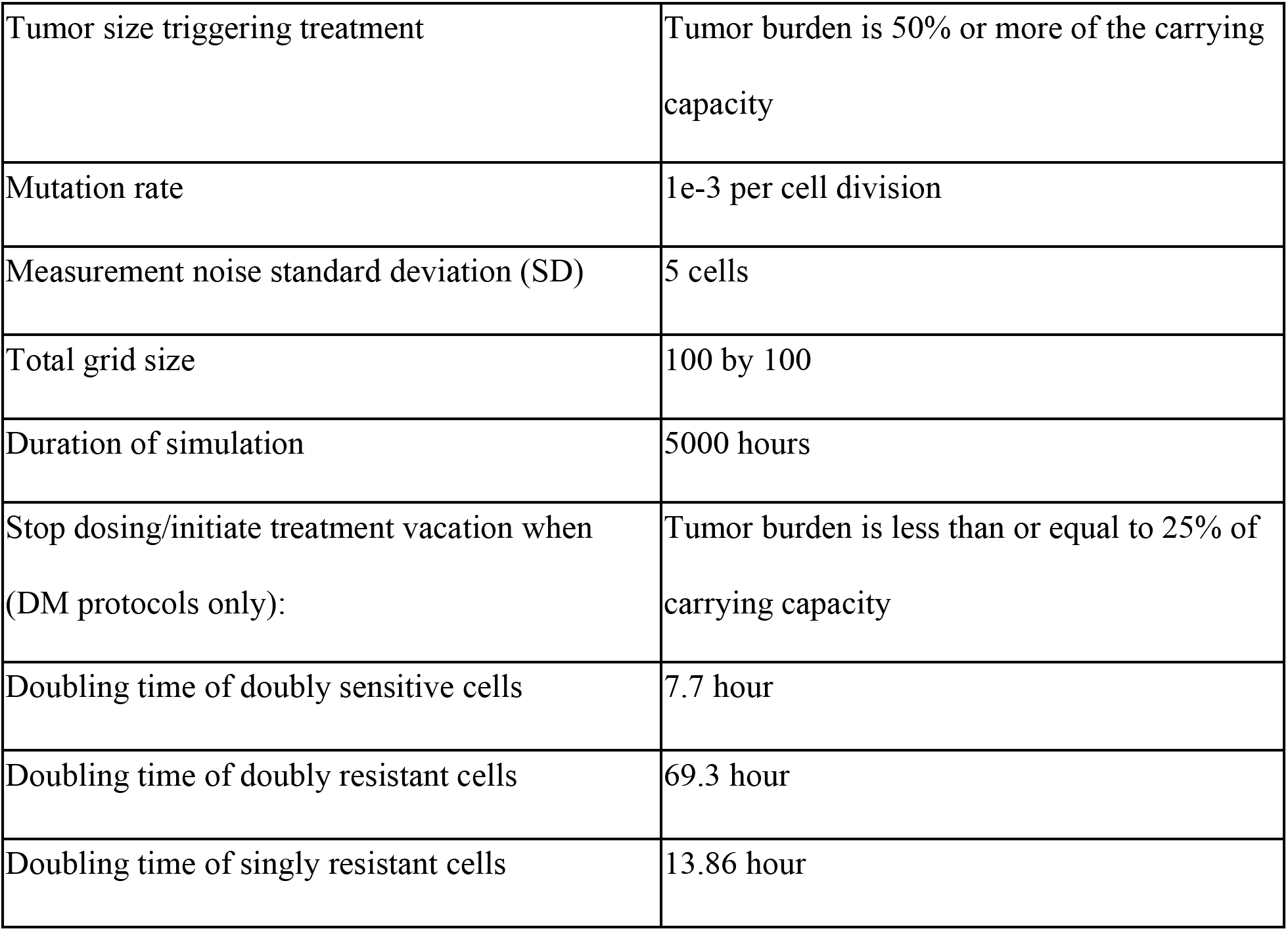
Parameter values for matching the results of the mice experiments using two cytotoxic drugs.

| Parameter | Value |
| --- | --- |
| *Cell division rate: doubly sensitive* | *0.10 per hour* |
| *Cell division rate: singly resistant* | *0.06 per hour* |
| Cell division rate: doubly resistant | 0.02 per hour |
| Background death rate | 0.01 per hour |
| *Replacement probability* | *1.0* |
| Delta Tumor | 10% |
| Delta Dose | 50% |
| Probability of death due to drug potency ($\Psi$) | 0.04 per unit drug concentration |
| Maximum tolerated dose (MTD): Drug 1 | 2.5 units |
| Maximum tolerated dose (MTD): Drug 2 | 2.5 units |
| Minimum drug dose | 0.5 units |
| Drug on time | 1 hour |
| Frequency of drug application | Once every 24 hours |
| Check tumor burden | Every 3 days |
| Drug decay | 10% per hour |
| Drug diffusion rate | 2.0 |
| Tumor size triggering treatment | Tumor burden is 50% or more of the carrying capacity |
| Mutation rate | 1e-3 per cell division |
| Measurement noise standard deviation (SD) | 5 cells |
| Total grid size | 100 by 100 |
| Duration of simulation | 5000 hours |
| Stop dosing/initiate treatment vacation when (DM protocols only): | Tumor burden is less than or equal to 25% of carrying capacity |
| Doubling time of doubly sensitive cells | 7.7 hour |
| Doubling time of doubly resistant cells | 69.3 hour |
| Doubling time of singly resistant cells | 13.86 hour |

This matched the mouse experiments in which the dosage of gemcitabine and capecitabine were not lowered when they were used in combination.

## LIST OF SUPPLEMENTARY MATERIALS

Fig S1 to S4 for supplementary figures

Table S1 to S3 for supplementary tables

Data S1 to S2 for supplementary Methods

## ACKNOWLEDGMENTS

The findings, opinions, and recommendations expressed here are those of the authors and not necessarily those of the universities where the research was performed or the National Institutes of Health. This work was supported in part by NIH grants U54 CA217376, U2C CA233254, U01CA232382 as well as CDMRP Breast Cancer Research Program Award BC132057 and the Arizona Biomedical Research Commission grant ADHS18-198847 to CCM and support from the Moffitt Center of Excellence for Evolutionary Therapy.The authors have no competing interests, all authors have approved the manuscript for submission and the content of the manuscript has not been published or submitted for publication elsewhere.

