## Supplementary material for "Testing Adaptive Therapy Protocols using Gemcitabine and Capecitabine on a Mouse Model of Endocrine-Resistant Breast Cancer": SUPPLEMENTARY MATERIALS.pdf

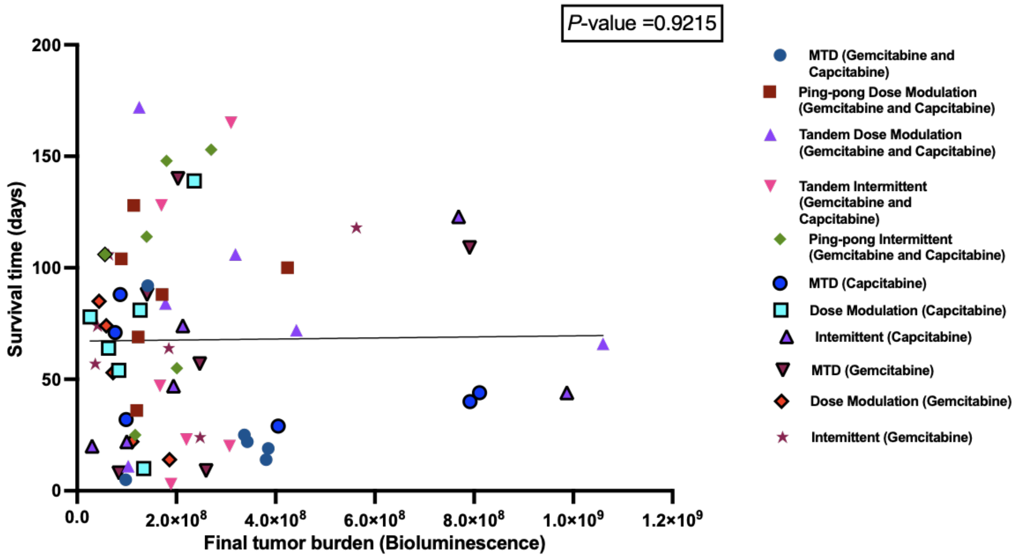


**Figure S1. Linear regression analysis of final Tumor burden with survival time (days) for all groups of this study.** There was no significant correlation between survival time to the last measured tumor burden (P-value = 0.92, $r^2 = 0.04$).

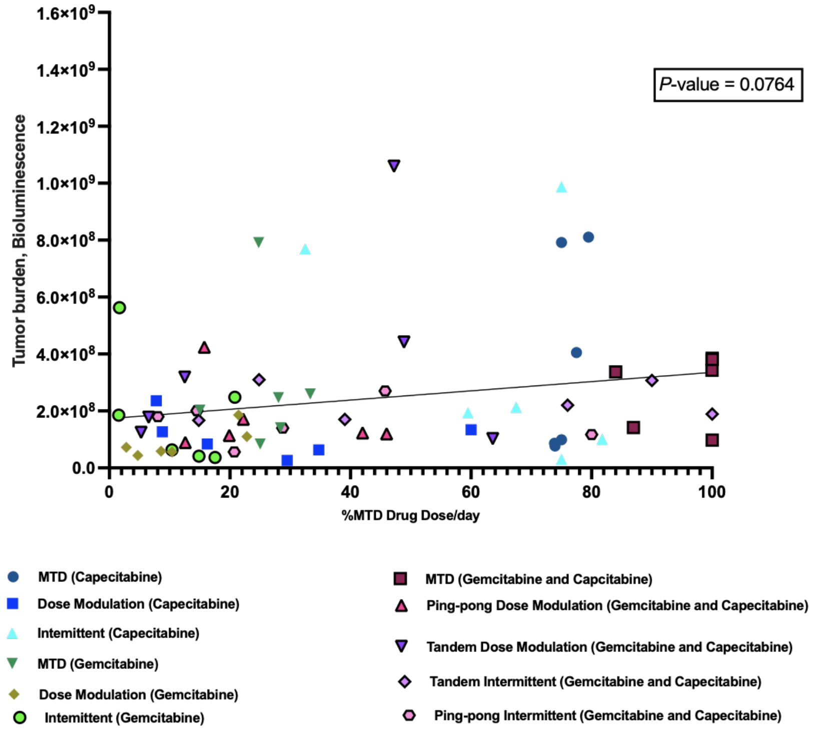


**Figure S2. Linear regression analysis of %MTD drug dose per day of mice received to final tumor burden.** There was no significant correlation between the %MTD drug dose that each mouse received per day to last measured tumor burden for that mouse (P-value = 0.076, $r^2$ = 0.048).

A

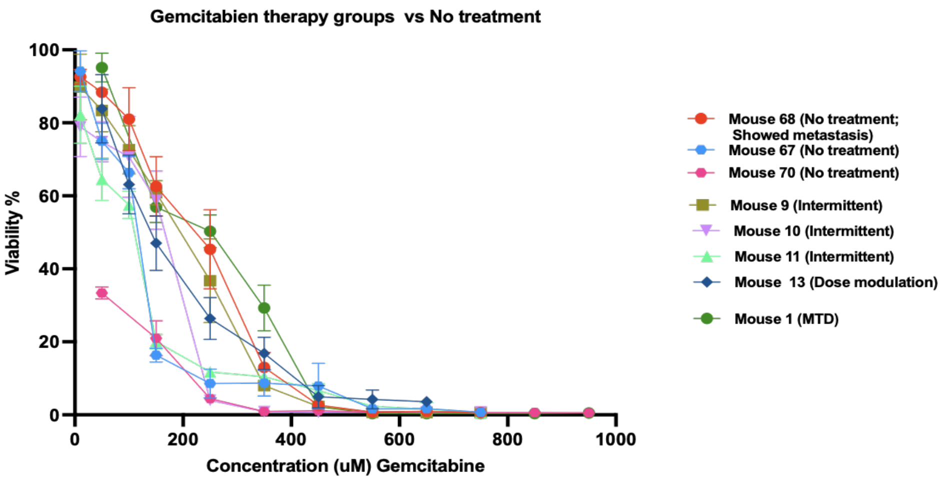
Gemcitabien therapy groups vs No treatment
Viability %
Concentration (uM) Gemcitabine
Mouse 68 (No treatment; Showed metastasis)
Mouse 67 (No treatment)
Mouse 70 (No treatment)
Mouse 9 (Intermittent)
Mouse 10 (Intermittent)
Mouse 11 (Intermittent)
Mouse 13 (Dose modulation)
Mouse 1 (MTD)

B

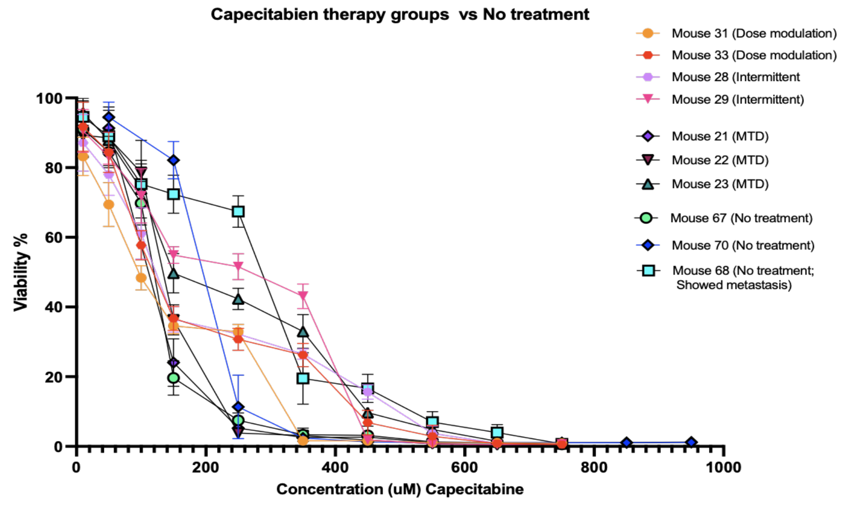
Capecitabien therapy groups vs No treatment
Viability %
Concentration (uM) Capecitabine
Mouse 31 (Dose modulation)
Mouse 33 (Dose modulation)
Mouse 28 (Intermittent
Mouse 29 (Intermittent)
Mouse 21 (MTD)
Mouse 22 (MTD)
Mouse 23 (MTD)
Mouse 67 (No treatment)
Mouse 70 (No treatment)
Mouse 68 (No treatment; Showed metastasis)

C

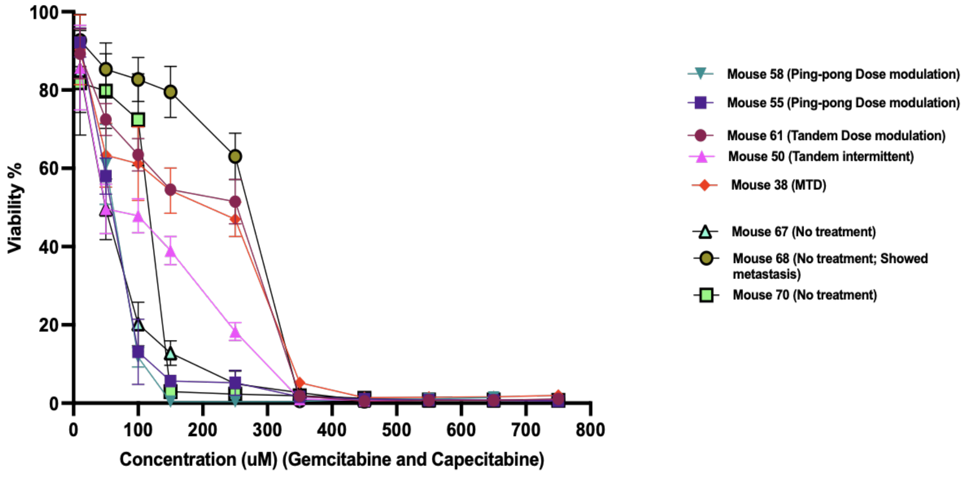
Viability %
Concentration (uM) (Gemcitabine and Capecitabine)
Mouse 58 (Ping-pong Dose modulation)
Mouse 55 (Ping-pong Dose modulation)
Mouse 61 (Tandem Dose modulation)
Mouse 50 (Tandem intermittent)
Mouse 38 (MTD)
Mouse 67 (No treatment)
Mouse 68 (No treatment; Showed metastasis)
Mouse 70 (No treatment)

**Figure S3. DDR analysis of Cell Lines Retrieved from Different Treatment Groups. (A)** DDR analysis of cell lines retrieved from the mice treated with gemcitabine single therapy. DDR comparison of adaptive therapy groups and MTD with the cell lines retrieved from the mice in no treatment arm and then treated with gemcitabine. **(B)** DDR analysis of cell lines retrieved from the mice treated with capecitabine single therapy. DDR comparison of adaptive therapy groups and MTD with the cell lines retrieved from the mice in no treatment arm and then treated with capecitabine. **(C)** DDR analysis of cell lines retrieved from the mice treated with a combination of capecitabine and gemcitabine. DDR comparison of adaptive therapy groups and MTD with the cell lines retrieved from the mice in no treatment arm and then treated with a combination of capecitabine and gemcitabine.

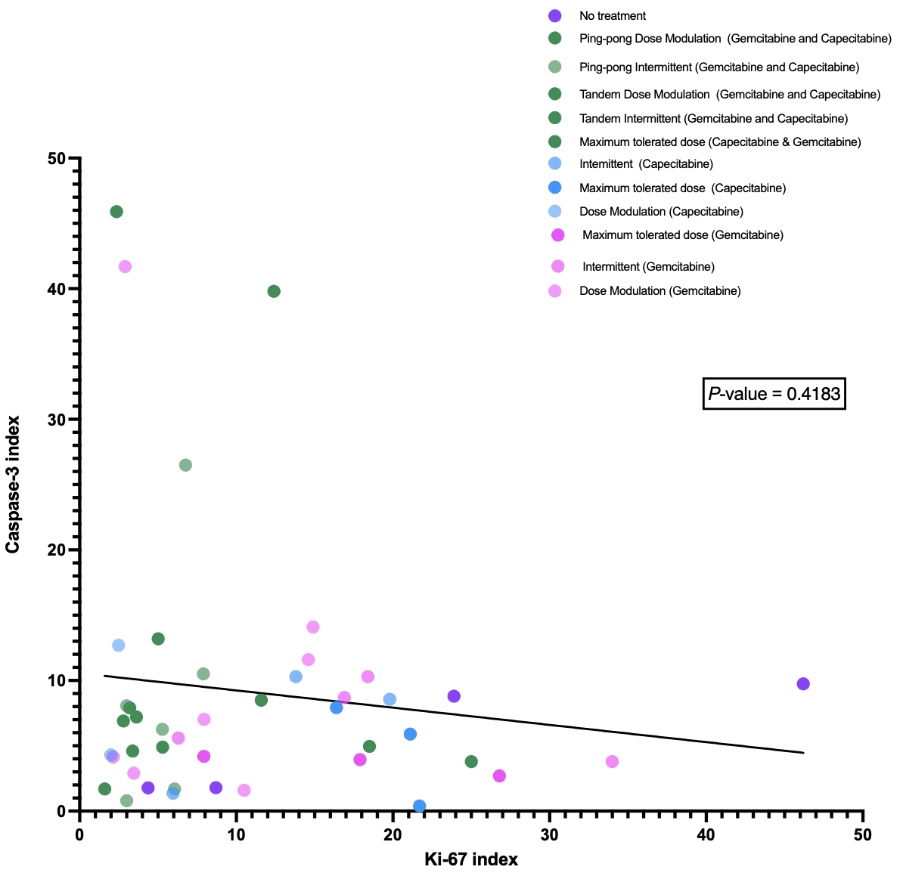


**Figure S4. Linear regression analysis between caspases-3 and Ki-67 Immunohistochemistry analysis of tumors of mice from different arms of our preclinical breast cancer model.** This result shows there is a negative correlation but not significant between Ki-67 and caspase-3 expression (Linear regression, P-value = 0.4183; $r^2 = 0.015$).

### Table S1: Tumor burden, drug dosing and outcome for each mouse.

<table>
<tr><th>Number of mouse and group of treatment</th><th>Reason of euthanizing mouse</th><th>Tumor burden graph during the time of study</th></tr>
<tr><td colspan="3">Capecitabine Dose Modulation</td></tr>
<tr><td>Mouse 31 Capecitabine Dose modulation</td><td>End point tumor size by caliper</td><td>
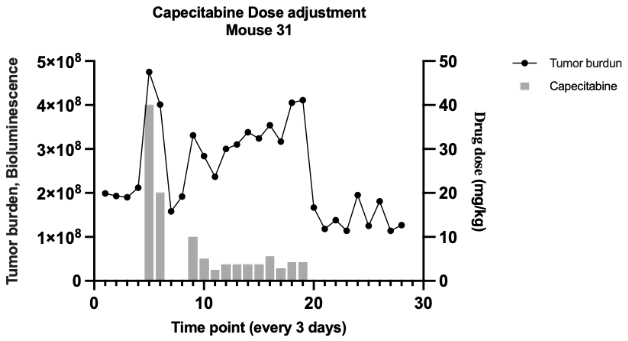

</td></tr>
<tr><td>Mouse 32 Capecitabine Dose modulation</td><td>Tumor size interfered with walking</td><td>
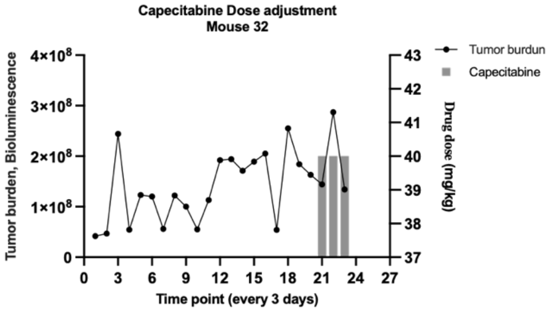

</td></tr>
<tr><td>Mouse 33 Capecitabine Dose modulation</td><td>Main tumor was small but it showed metastasis to underarm</td><td>
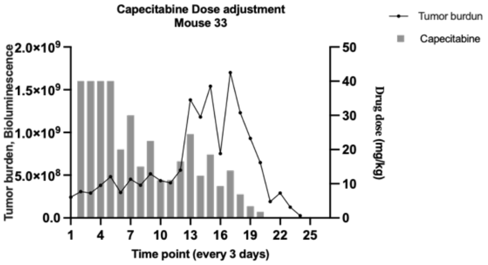

</td></tr>
</table>

<table>
<tr><td>Mouse 34 Capecitabine Dose modulation</td><td>Tumor size interfere walking</td><td>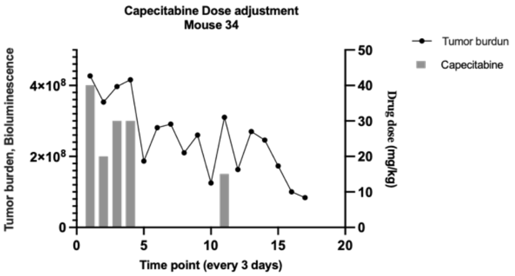
</td></tr>
<tr><td>Mouse 35 Capecitabine Dose modulation</td><td>Tumor size interfere walking</td><td>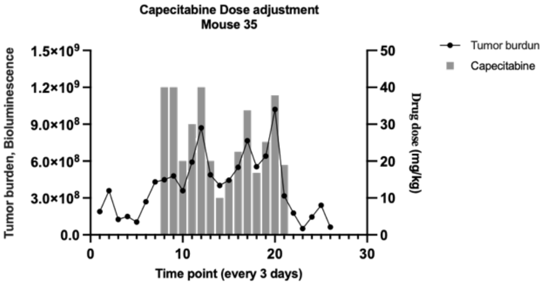
</td></tr>
<tr><td>Mouse 36 Capecitabine Dose modulation</td><td>Died</td><td>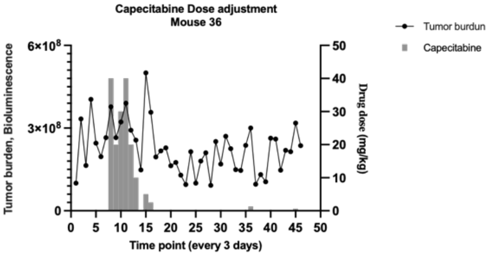
</td></tr>
<tr><td colspan="3">Capecitabine Intermittent</td></tr>
<tr><td>Mouse 25 Capecitabine Intermittent</td><td>Interfere with walking</td><td>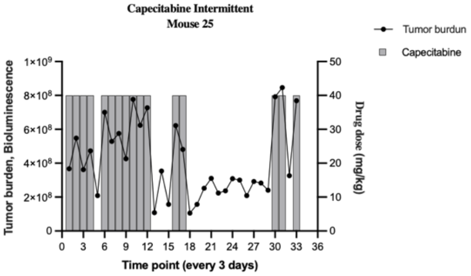
</td></tr>
</table>

| | | |
|---|---|---|
| **Mouse 26 Capecitabine Intermittent** | **Main tumor was small but it showed metastasis to underarm** | 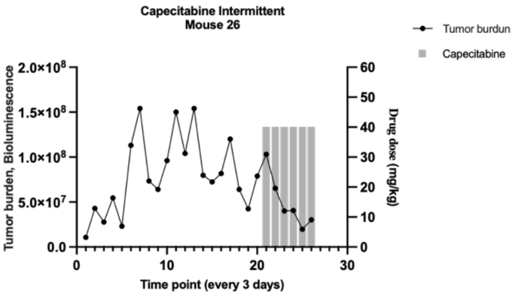 |
| **Mouse 27 Capecitabine Intermittent** | **Interfere with walking and skinny** | 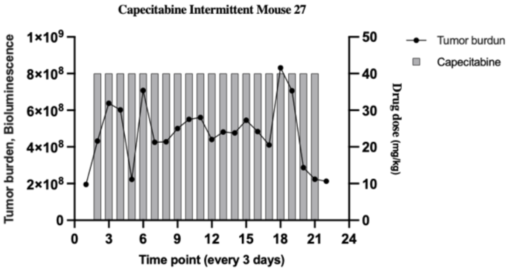 |
| **Mouse 28 Capecitabine Intermittent** | **Skinny** | 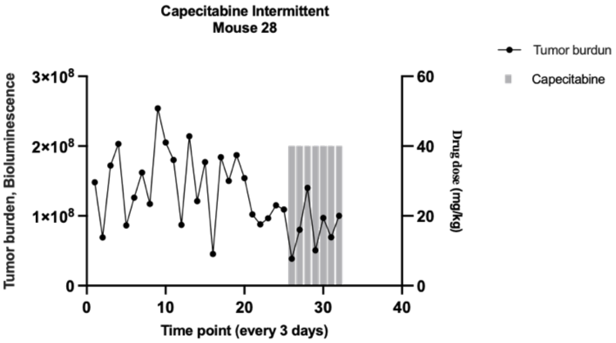 |
| **Mouse 29 Capecitabine Intermittent** | **Tumor size endpoint caliper** | 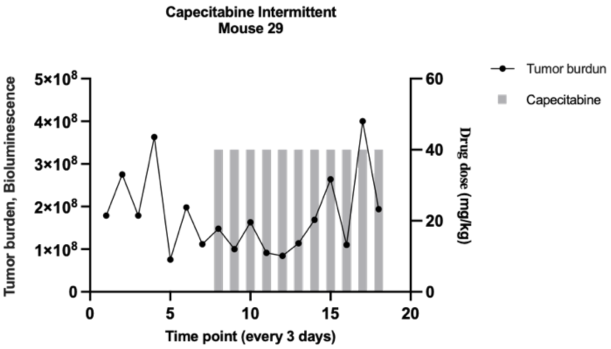 |

<table>
<tr><td>Mouse 30 Capecitabine Intermittent</td><td>Tumor size endpoint caliper</td><td>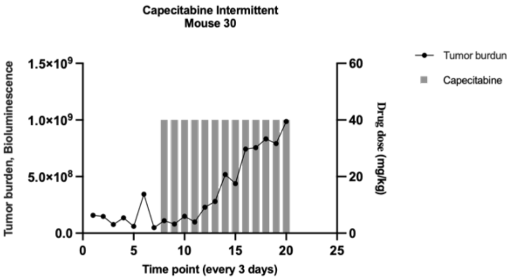
</td></tr>
<tr><td colspan="3">Capecitabine Maximum tolerated dose</td></tr>
<tr><td>Mouse 19 Capecitabine Maximum tolerated dose</td><td>Hunched</td><td>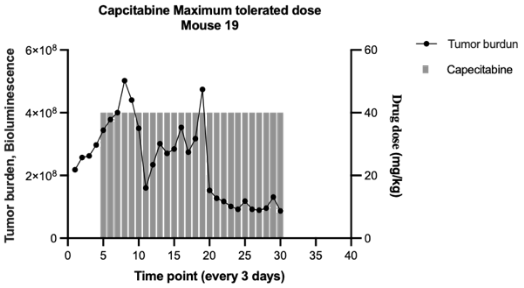
</td></tr>
<tr><td>Mouse 20 Capecitabine Maximum tolerated dose</td><td>Tumor size end point caliper</td><td>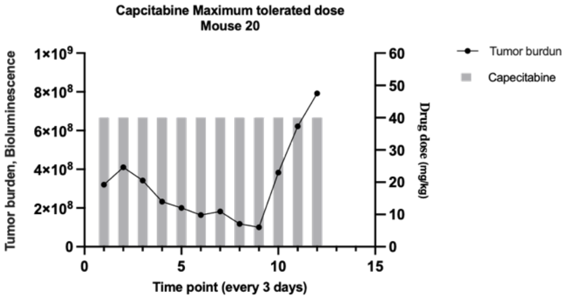
</td></tr>
<tr><td>Mouse 21 Capecitabine Maximum tolerated dose</td><td>Tumor size end point caliper</td><td>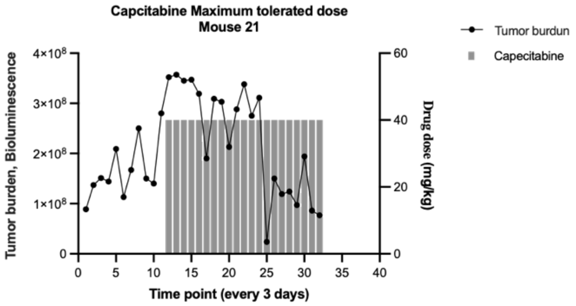
</td></tr>
</table>

<table>
<tr><td>Mouse 22 Capecitabine Maximum tolerated dose</td><td>Skinny, Hunched</td><td>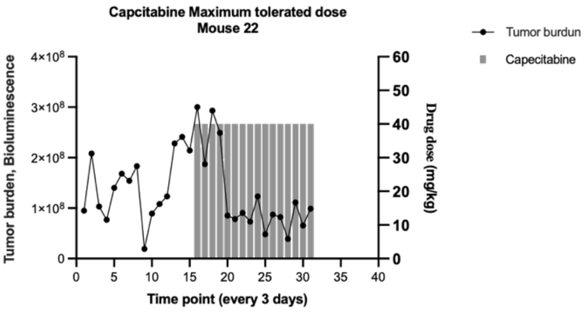
</td></tr>
<tr><td>Mouse 23 Capecitabine Maximum tolerated dose</td><td>Tumor size end point caliper</td><td>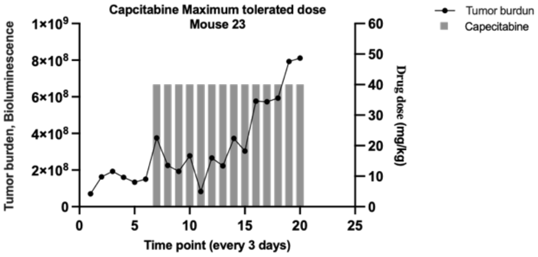
</td></tr>
<tr><td>Mouse 24 Capecitabine Maximum tolerated dose</td><td>Tumor size end point caliper</td><td>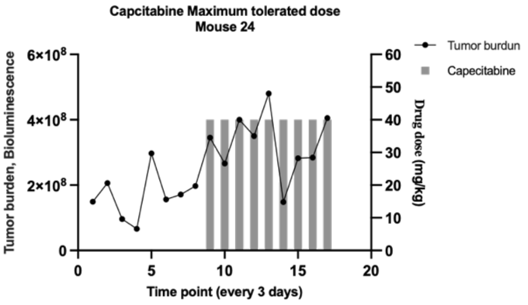
</td></tr>
<tr><td colspan="3">Gemcitabine Dose modulation</td></tr>
<tr><td>Mouse 13 Gemcitabine Dose modulation</td><td>Died</td><td>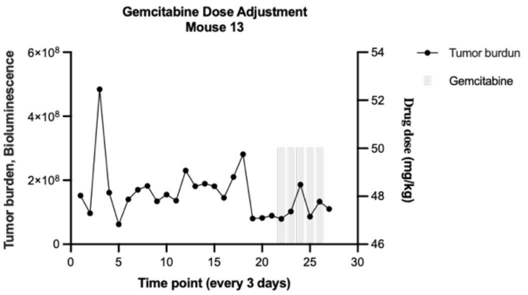
</td></tr>
</table>

<table>
<tr><td>Mouse 14 Gemcitabine Dose modulation</td><td>Died</td><td>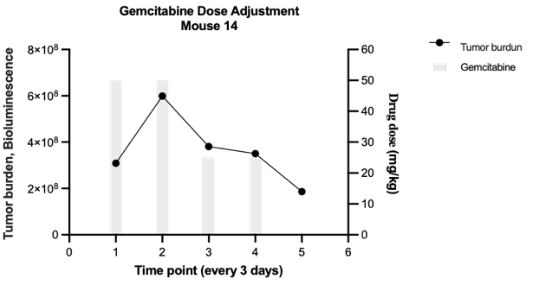
</td></tr>
<tr><td>Mouse 15 Gemcitabine Dose modulation</td><td>Tumor size end point caliper</td><td>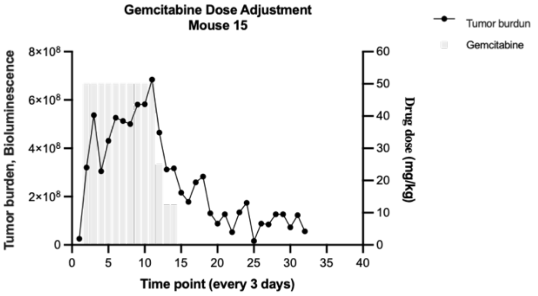
</td></tr>
<tr><td>Mouse 16 Gemcitabine Dose modulation</td><td>Tumor size end point caliper</td><td>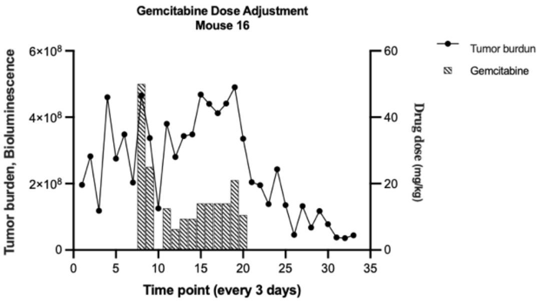
</td></tr>
<tr><td>Mouse 17 Gemcitabine Dose modulation</td><td>Tumor size end point caliper</td><td>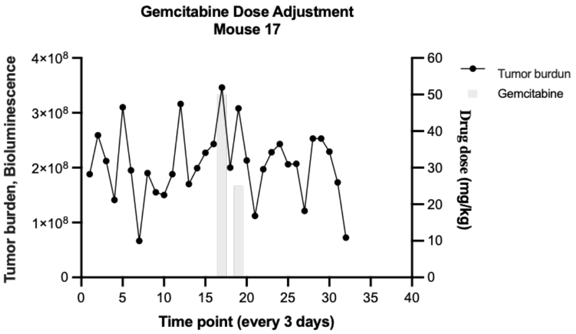
</td></tr>
</table>

<table>
<tr><td>Mouse 18 Gemcitabine Dose modulation</td><td>Died (skinny, hunched)</td><td>
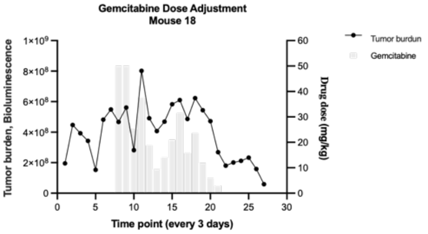

</td></tr>
<tr><td colspan="3">Gemcitabine Intermittent</td></tr>
<tr><td>Mouse 7 Gemcitabine Intermittent</td><td>Died</td><td>
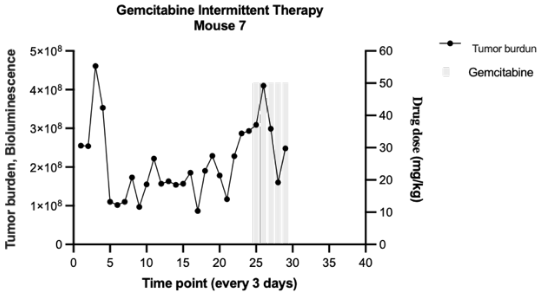

</td></tr>
<tr><td>Mouse 8 Gemcitabine Intermittent</td><td>Tumor size end point caliper</td><td>
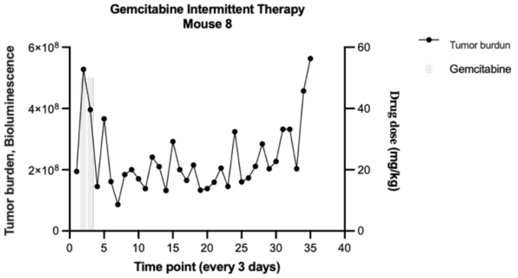

</td></tr>
<tr><td>Mouse 9 Gemcitabine Intermittent</td><td>Tumor size end point caliper</td><td>
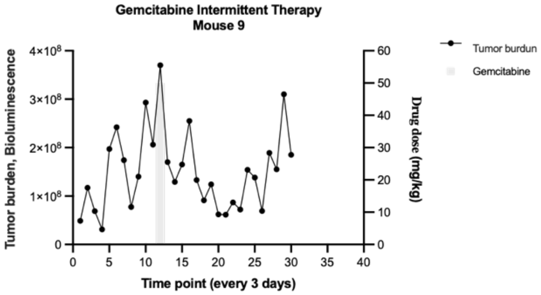

</td></tr>
</table>

| | | |
|---|---|---|
| **Mouse 10 Gemcitabine Intermittent** | **Main tumor was small but it showed metastasis to underarm** | 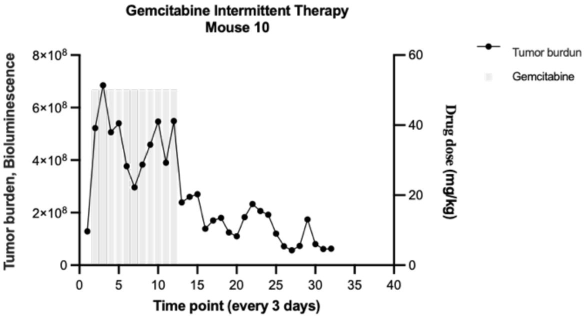 |
| **Mouse 11 Gemcitabine Intermittent** | **skinny, hunched** | 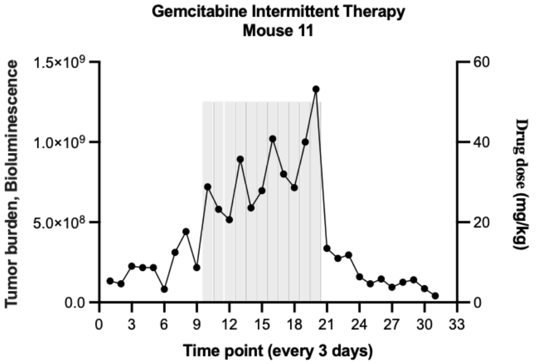 |
| **Mouse 12 Gemcitabine Intermittent** | **Died** | 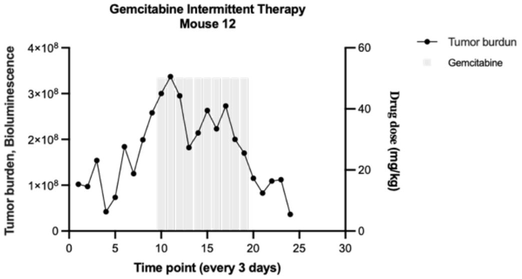 |
| **Gemcitabine Maximum tolerated dose** | | |
| **Mouse 1 Gemcitabine Maximum tolerated dose** | **Weight loss** | 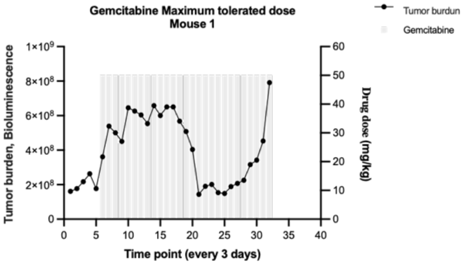 |

| | | |
|---|---|---|
| **Mouse 2 Gemcitabine Maximum tolerated dose** | **Died** | 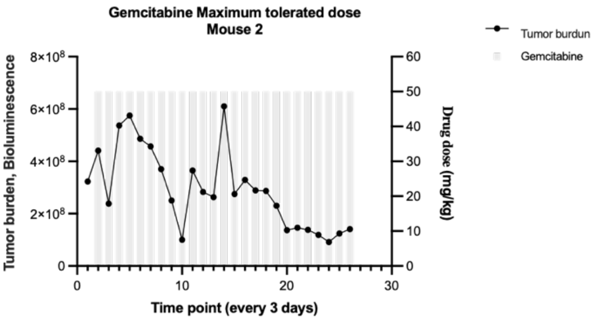  |
| **Mouse 3 Gemcitabine Maximum tolerated dose** | **Died** | 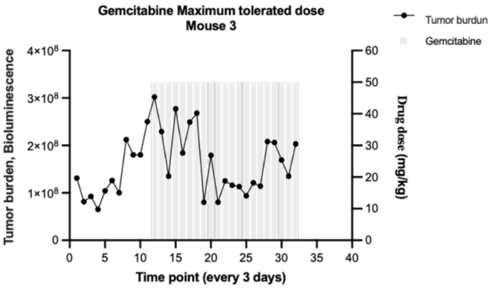  |
| **Mouse 4 Gemcitabine Maximum tolerated dose** | **Skinny (weight loss)** | 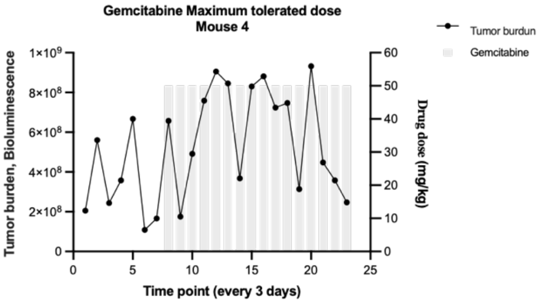  |
| **Mouse 5 Gemcitabine Maximum tolerated dose** | **Died** | 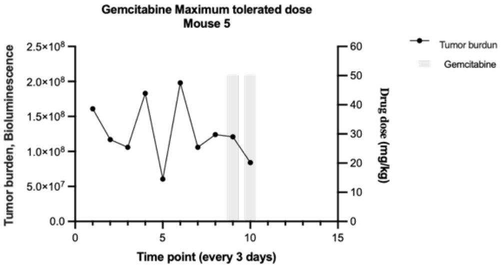  |

| | | |
|---|---|---|
| **Mouse 6 Gemcitabine Maximum tolerated dose** | **Tumor size endpoint Caliper** | 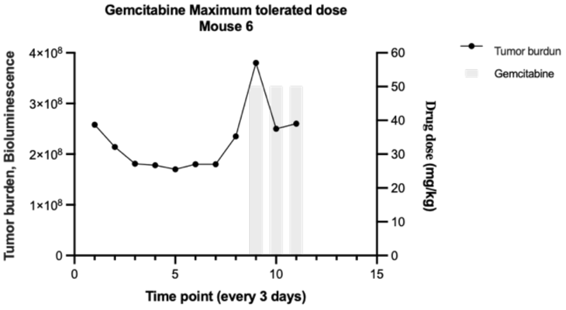 |
| **Gemcitabine & Capecitabine Maximum tolerated dose** | | |
| **Mouse 37 Gemcitabine & Capecitabine Maximum tolerated dose** | **Skinny (weight loss)** | 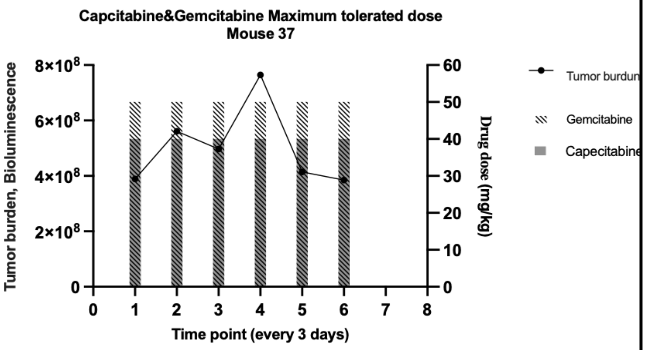 |
| **Mouse 38 Gemcitabine & Capecitabine Maximum tolerated dose** | **Skinny (weight loss)** | 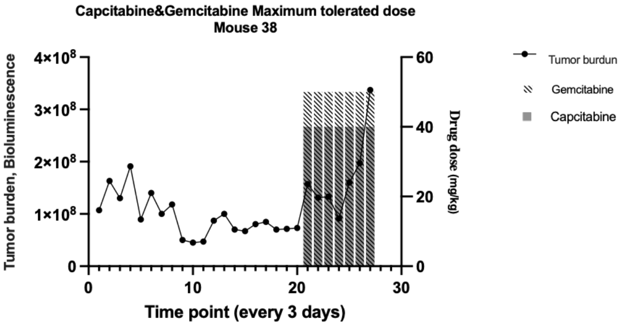 |
| **Mouse 39 Gemcitabine & Capecitabine Maximum tolerated dose** | **Skinny (weight loss)** | 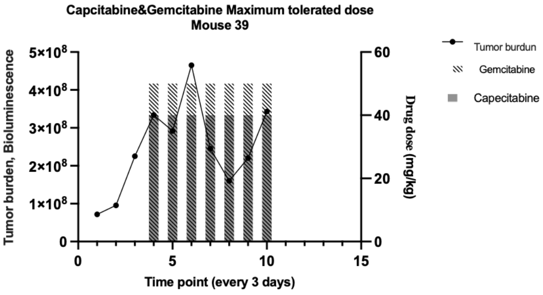 |

<table>
<tr><td>Mouse 40 Gemcitabine & Capecitabine Maximum tolerated dose</td><td>Skinny (weight loss)</td><td>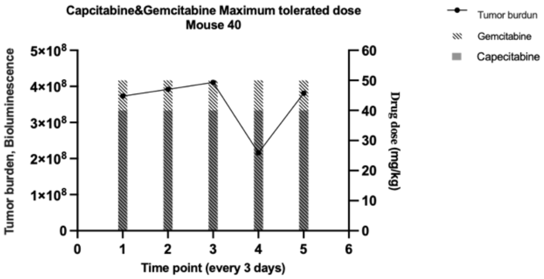
</td></tr>
<tr><td>Mouse 41 Gemcitabine & Capecitabine Maximum tolerated dose</td><td>Very weak and Skinny (weight loss)</td><td>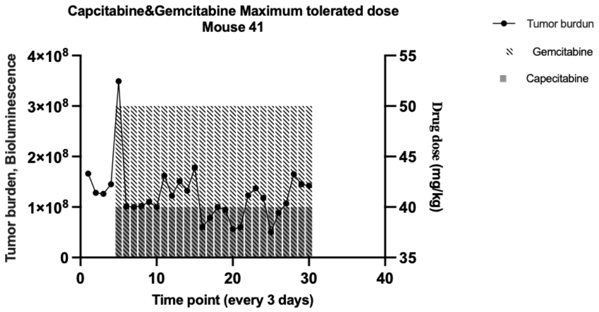
</td></tr>
<tr><td>Mouse 42 Gemcitabine & Capecitabine Maximum tolerated dose</td><td>Died</td><td>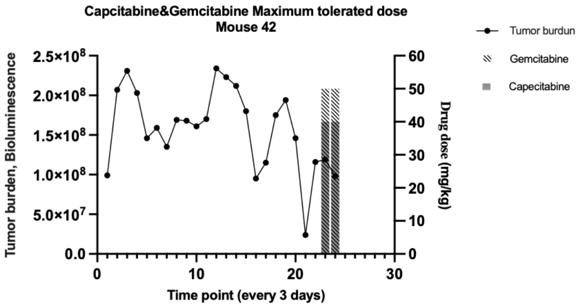
</td></tr>
<tr><td colspan="3">Gemcitabine & Capecitabine Ping-pong Intermittent</td></tr>
<tr><td>Mouse 43 Gemcitabine & Capecitabine Ping-pong Intermittent</td><td>Tumor size endpoint Caliper</td><td>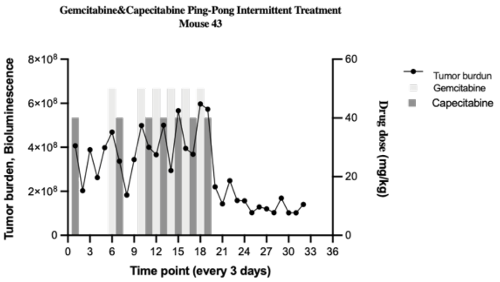
</td></tr>
</table>

| | | |
|---|---|---|
| **Mouse 44 Gemcitabine & Capecitabine Ping-pong Intermittent** | **Interfere with walking** | 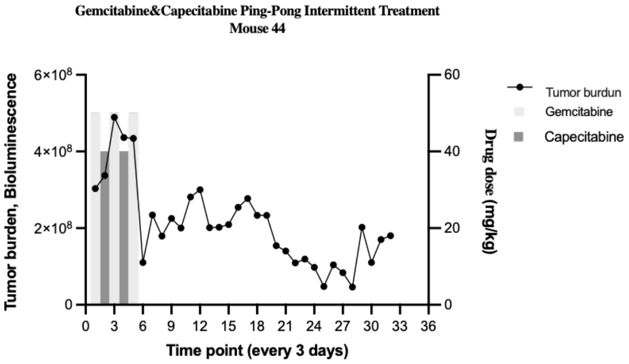 |
| **Mouse 45 Gemcitabine & Capecitabine Ping-pong Intermittent** | **Tumor size endpoint Caliper** | 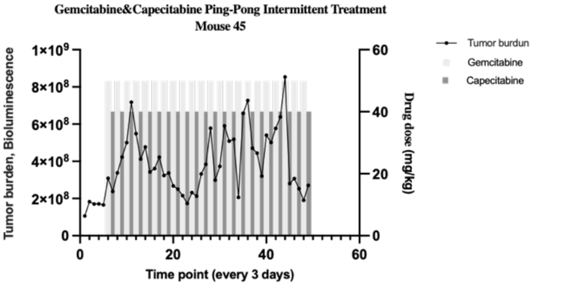 |
| **Mouse 46 Gemcitabine & Capecitabine Ping-pong Intermittent** | **Tumor size endpoint Caliper** | 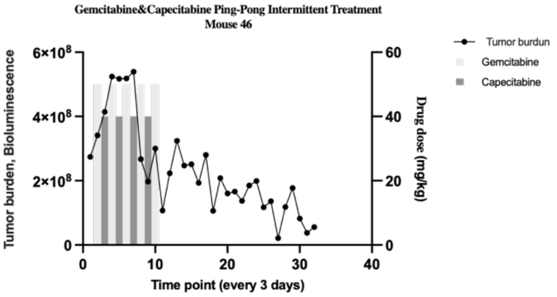 |
| **Mouse 47 Gemcitabine & Capecitabine Ping-pong Intermittent** | **Interfere with walking** | 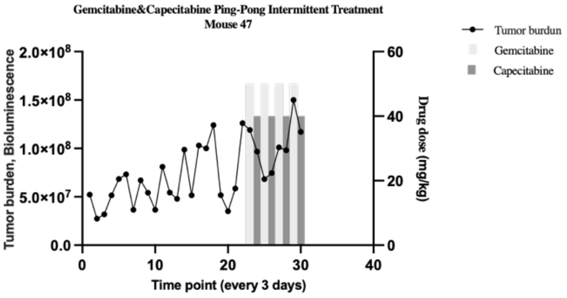 |

<table>
<tr>
<td>Mouse 48 Gemcitabine & Capecitabine Ping-pong Intermittent</td>
<td>Tumor size endpoint Caliper</td>
<td>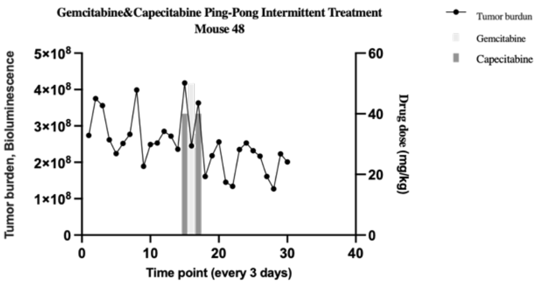
</td>
</tr>
<tr>
<td colspan="3">Gemcitabine & Capecitabine Tandem Intermittent</td>
</tr>
<tr>
<td>Mouse 49 Gemcitabine & Capecitabine Tandem Intermittent</td>
<td>weak and unhealthy</td>
<td>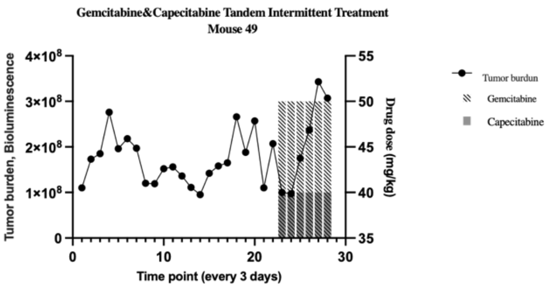
</td>
</tr>
<tr>
<td>Mouse 50 Gemcitabine & Capecitabine Tandem Intermittent</td>
<td>Tumor size interfere walking</td>
<td>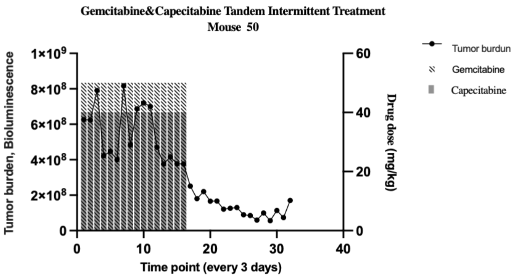
</td>
</tr>
<tr>
<td>Mouse 51 Gemcitabine & Capecitabine Tandem Intermittent</td>
<td>tumor size endpoint</td>
<td>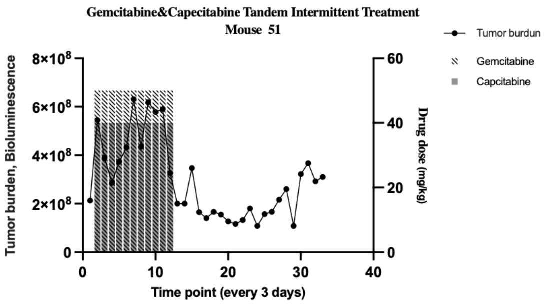
</td>
</tr>
</table>

<table>
<tr><td>Mouse 52 Gemcitabine & Capecitabine Tandem Intermittent</td><td>skinny and bleeding</td><td>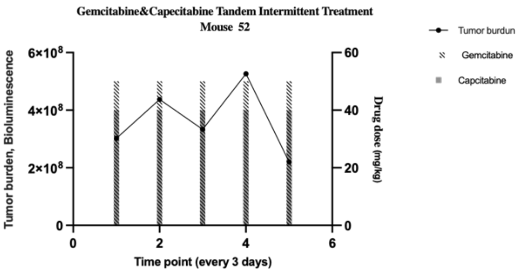
</td></tr>
<tr><td>Mouse 53 Gemcitabine & Capecitabine Tandem Intermittent</td><td>skinny</td><td>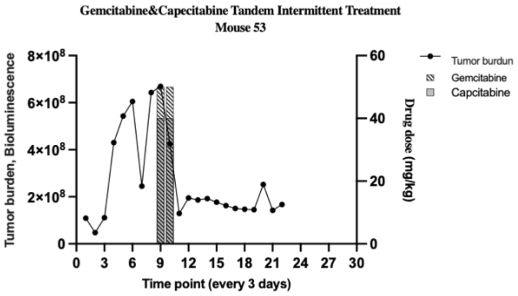
</td></tr>
<tr><td>Mouse 54 Gemcitabine & Capecitabine Tandem Intermittent</td><td>Died</td><td>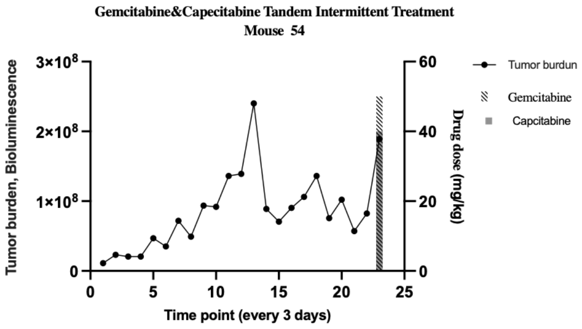
</td></tr>
<tr><td colspan="3">Gemcitabine & Capecitabine Ping-Pong Dose modulation</td></tr>
<tr><td>Mouse 55 Gemcitabine & Capecitabine Ping-Pong Dose modulation</td><td>tumor size interferes walking</td><td>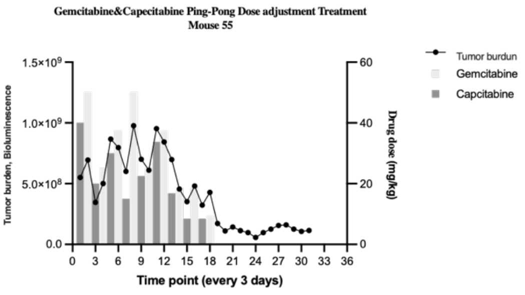
</td></tr>
</table>

| | | |
|---|---|---|
| **Mouse 56 Gemcitabine & Capecitabine Ping-Pong Dose modulation** | **unhealthy, skinny, invasive** | 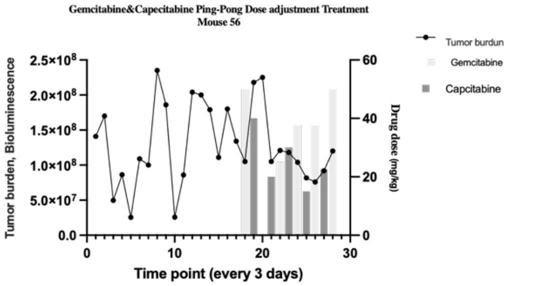  |
| **Mouse 57 Gemcitabine & Capecitabine Ping-Pong Dose modulation** | **tumor size interferes with walking** | 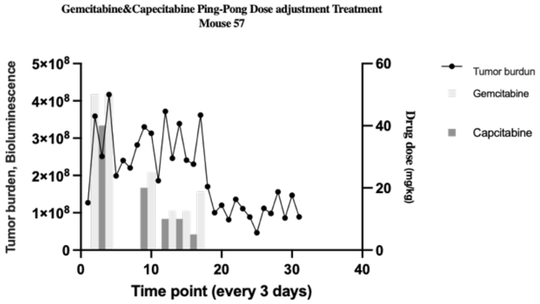  |
| **Mouse 58 Gemcitabine & Capecitabine Ping-Pong Dose modulation** | **metastasis underarm** | 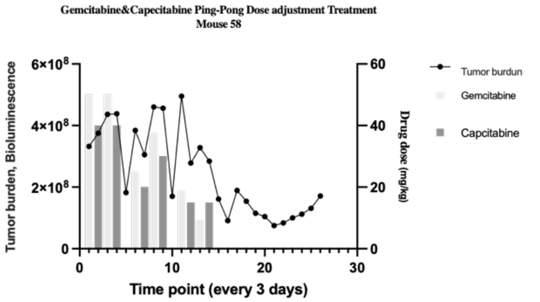  |
| **Mouse 59 Gemcitabine & Capecitabine Ping-Pong Dose modulation** | **tumor size interferes with walking** | 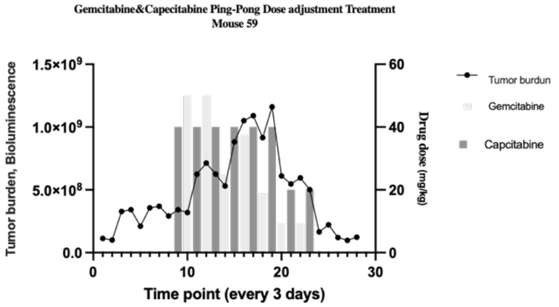  |

| | | |
|---|---|---|
| **Mouse 60 Gemcitabine & Capecitabine Ping-Pong Dose modulation** | **tumor size endpoint** | 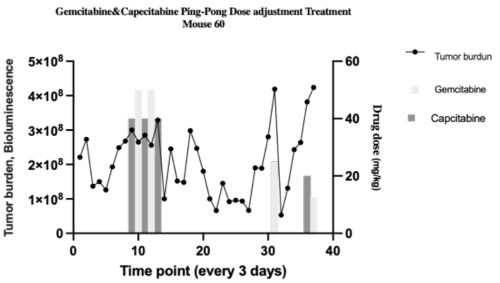 |
| **Gemcitabine & Capecitabine Tandem Dose modulation** | | |
| **Mouse 61 Gemcitabine & Capecitabine Tandem Dose modulation** | **Died during surgery for implanting 2nd time estrogen** | 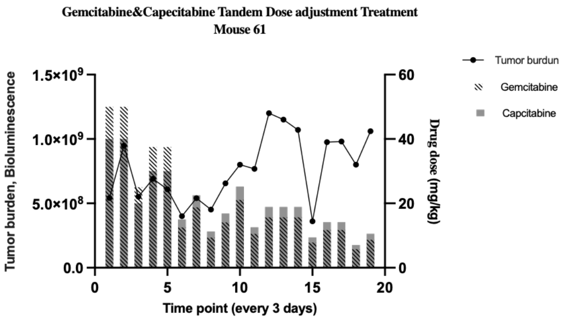 |
| **Mouse 62 Gemcitabine & Capecitabine Tandem Dose modulation** | **metastasis underarm** | 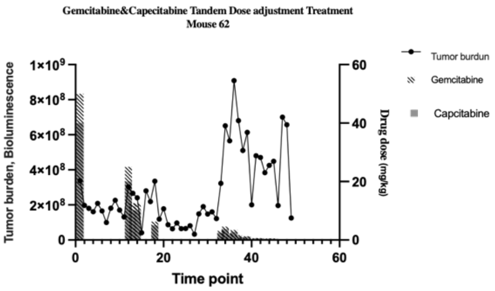 |
| **Mouse 63 Gemcitabine & Capecitabine Tandem Dose modulation** | **tumor invasive** | 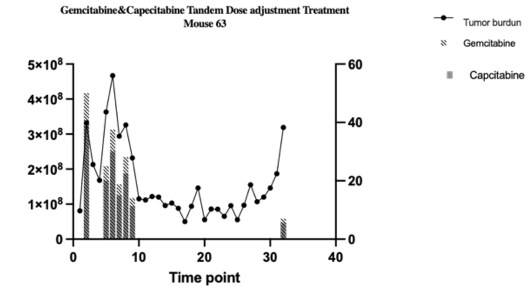 |

| | | |
|---|---|---|
| **Mouse 64 Gemcitabine & Capecitabine Tandem Dose modulation** | **tumor size endpoint** | 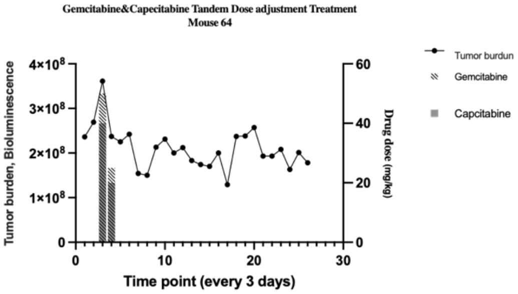 |
| **Mouse 65 Gemcitabine & Capecitabine Tandem Dose modulation** | **interfere with walking** | 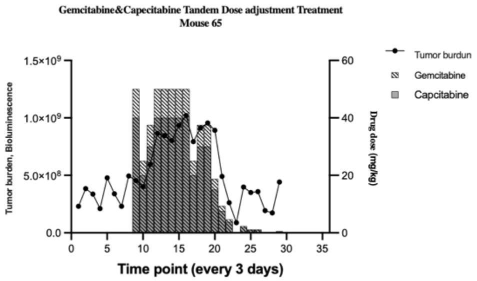 |
| **Mouse 66 Gemcitabine & Capecitabine Tandem Dose modulation** | **Died** | 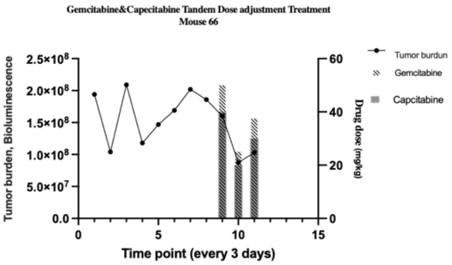 |
| **Control no treatment** | | |
| **Mouse 67 Control no treatment** | **skinny** | 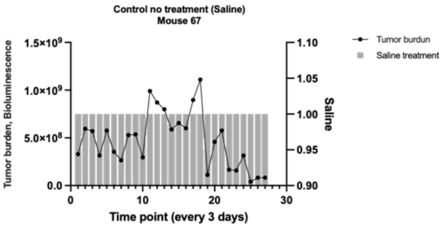 |

| | | |
|---|---|---|
| **Mouse 68 Control no treatment** | **metastasis underarm** | 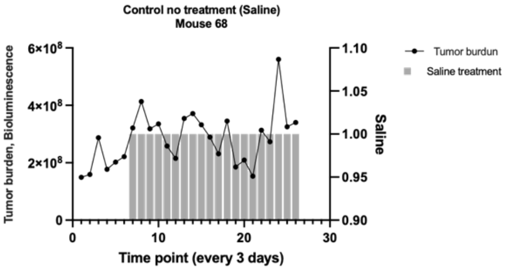 |
| **Mouse 69 Control no treatment** | **skinny and hair loss and tumor size** | 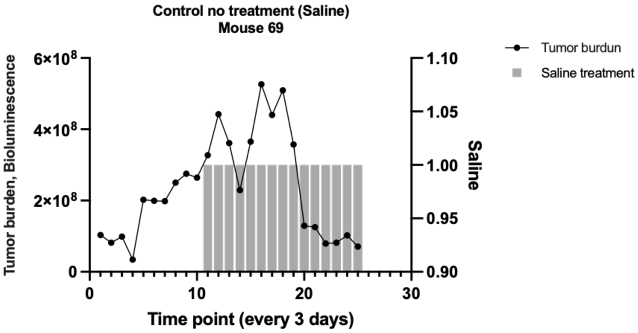 |
| **Mouse 70 Control no treatment** | **tumor size end point caliper** | 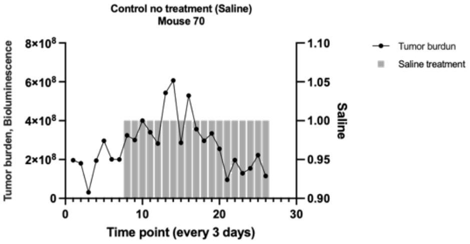 |

**Table S2: Drug Dose applied in each group of treatment.**

| Group of treatment | Average of cumulative dose received | Average dose per day |
|---|---|---|
| Gemcitabine Dose Modulation | GEM: 256 mg/kg | GEM: 5.8 mg/kg |
| Gemcitabine Intermittent | GEM: 33.3 mg/kg | GEM: 5.5 mg/kg |
| Gemcitabine Maximum Tolerated Dose | GEM: 783.3 mg/kg | GEM: 12.8 mg/kg |
| Capecitabine Dose Modulation | CAP: 533.8 mg/kg | CAP: 11.8 mg/kg |
| Capecitabine Intermittent | CAP:1226.6 mg/kg | CAP: 26 mg/kg |
| Capecitabine Maximum Tolerated Dose | CAP: 1373.3 mg/kg | CAP: 28.4 mg/kg |
| Capecitabine + Gemcitabine Ping-Pong Dose Modulation | GEM: 379.1 mg/kg<br>CAP:475.5 mg/kg | GEM: 4.9 mg/kg<br>CAP: 6.1 mg/kg |
| Capecitabine + Gemcitabine in Tandem Dose Modulation | GEM: 229 mg/kg<br>CAP:492.1 mg/kg | GEM: 4.3 mg/kg<br>CAP: 8.7 mg/kg |
| Capecitabine + Gemcitabine Ping-Pong Intermittent | GEM: 550 mg/kg<br>CAP:660 mg/kg | GEM: 6.4 mg/kg<br>CAP: 8 mg/kg |
| Capecitabine + Gemcitabine in Tandem Intermittent | GEM: 341.6 mg/kg<br>CAP: 606.6 mg/kg | GEM: 9 mg/kg<br>CAP:12.5 mg/kg |
| Capecitabine + Gemcitabine Maximum Tolerated Dose | GEM: 433.3 mg/kg<br>CAP: 766.6 mg/kg | GEM: 20.6 mg/kg<br>CAP: 30.6 mg/kg |

### Table S3: Immunohistochemistry Results

| Number Of Mouse | Percentage Of Ki-67 Positive | Figure Of Detected Section |
|---|---|---|
| Control (No Treatment) | | |
| #68 Control (No Treatment) | 8.7% | 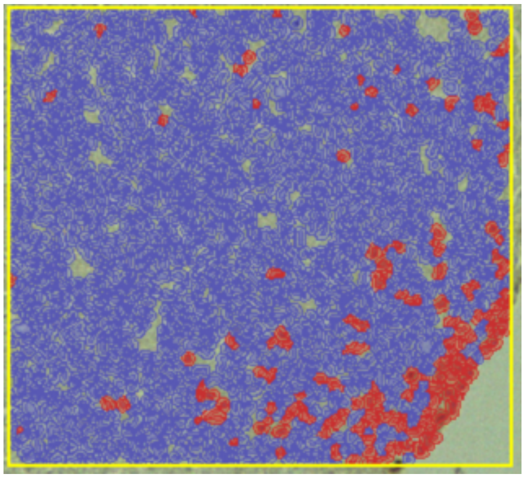 |
| #69 CONTROL NO TREATMENT | 46.2% | 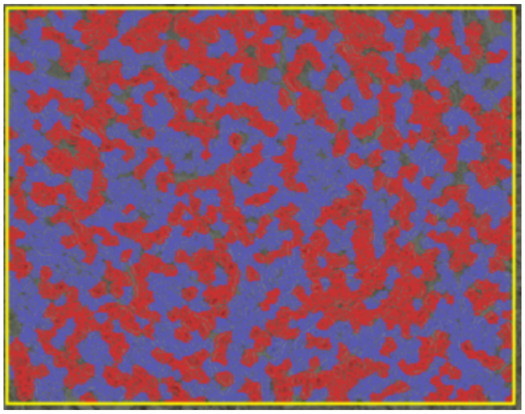 |

| | | |
|---|---|---|
| **#67 Control (No Treatment)** | **4.37%** | 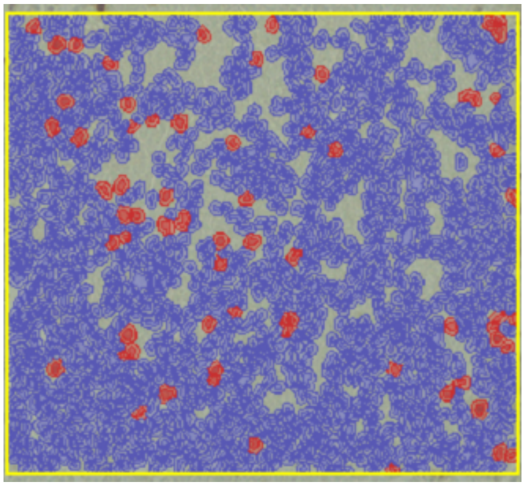 |
| **#70 Control (No Treatment)** | **23.9%** | 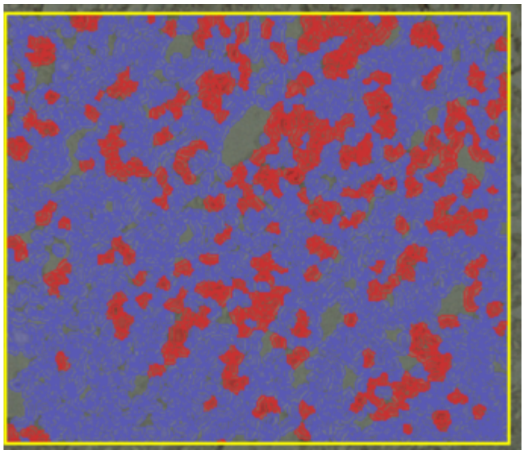 |
| **Capecitabine** | | |
| **#20 Capecitabine Maximum Tolerated Dose** | **21.1%** | 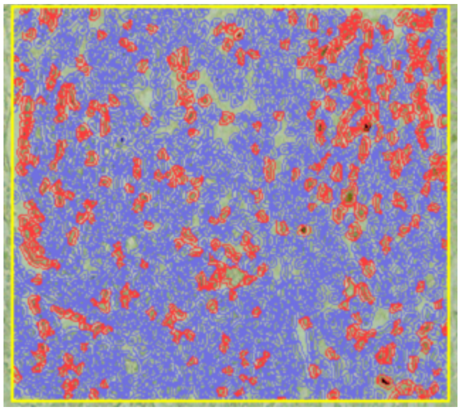 |

| | | |
|---|---|---|
| **#22 Capecitabine Maximum Tolerated Dose** | **16.4%** | 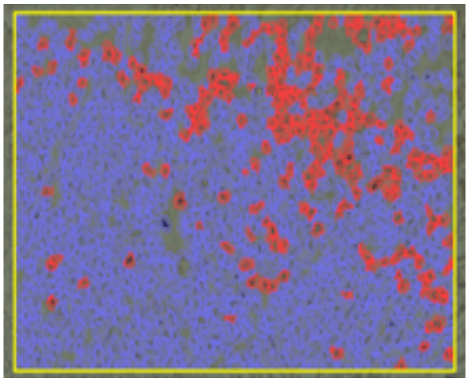 |
| **#23 Capecitabine Maximum Tolerated Dose** | **21.7%** | 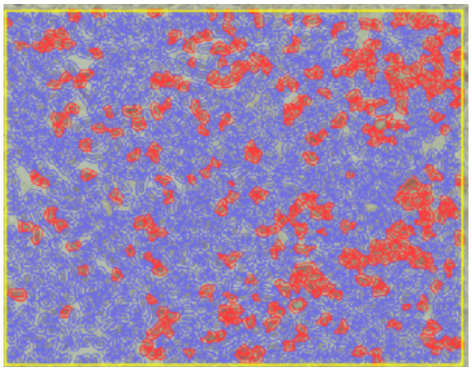 |
| **#34 Capecitabine Dose Modulation** | **2%** | 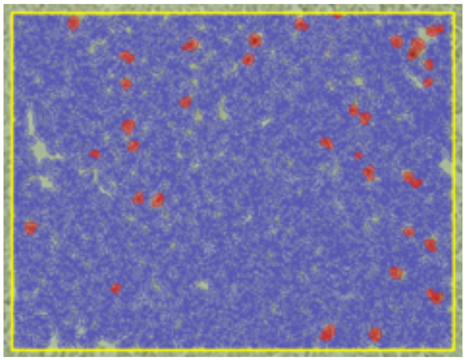 |
| **#35 Capecitabine Dose Modulation** | **2.48%** | 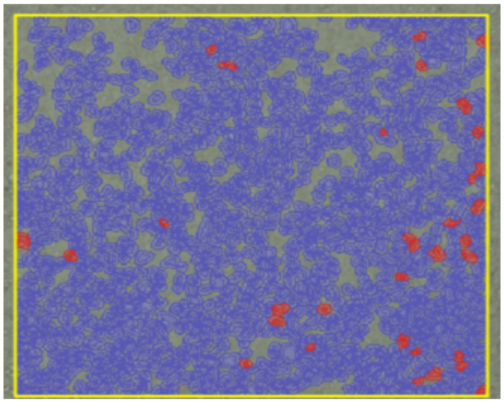 |

| | | |
|---|---|---|
| **#26**<br>**Capecitabine Intermittent** | **13.8%** | 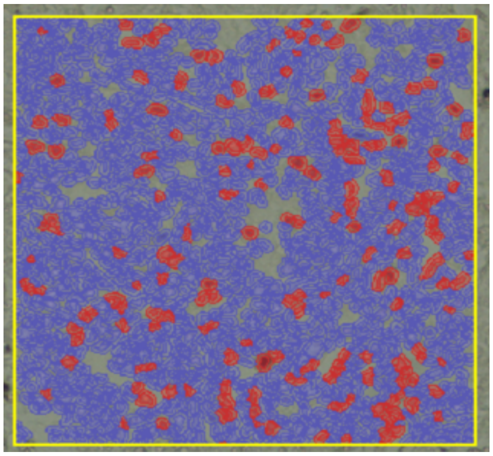 |
| **#25**<br>**Capecitabine Intermittent** | **5.97%** | 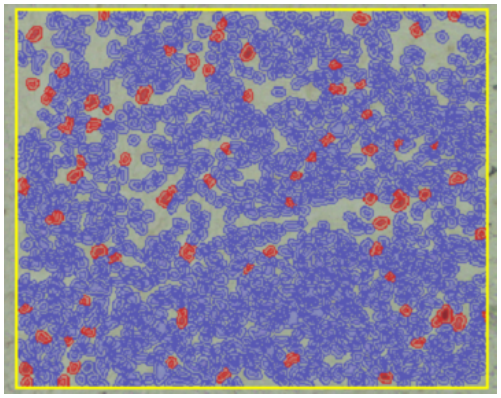 |
| **#29**<br>**Capecitabine Intermittent** | **19.8%** | 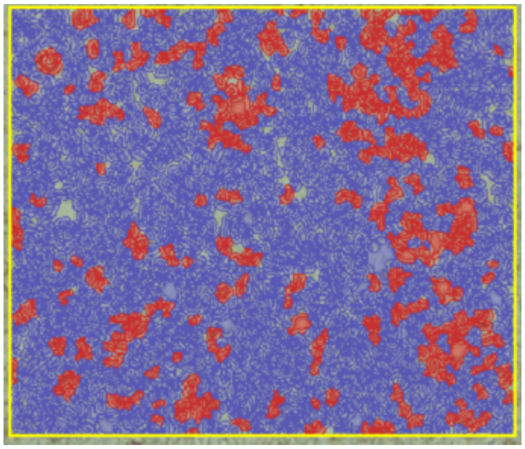 |
| | **Gemcitabine** | |

| | | |
|---|---|---|
| **#1 Gemcitabine Maximum Tolerated Dose** | **17.9%** | 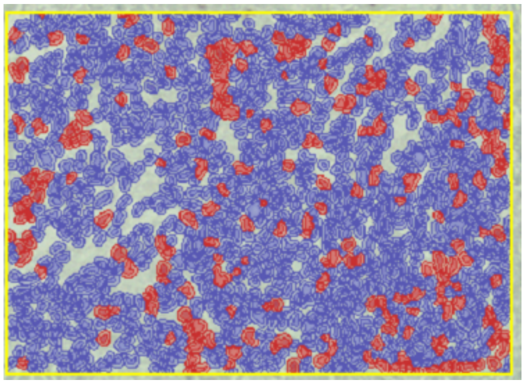 |
| **#2 Gemcitabine Maximum Tolerated Dose** | **26.8%** | 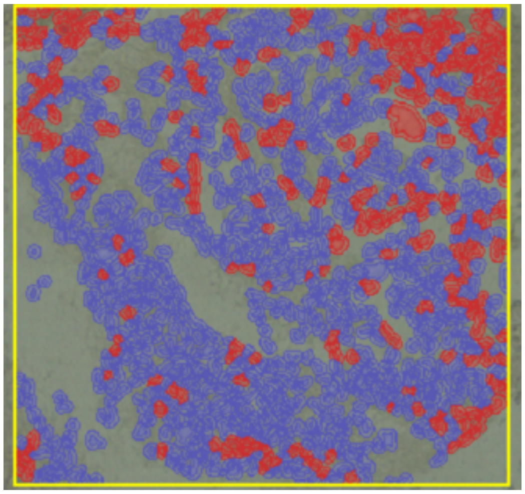 |
| **#6 Gemcitabine Maximum Tolerated Dose** | **7.93%** | 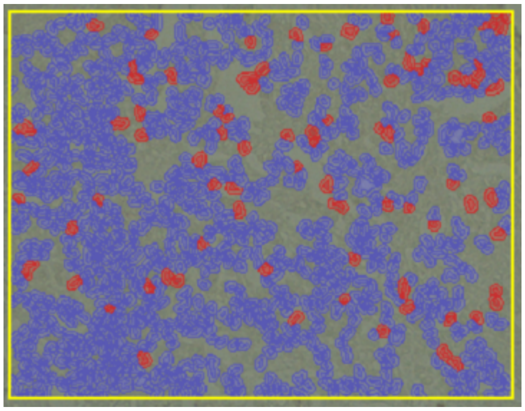 |

| | | |
|---|---|---|
| **#13 Gemcitabine Dose Modulation** | **10.5%** | 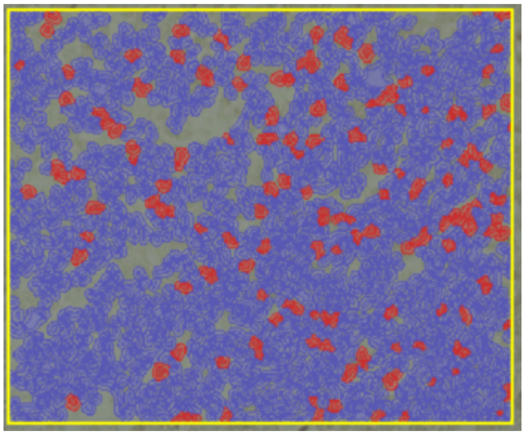 |
| **#15 Gemcitabine Dose Modulation** | **7.95%** | 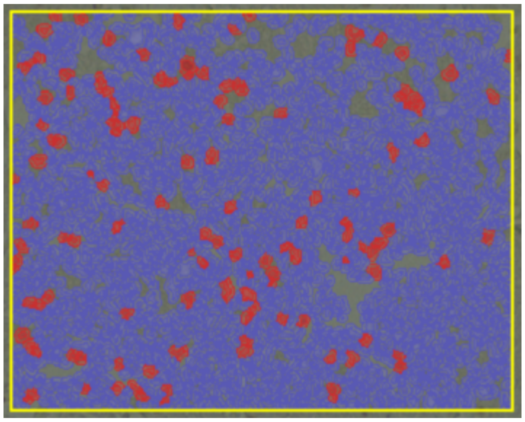 |
| **#16 Gemcitabine Dose Modulation** | **3.46%** | 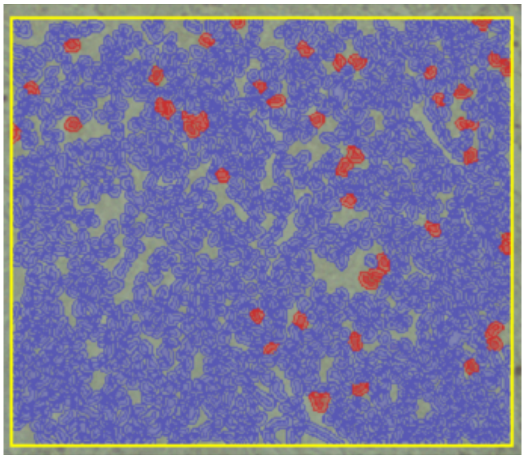 |

| | | |
|---|---|---|
| **#17**<br>**Gemcitabine**<br>**Dose Modulation** | **14.6%** | 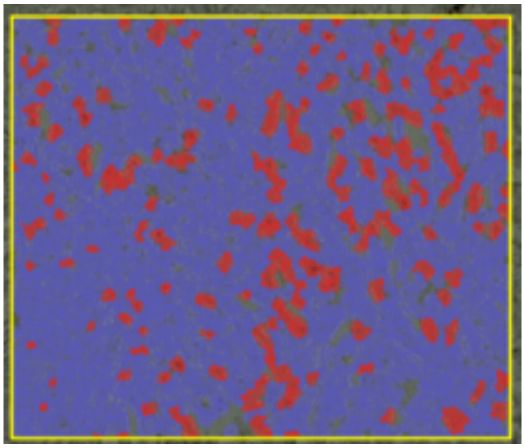 |
| **#18**<br>**Gemcitabine**<br>**Dose Modulation** | **2.9%** | 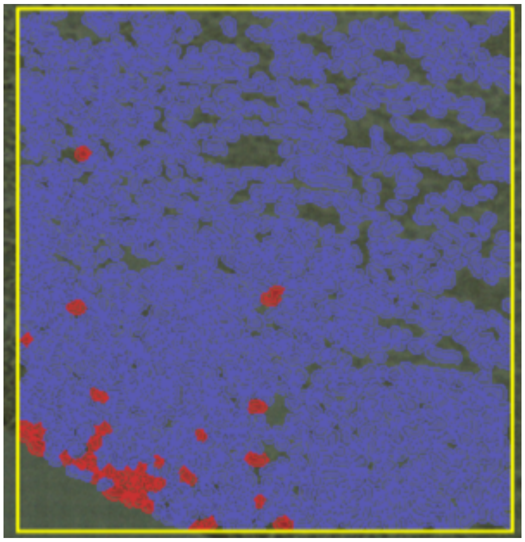 |
| **#14**<br>**Gemcitabine**<br>**Dose Modulation** | **14.9** | 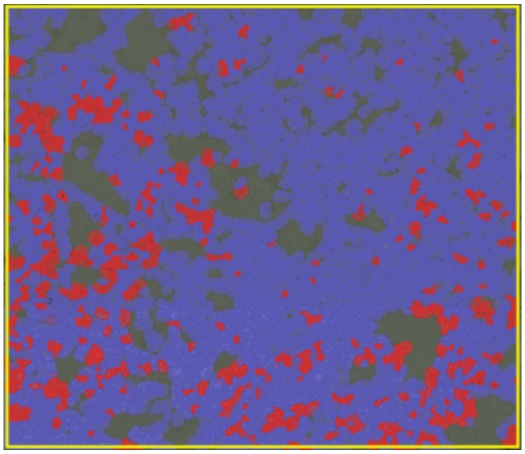 |

| | | |
|---|---|---|
| **#8**<br>**Gemcitabine Intermittent** | **16.9%** | 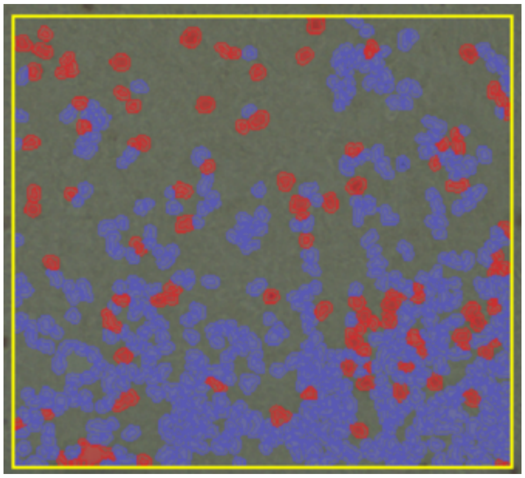 |
| **#11**<br>**Gemcitabine Intermittent** | **34%** | 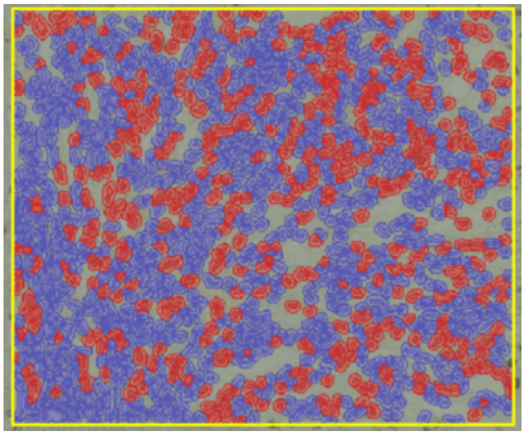 |
| **#10**<br>**Gemcitabine Intermittent** | **6.3%** | 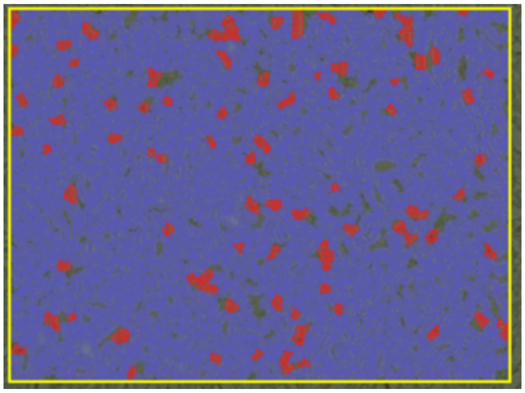 |

| | | |
|---|---|---|
| **#7**<br>**Gemcitabine Intermittent** | **18.4%** | 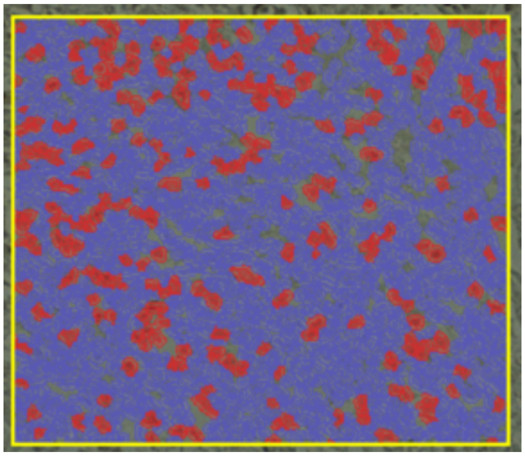 |
| **#9**<br>**Gemcitabine Intermittent** | **2.13%** | 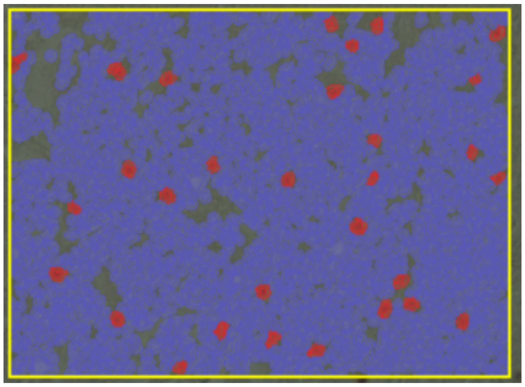 |
| **Capecitabine & Gemcitabine** | | |
| **#38**<br>**Gemcitabine & Capecitabine**<br>**Maximum Tolerated Dose** | **25%** | 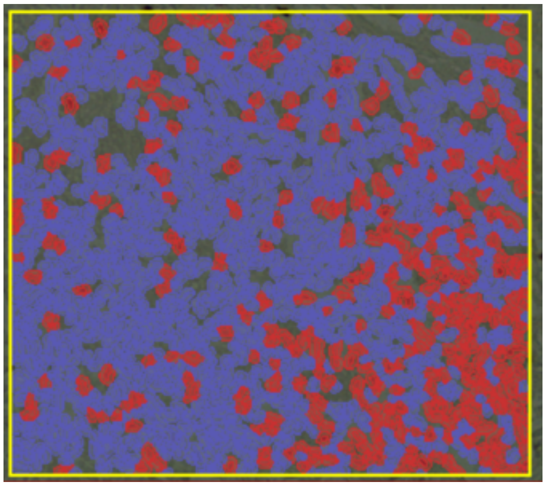 |

| | | |
|---|---|---|
| **#58**<br>**Gemcitabine & Capecitabine Ping-Pong Dose Modulation** | **3.63%** | 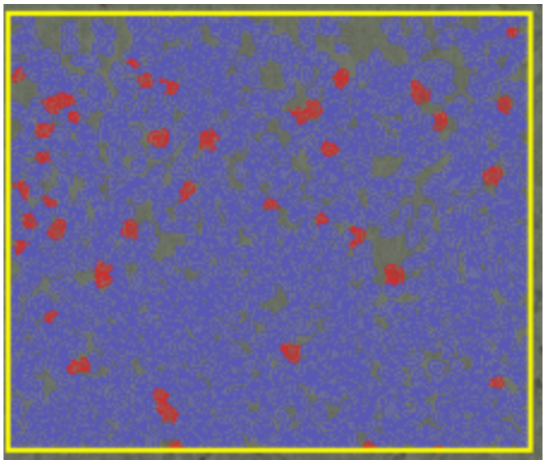 |
| **#55**<br>**Gemcitabine & Capecitabine Ping-Pong Dose Modulation** | **2.8%** | 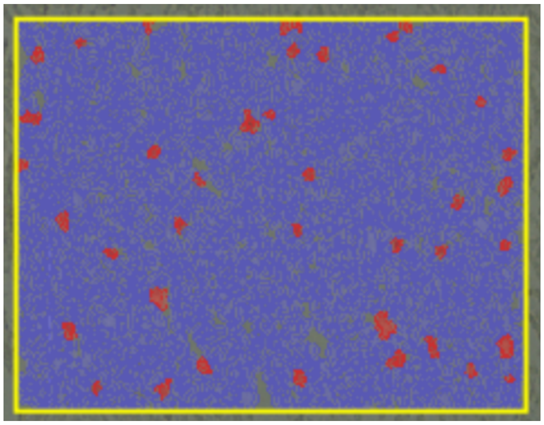 |
| **#56**<br>**Gemcitabine & Capecitabine Ping-Pong Dose Modulation** | **5.3%** | 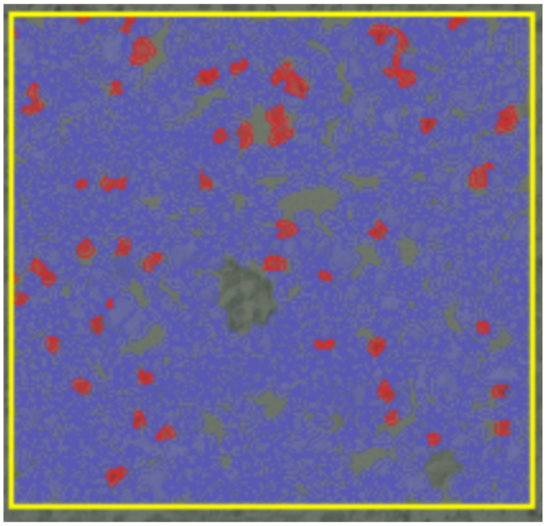 |
| **Gemcitabine & Capecitabine Ping-Pong Intermittent** | | |

| | | |
|---|---|---|
| **#46**<br>**Gemcitabine & Capecitabine Ping-Pong Intermittent** | **5.28%** | 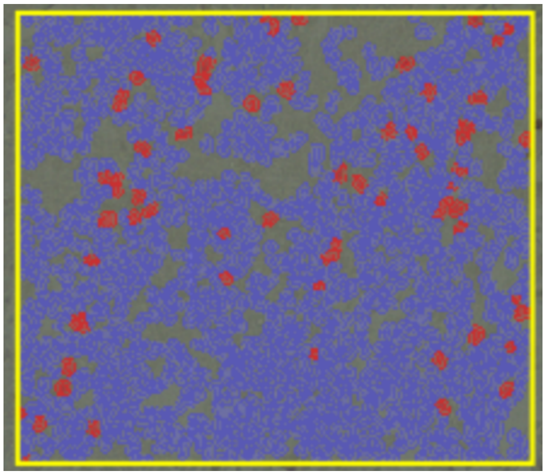 |
| **#44**<br>**Gemcitabine & Capecitabine Ping-Pong Intermittent** | **6.06%** | 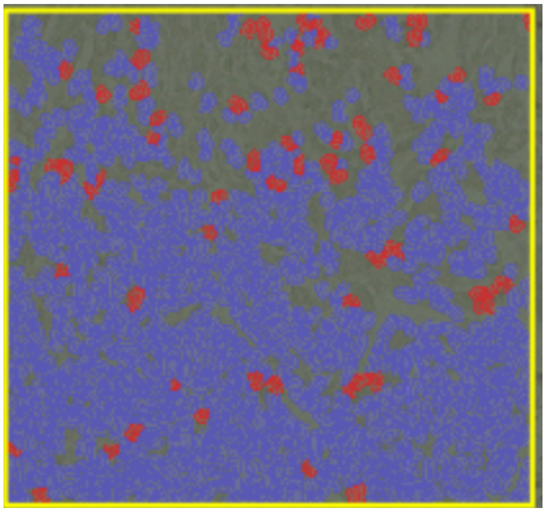 |
| **#48**<br>**Gemcitabine & Capecitabine Ping-Pong Intermittent** | **3%** | 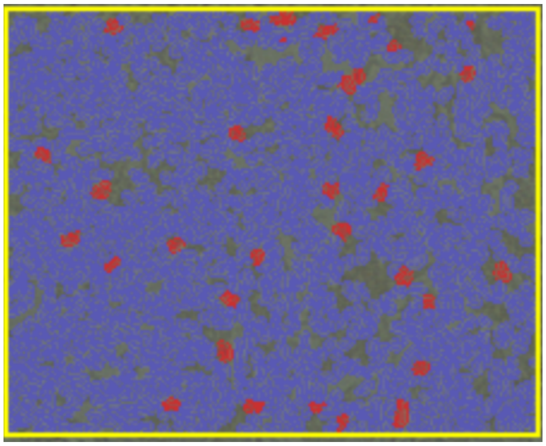 |

| | | |
|---|---|---|
| **#47**<br>**Gemcitabine & Capecitabine**<br>**Ping-Pong Intermittent** | **7.9%** | 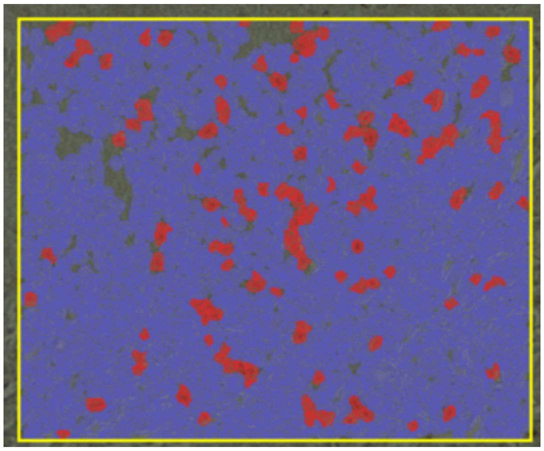 |
| **#45**<br>**Gemcitabine & Capecitabine**<br>**Ping-Pong Intermittent** | **6.77%** | 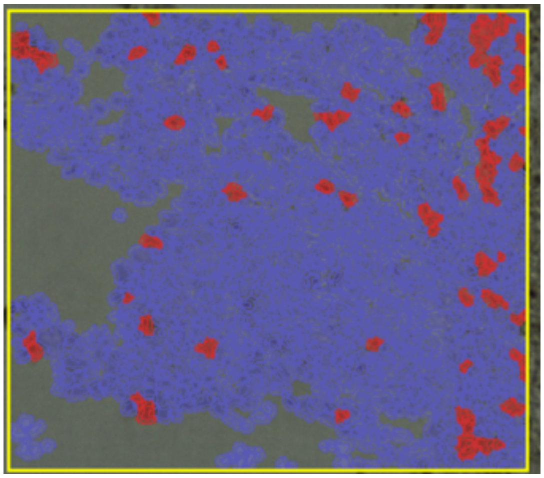 |
| **#63**<br>**Gemcitabine & Capecitabine**<br>**Tandem Dose Modulation** | **3.38%** | 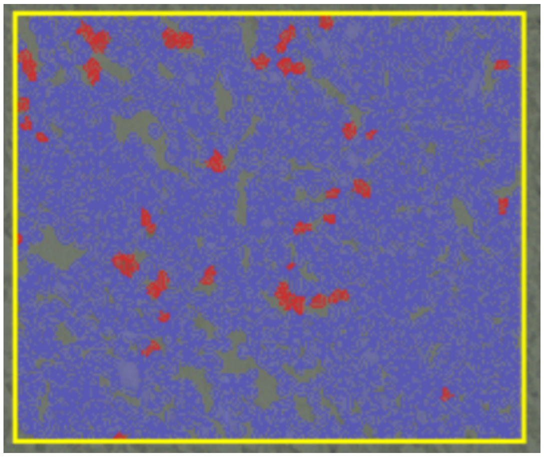 |

| | | |
|---|---|---|
| **#64**<br>**Gemcitabine & Capecitabine Tandem Dose Modulation** | **1.6%** | 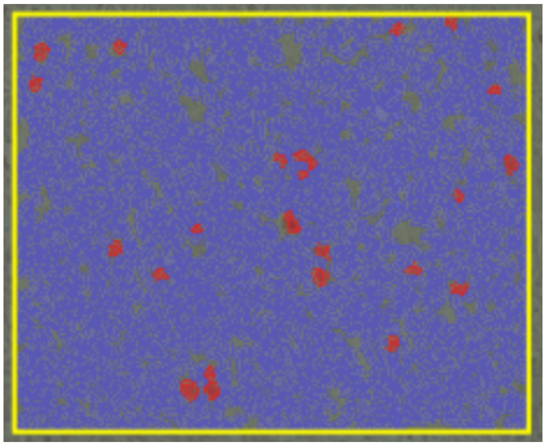 |
| **#65**<br>**Gemcitabine & Capecitabine Tandem Dose Modulation** | **12.4%** | 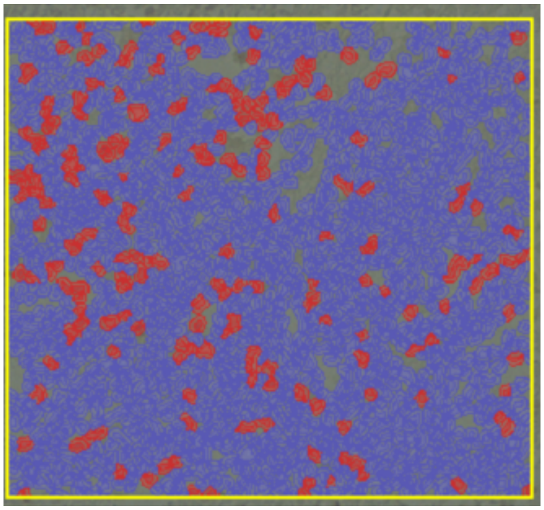 |
| **#62**<br>**Gemcitabine & Capecitabine Tandem Dose Modulation** | **11.59%** | 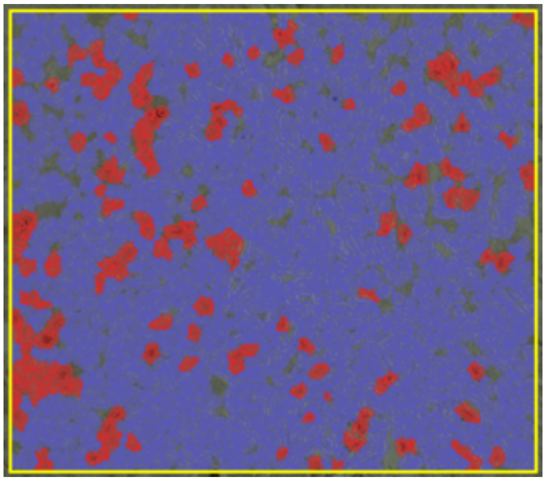 |

| | | |
|---|---|---|
| **#51**<br>**Gemcitabine & Capecitabine**<br>**Tandem Intermittent** | **3.2%** | 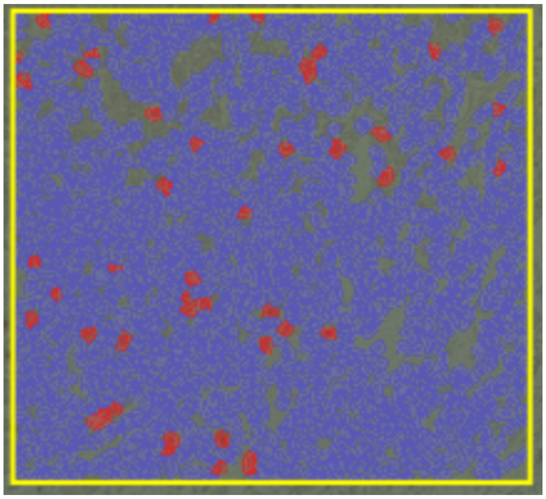 |
| **#52**<br>**Gemcitabine & Capecitabine**<br>**Tandem Intermittent** | **18.5%** | 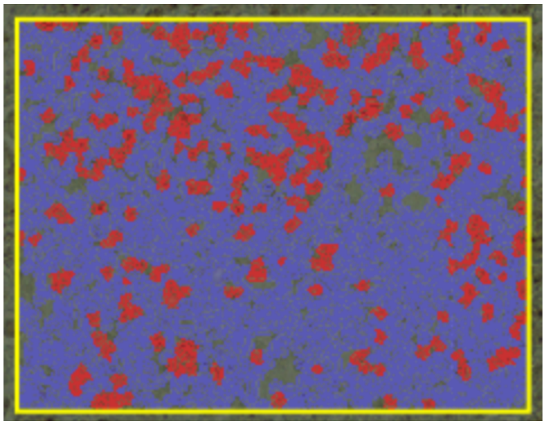 |
| **#50**<br>**Gemcitabine & Capecitabine**<br>**Tandem Intermittent** | **5.02%** | 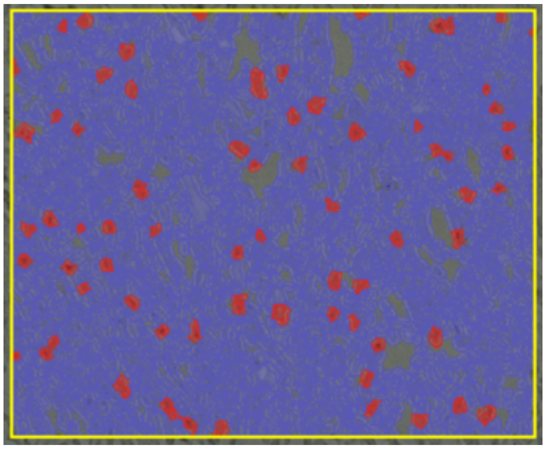 |

| | | |
|---|---|---|
| **#49**<br>**Gemcitabine & Capecitabine Tandem Intermittent** | **2.36%** | 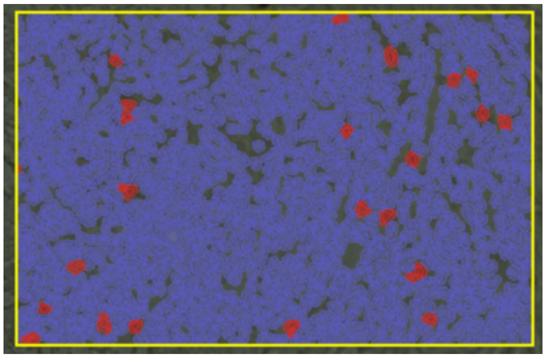 |

| **Number Of Mouse** | **Percentage Of Caspase-3 Positive** | **Figure Of Detected Section** |
|---|---|---|
| **Control (No Treatment)** | | |
| **Control (No Treatment)** | **1.8** | 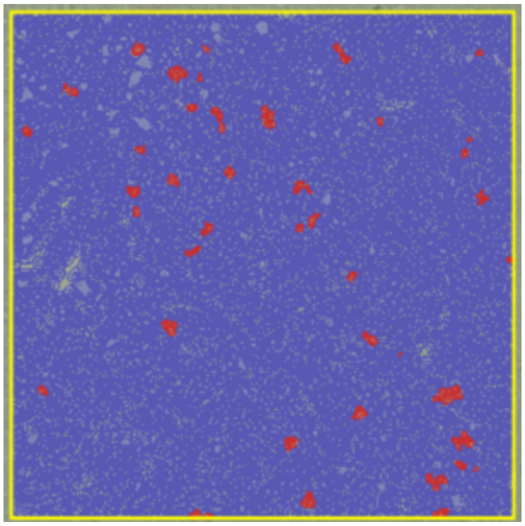 |

| | | |
|---|---|---|
| **Control (No Treatment)** | **9.74** | 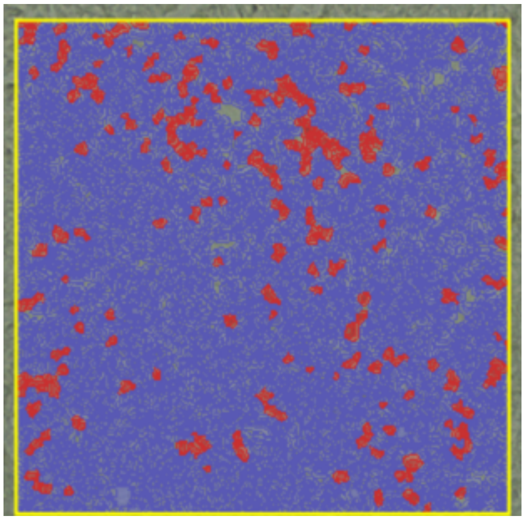 |
| **Control (No Treatment)** | **1.78** | 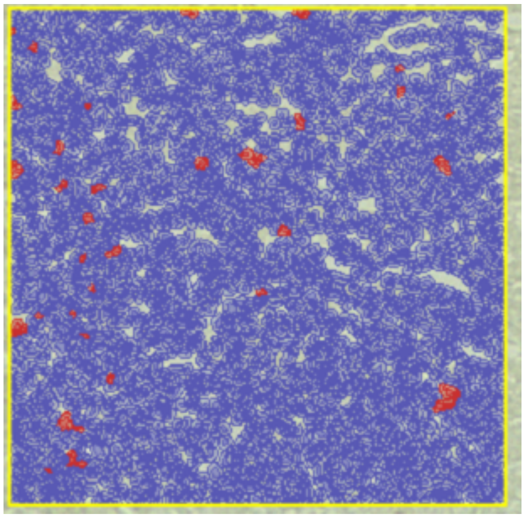 |
| **Control (No Treatment)** | **8.8** | 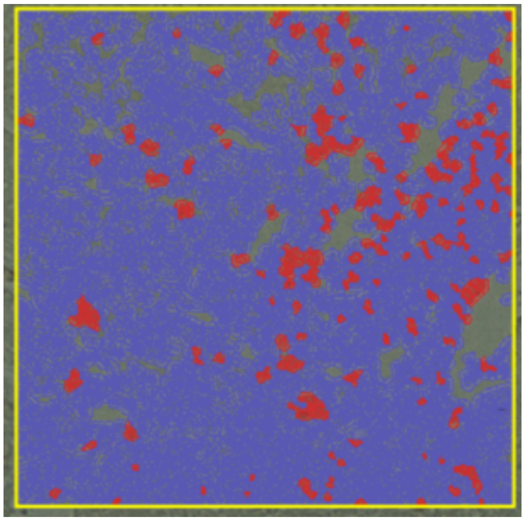 |

| **Capecitabine** | | |
|---|---|---|
| **Capecitabine Maximum Tolerated Dose** | **5.9** | 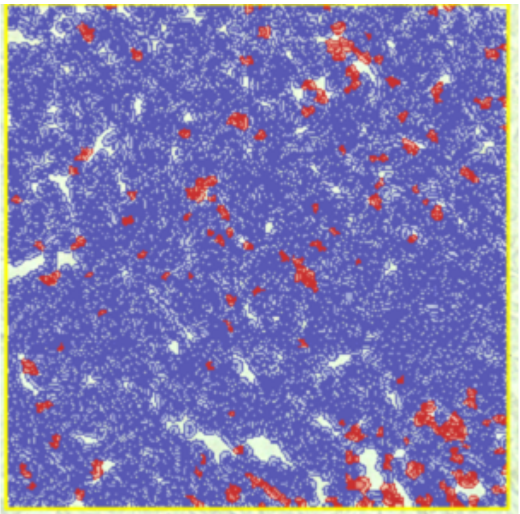 |
| **Capecitabine Maximum Tolerated Dose** | **7** | 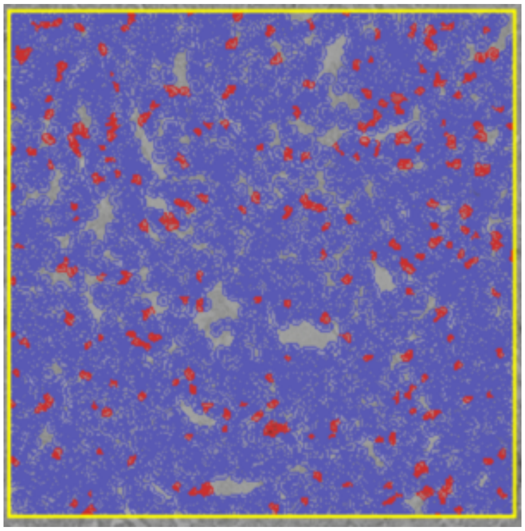 |

| | | |
|---|---|---|
| **Capecitabine Maximum Tolerated Dose** | **0.4** | 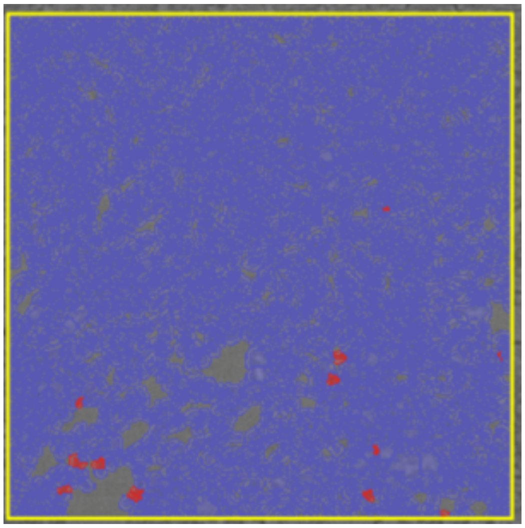 |
| **Capecitabine Dose Modulation** | **4.3** | 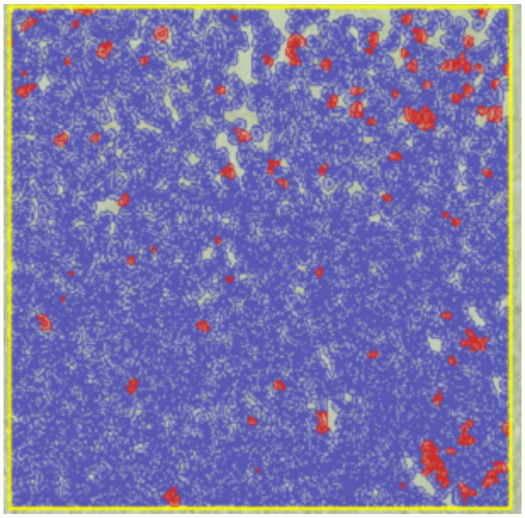 |
| **Capecitabine Dose Modulation** | **21.8** | 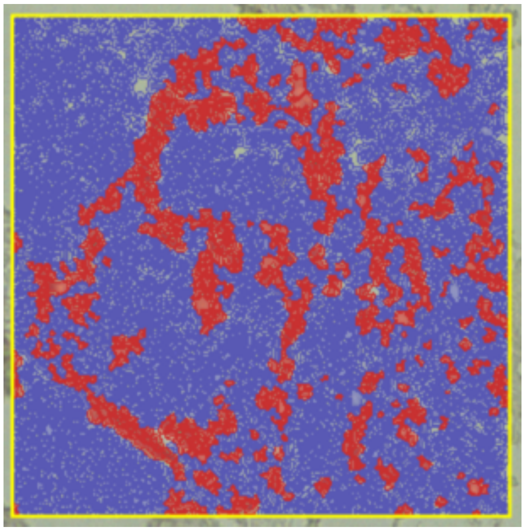 |

| | | |
|---|---|---|
| **Capecitabine Dose Modulation** | **12.7** | 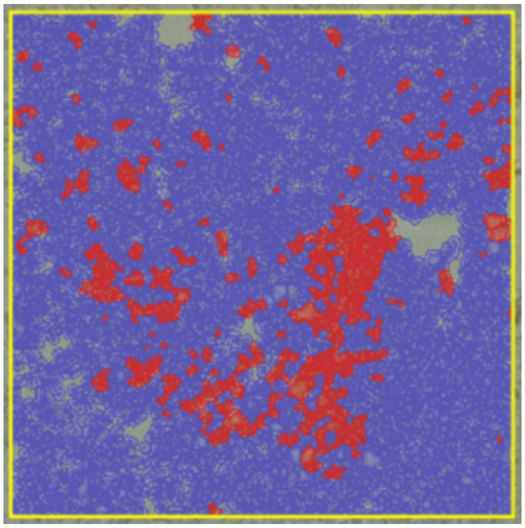 |
| **Capecitabine Intermittent** | **10.3** | 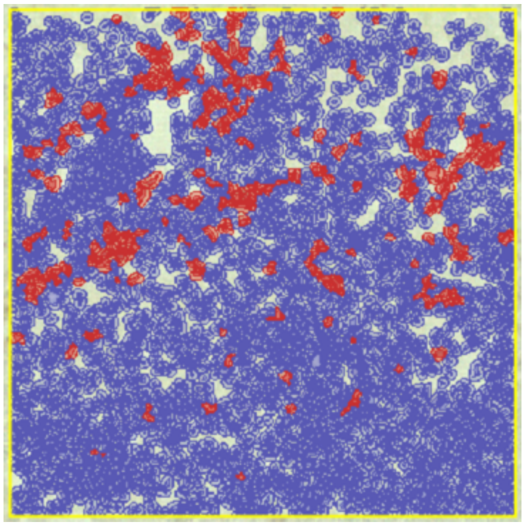 |
| **Capecitabine Intermittent** | **1.36** | 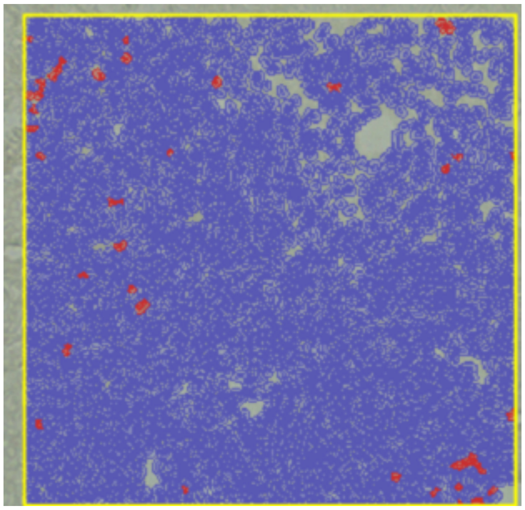 |

| | | |
|---|---|---|
| **Capecitabine Intermittent** | **8.56** | 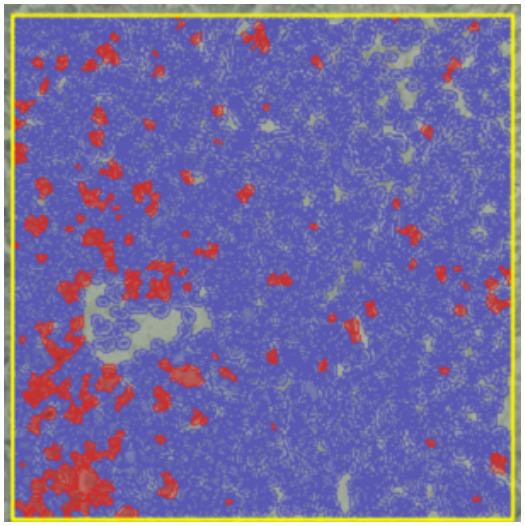 |
| **Gemcitabine** | | |
| **Gemcitabine Maximum Tolerated Dose** | **3.94** | 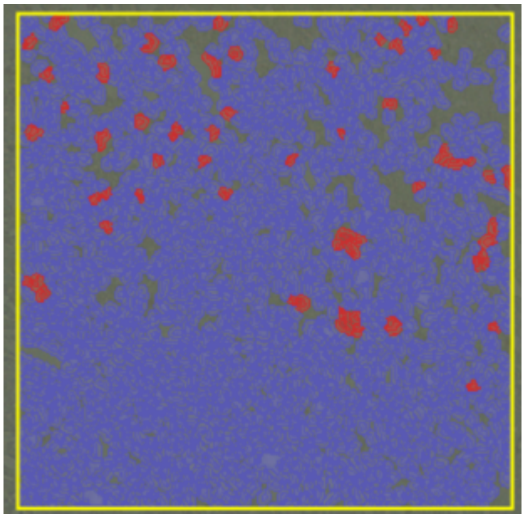 |

| | | |
|---|---|---|
| **Gemcitabine Maximum Tolerated Dose** | **2.7** | 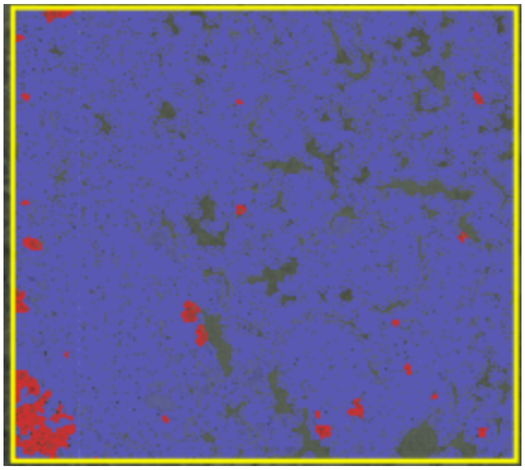 |
| **Gemcitabine Maximum Tolerated Dose** | **4.19** | 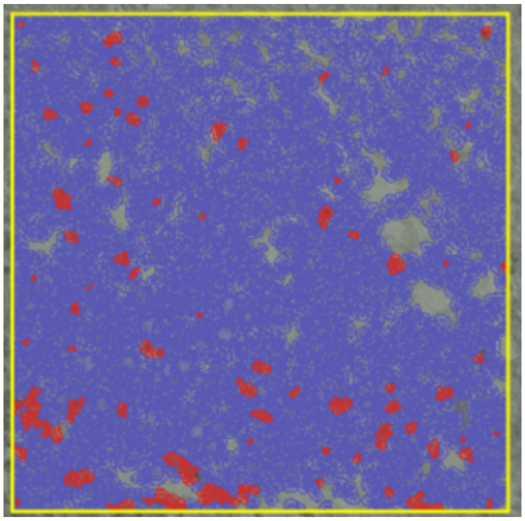 |
| **Gemcitabine Dose Modulation** | **1.6** | 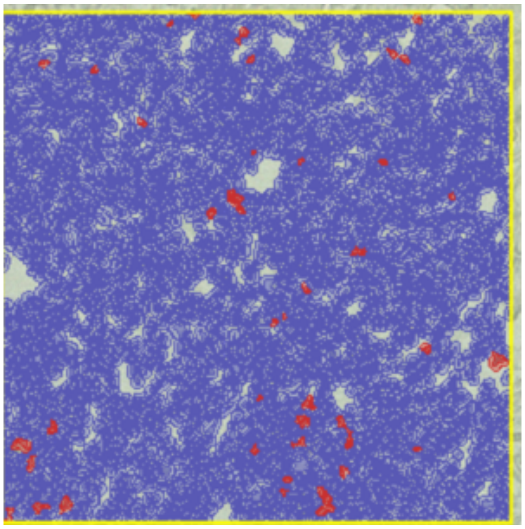 |

| | | |
|---|---|---|
| **Gemcitabine Dose Modulation** | **7.02** | 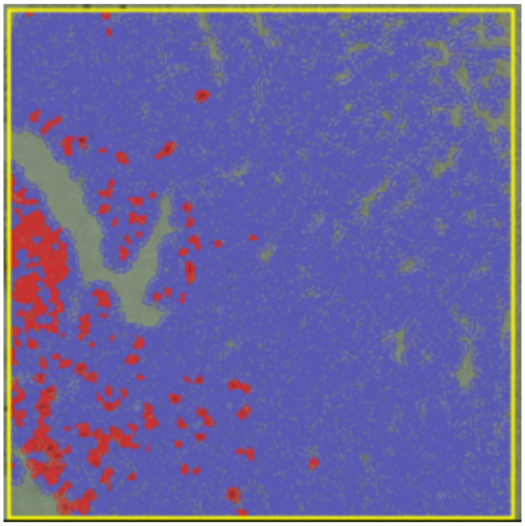 |
| **Gemcitabine Dose Modulation** | **2.9** | 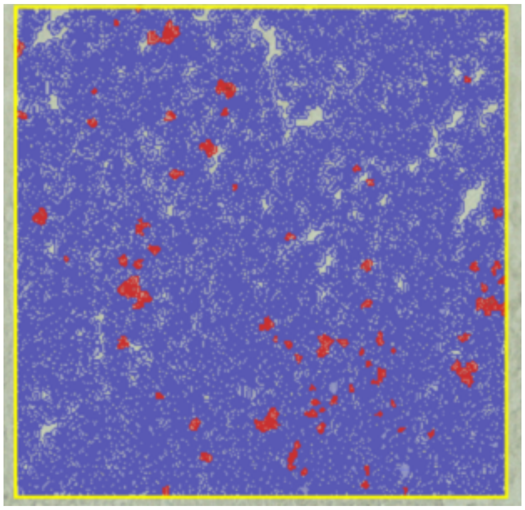 |
| **Gemcitabine Dose Modulation** | **11.6** | 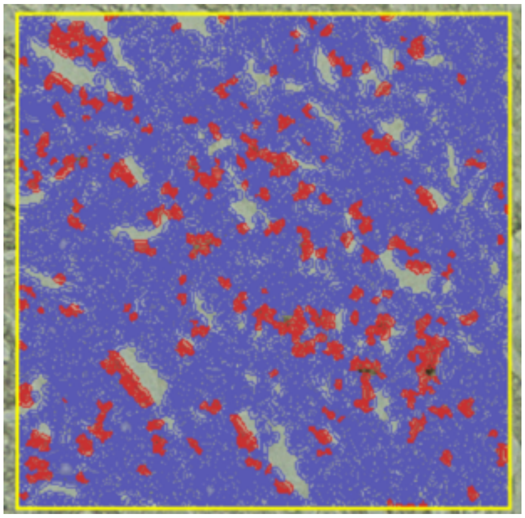 |

| | | |
|---|---|---|
| **Gemcitabine Dose Modulation** | **41.7** | 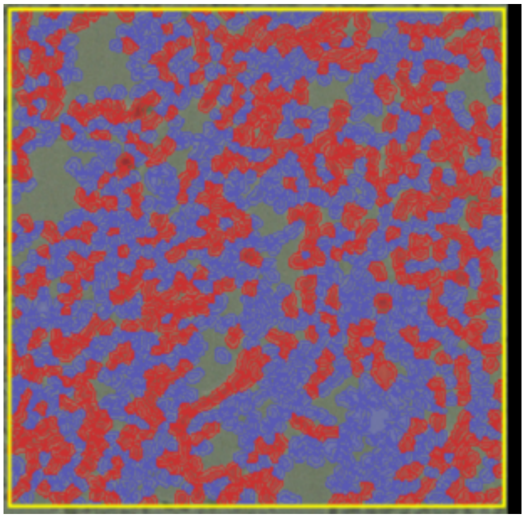 |
| **Gemcitabine Dose Modulation** | **14.1** | 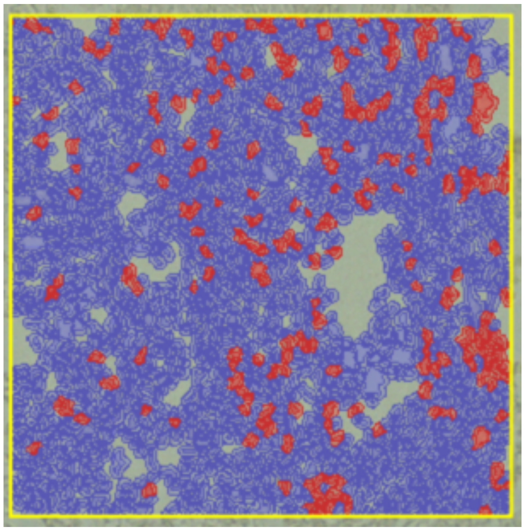 |

| | | |
|---|---|---|
| **Gemcitabine Intermittent** | **8.7** | 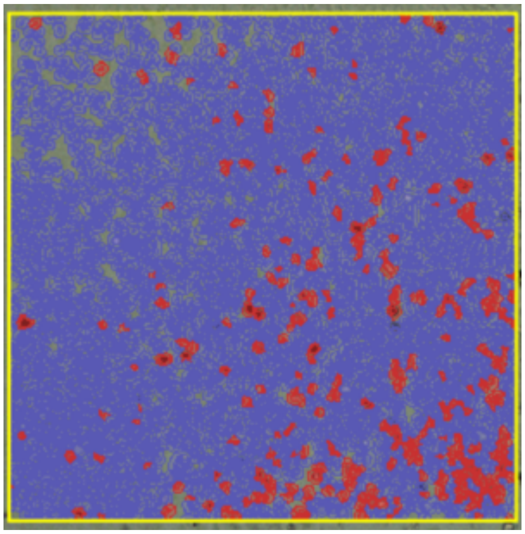 |
| **Gemcitabine Intermittent** | **3.8** | 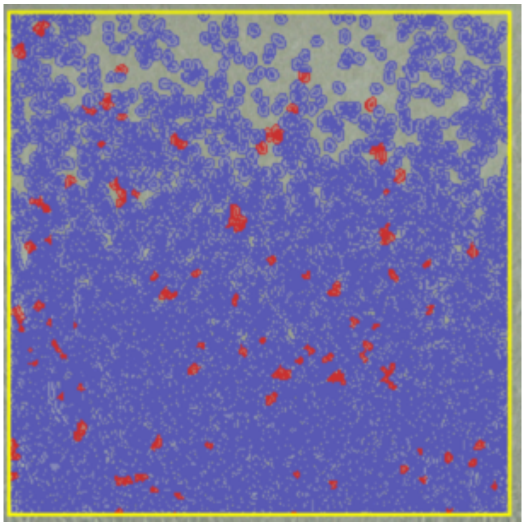 |
| **Gemcitabine Intermittent** | **5.6** | 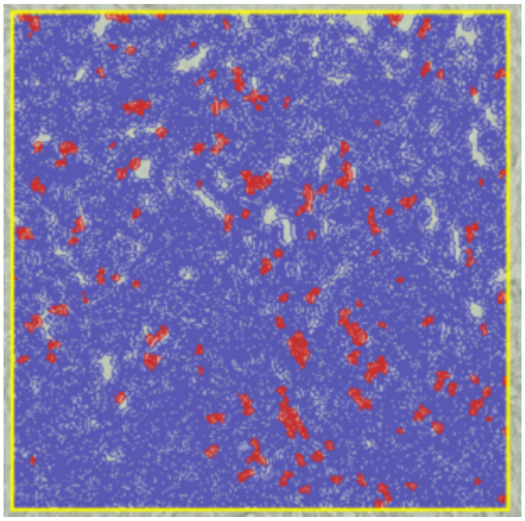 |

| | | |
|---|---|---|
| **Gemcitabine Intermittent** | **10.3** | 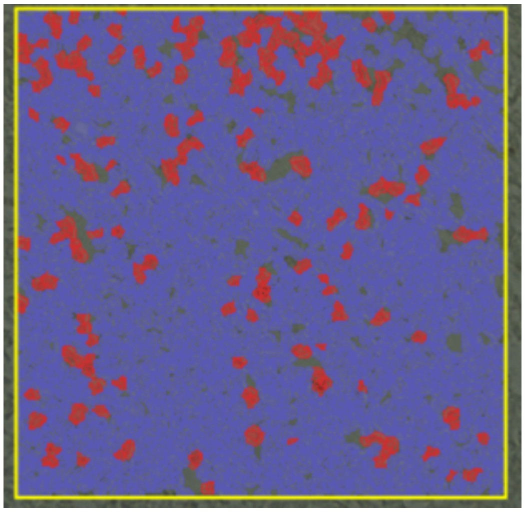 |
| **Gemcitabine Intermittent** | **4.16** | 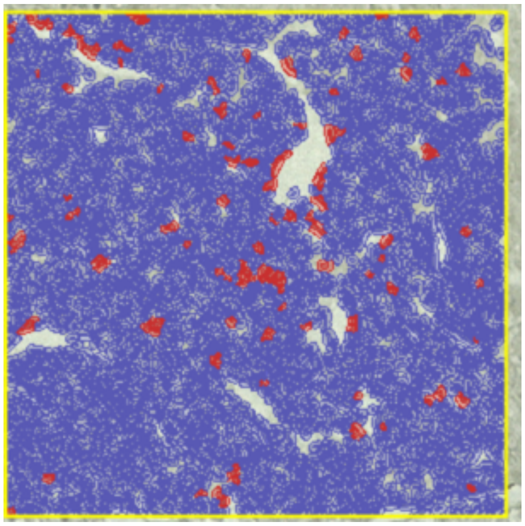 |
| **Capecitabine & Gemcitabine** | | |

| | | |
|---|---|---|
| **Gemcitabine & Capecitabine Maximum Tolerated Dose** | **3.8** | 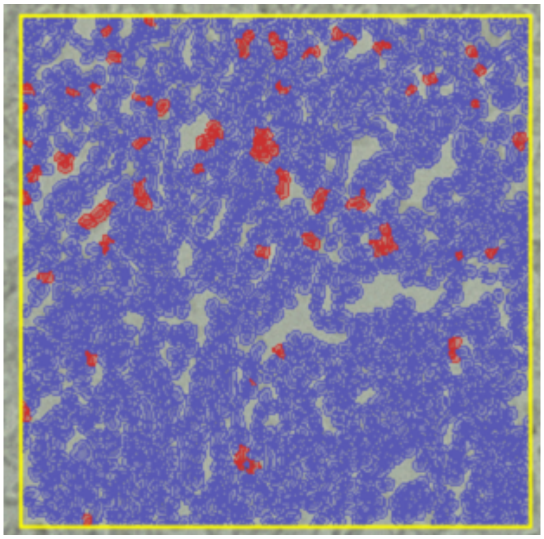 |
| **Gemcitabine & Capecitabine Ping-Pong Dose Modulation** | **7.2** | 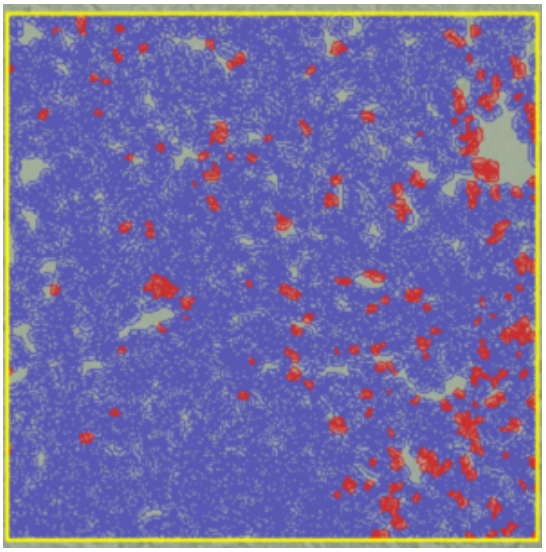 |

| | | |
|---|---|---|
| **Gemcitabine & Capecitabine Ping-Pong Dose Modulation** | **6.9** | 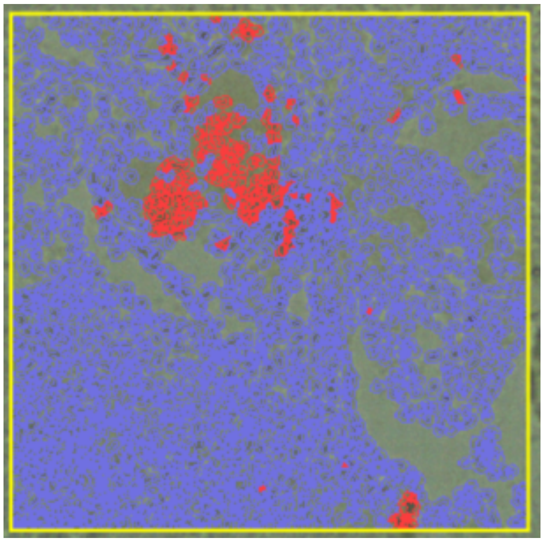 |
| **Gemcitabine & Capecitabine Ping-Pong Dose Modulation** | **4.5** | 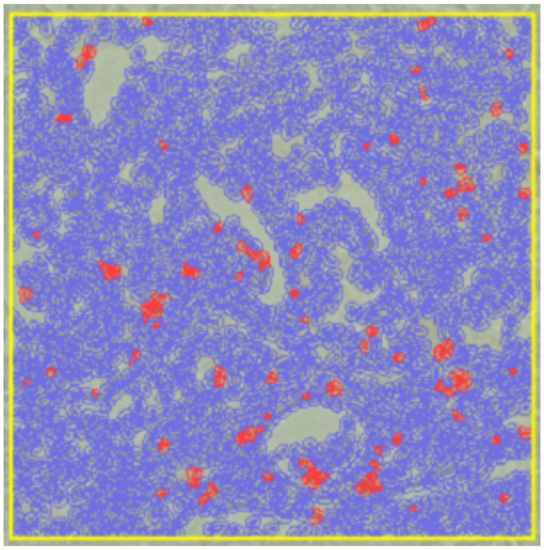 |

| | | |
|---|---|---|
| **Gemcitabine & Capecitabine Ping-Pong Intermittent** | **6.26** | 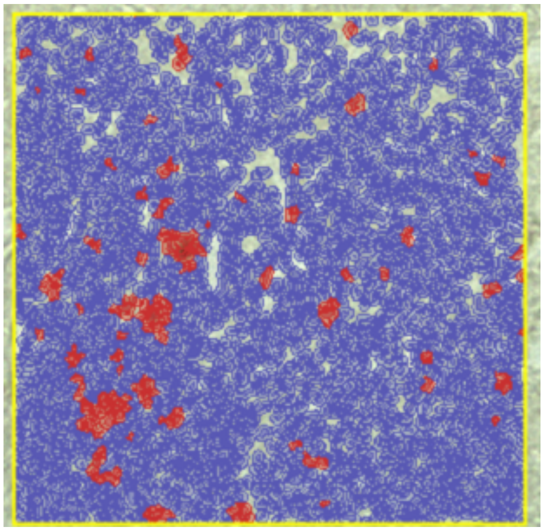 |
| **Gemcitabine & Capecitabine Ping-Pong Intermittent** | **1.7** | 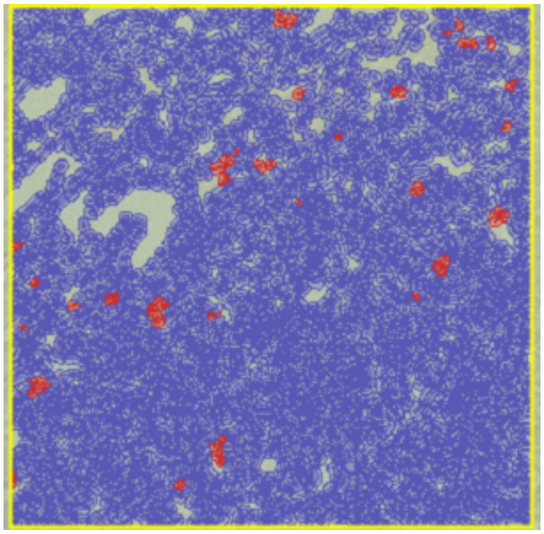 |

| | | |
|---|---|---|
| **Gemcitabine & Capecitabine Ping-Pong Intermittent** | **0.8** | 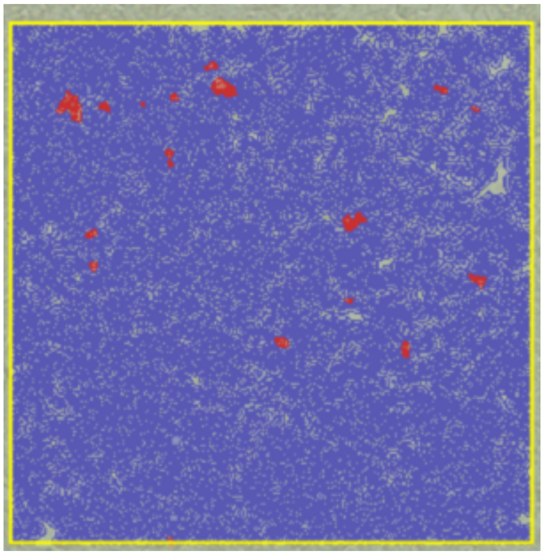 |
| **Gemcitabine & Capecitabine Ping-Pong Intermittent** | **8.06** | 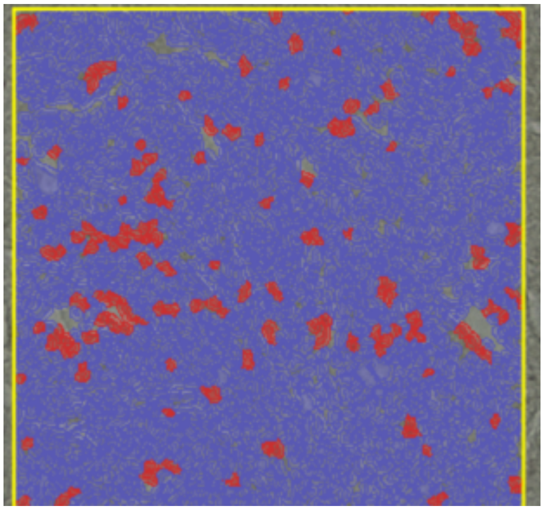 |

| | | |
|---|---|---|
| **Gemcitabine & Capecitabine Ping-Pong Intermittent** | **10.5** | 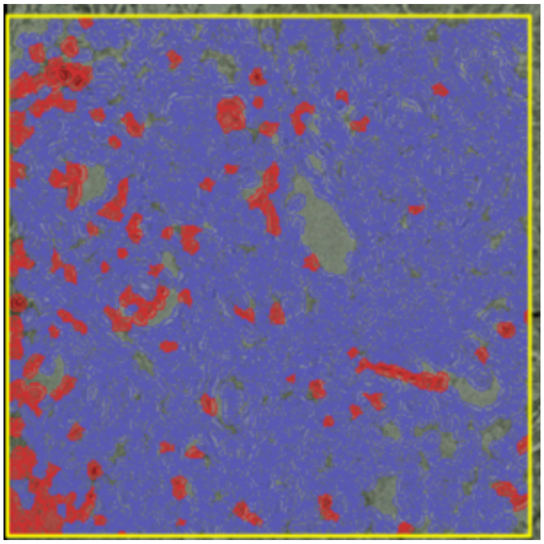 |
| **Gemcitabine & Capecitabine Ping-Pong Intermittent** | **26.5** | 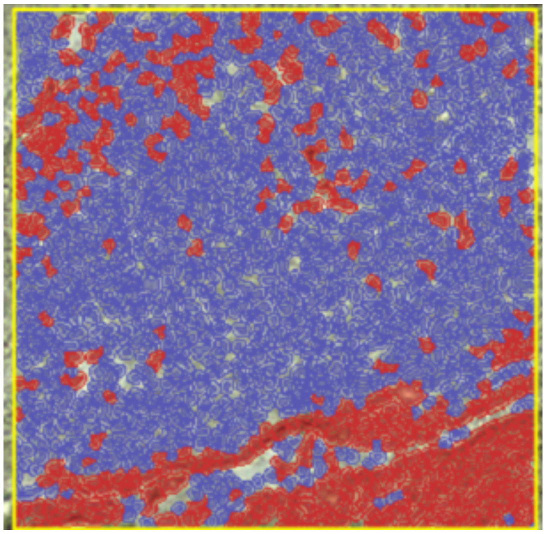 |

| | | |
|---|---|---|
| **Gemcitabine & Capecitabine Tandem Dose Modulation** | **4.6** | 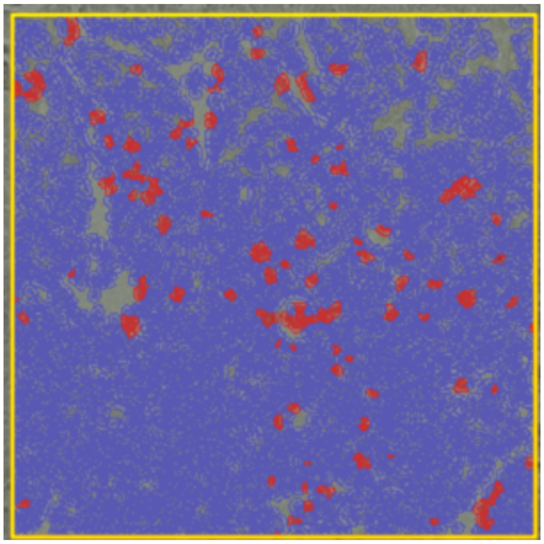 |
| **Gemcitabine & Capecitabine Tandem Dose Modulation** | **1.7** | 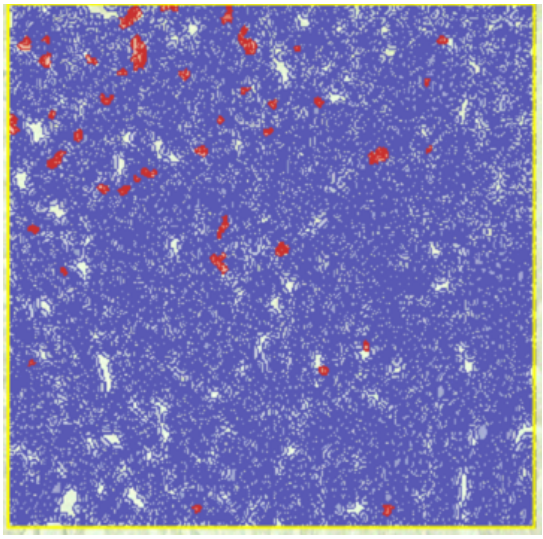 |

| | | |
|---|---|---|
| **Gemcitabine & Capecitabine Tandem Dose Modulation** | **39.8** | 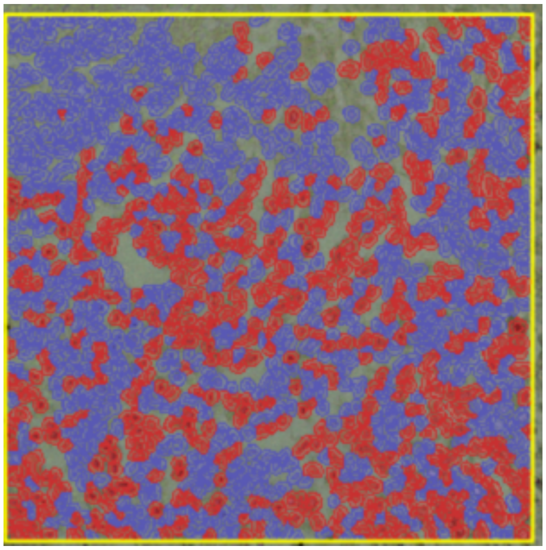 |
| **Gemcitabine & Capecitabine Tandem Dose Modulation** | **8.49** | 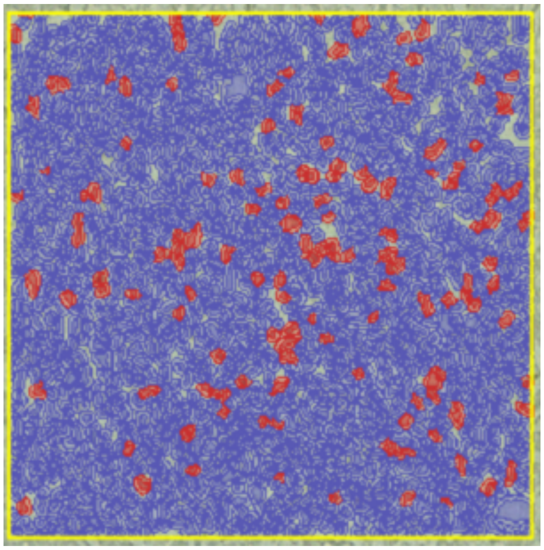 |

| | | |
|---|---|---|
| **Gemcitabine & Capecitabine Tandem Intermittent** | **7.9** | 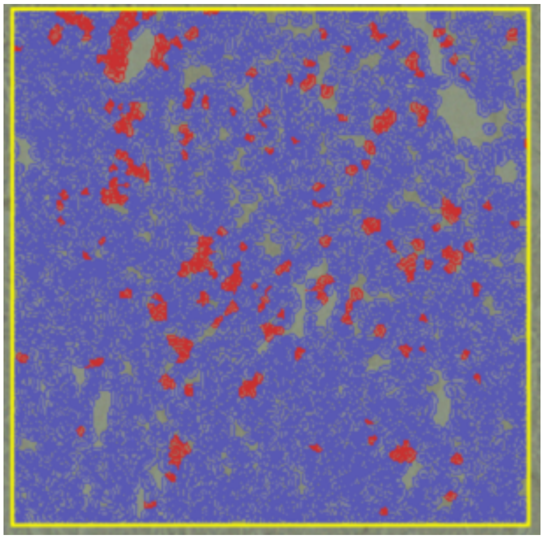 |
| **Gemcitabine & Capecitabine Tandem Intermittent** | **4.96** | 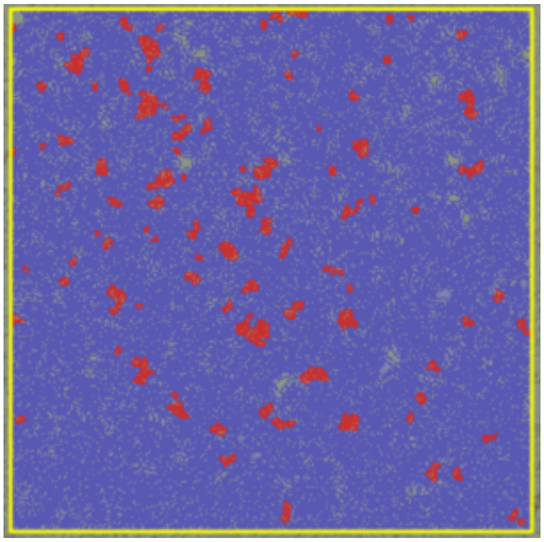 |

| | | |
|---|---|---|
| **Gemcitabine & Capecitabine Tandem Intermittent** | **13.2** | 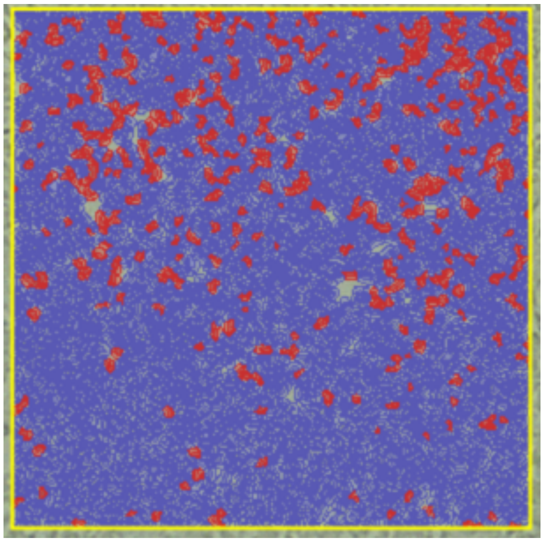 |
| **Gemcitabine & Capecitabine Tandem Intermittent** | **45.9** | 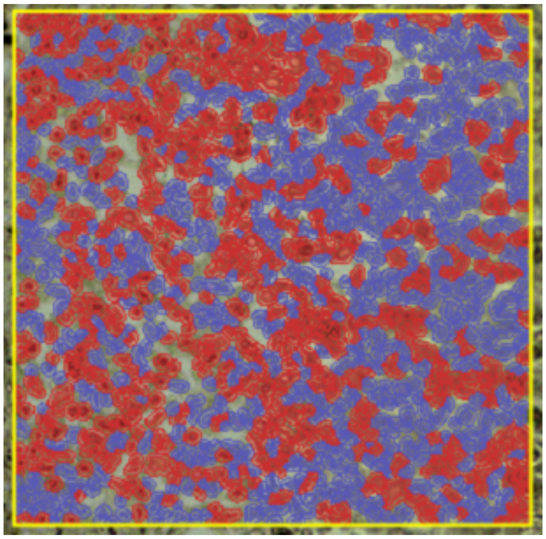 |

**Supplementary Methods**

**Data S1:**

**Characteristics of the drugs used**

Capecitabine is a prodrug that is metabolized into fluorouracil, which blocks thymidine synthesis, thereby blocking DNA synthesis. It is also incorporated into DNA and RNA as a false nucleotide (*38*). Gemcitabine is phosphorylated into dFdCTP which is incorporated into DNA in place of cytosine and blocks further DNA synthesis (*39*).

**Histological Analysis**

We processed the tumors by soaking them in PAXgene tissue fix containers for 2 to 48 hours depending on tissue size. After that, the PAXgene tissue fixative was removed and replaced with PAXgene tissue stabilizer within the same container. The samples were embedded in a paraffin block for hematoxylin and eosin (H&E) staining and immunohistochemistry (IHC). Tumor tissue was sectioned in 6 um slices using Leica Ultracut-R Microtome. Slides were deparaffinized in Epredia™ Signature Series™ Clear-Rite 3™ and rehydrated in an ethanol series. Hematoxylin staining was performed for 1 minute followed by 10 minutes of washing in tap water, 3 dips in 0.1% Acetic acid, and 3 dips in lithium carbonate. Next, we stained the samples in eosin for 1 minute followed by washing them in $DIH_2O$, and then the slides were passed through several alcohol changes followed by rinsing for 3 minutes with Clear-Rite. In the end, we applied a DPX mountant (Sigma-Aldrich) to preserve the stain and cover the slice with coverslips. The slices were imaged at the ASU core facilities using EVOS® FL Auto (The EVOS inverted microscope system, Life Technology).

**Immunohistochemistry**

We performed IHC using the following antibodies: Ki67 is used as a marker of proliferation, to assess the proliferative activity of the tumor and to determine tumor response to our treatment protocols (Ki-67 Recombinant Rabbit Monoclonal Antibody (SP6), invitrogen#MA5-14520), and caspase 3 (Cleaved Caspase-3 (Asp175) Antibody #9661, Cell Signaling) which is an important component of apoptosis.

Caspase-3 is used as a reliable and sensitive method to determine and quantify the apoptosis in our histological samples (*40*) and can be used to detect drug resistance in the form of lower expression of caspase-3 (*41*). We calculated the apoptosis index as a percentage of the apoptotic cells out of the total number of cells examined (Table S3).The Ki-67 is a nuclear antigen that was used as a cell proliferation marker to measure the proliferation rate of tumors. Ki-67 is present in active phases of the cell cycle (*42*). We calculated the proliferative activity of the tumor as a percentage of tumor cells positive for Ki-67 out of the total number of cells examined which correlates to the S-phase fraction (Table S3).

EVOS microscope was used as an imaging system to analyze the morphological association in this breast cancer model. Tumor tissue sections (6 um) slices were deparaffinized in Epredia™ Signature Series™ Clear-Rite 3™ and rehydrated in an ethanol series. Then we blocked the samples with bovine serum albumin (BSA) blocking solution for 1 hour. Thereafter, primary antibodies were diluted in antibody dilution buffer (PBS, Tween20, and BSA, e.g for 50 ml dilution buffer, 48 ml PBS + 2 ML 20% Tween20 + 0.45 g BSA) and were incubated on slides overnight in a humidified box at 4 °C followed by washing with PBS for 10 minutes. Then, we applied secondary antibodies (diluted in dilution buffer), Donkey anti-Rabbit HRP,

(Cat#31458, Invitrogen) for 1 hour at room temperature. Next, DAB staining was done for 10 minutes as a chromogen followed by staining in hematoxylin for 1 minute. Then slides were washed in tap water for 10 minutes followed by dehydration in 95 % and 100% alcohol for 1 minute each and then mounted at the end. The tissue sections were imaged at the ASU core facilities using EVOS® FL Auto (The EVOS inverted microscope system, Life Technology). All the stained images with Ki-67 and caspase-3 antibodies were analyzed using Qupath software to determine the Ki-67 and caspase-3 positive index and the data were analyzed by GraphPad Prism software. The integrated density value of immunostain for fixed tissues was calculated using Qupath. Images were stored as TIF files with 10x resolution and were imported to Qupath as "Heme/DAB brightfield" images. The same size annotation was used for all the samples and in the setting, we used 0.01 for the threshold and Nucleus DAB OD means with a single threshold was set on the images.

**Data S2: Protocols for treatment with a single cytotoxic drug and two cytotoxic drugs described in Supplementary method:**

***MTD***: Drug 1 was administered at a maximum tolerated dose (MTD) once every 24 hours for the entire duration of the simulation.

***Dose Modulation***: Treatment started at MTD with Drug 1, and the dosage of the drug was adjusted according to the dose modulation adaptive therapy protocol, parameterized by Delta Tumor, and Delta Dose. This treatment protocol was equivalent to the standard dose modulation adaptive therapy protocol (AT-1) from previous experiments (*2*).

***Intermittent***: Treatment started at 100% of the MTD using Drug 1, drug being administered once every 24 hours. Treatment was stopped when a shrinkage in tumor burden by at least 50% relative to the tumor burden at which treatment was initiated is detected, and treatment was restarted when the tumor burden equaled or exceeded 100% of the value at which treatment was initiated.

***MTD*****:** This is identical to Standard Treatment (ST) with both Drug 1 and Drug 2 being administered at maximum tolerated dose (MTD) in a cocktail formulation once every 24 hours for the entire duration of the simulation.

***Tandem Dose Modulation*****:** This is identical to DM Cocktail Tandem treatment starting at MTD for both the drugs, and dosages of both drugs being adjusted simultaneously as parameterized by Delta Dose and Delta Tumor.

***Ping-pong Dose Modulation*****:** This is identical to DM Ping-Pong Alternate Every Cycle with treatment starting with Drug 1 at MTD, which is followed by Drug 2 at MTD during the subsequent cycle. Drugs are switched every cycle and the dosages, as parameterized by Delta Dose are adjusted based on the tumor's response to the same drug, as parameterized by Delta Tumor the last time it was administered.

***Tandem Intermittent*****:** This is similar to FD Intermittent except that both Drug 1 and Drug 2 are administered as a cocktail formulation at 100% of the MTD for each drug, not at 75% of the MTD that we previously explored. Therefore, treatment starts as a cocktail formulation of both the drugs at 100% of the MTD, with the drugs being administered once every 24 hours. Treatment is stopped when the tumor burden falls by at least 50% relative to the tumor burden at which treatment was initiated, and treatment is restarted when the tumor burden equaled or exceeded 100% of the value at which treatment was initiated.

***Ping-pong Intermittent*****:** This is similar to Tandem Intermittent, except that the drugs are switched every time the tumor climbs back to 100% or more of the baseline tumor burden at which treatment was initiated.
